# Flow in fetoplacental microvessels in vitro enhances perfusion, barrier function, and matrix stability

**DOI:** 10.1101/2023.07.19.549736

**Authors:** Marta Cherubini, Scott Erickson, Prasanna Padmanaban, Per Haberkant, Frank Stein, Violeta Beltran-Sastre, Kristina Haase

## Abstract

Proper placental vascularization is vital for pregnancy outcomes, but assessing it with animal models and human explants has limitations. Here, we present a 3D *in vitro* model of human placenta terminal villi that includes fetal mesenchyme and vascular endothelium. By co-culturing HUVEC, placental fibroblasts, and pericytes in a macro-fluidic chip with a flow reservoir, we generate fully perfusable fetal microvessels. Pressure-driven flow is crucial for the growth and remodeling of these microvessels, resulting in early formation of interconnected placental vascular networks and maintained viability. Computational fluid dynamics simulations predict shear forces, which increase microtissue stiffness, decrease diffusivity and enhance barrier function as shear stress rises. Mass-spec analysis reveals the deposition of numerous extracellular proteins, with flow notably enhancing the expression of matrix stability regulators, proteins associated with actin dynamics, and cytoskeleton organization. Our model provides a powerful tool for deducing complex *in vivo* parameters, such as shear stress on developing vascularized placental tissue, and holds promise for unraveling gestational disorders related to the vasculature.

## Introduction

Human placentae are highly vascularized organs that undergo constant vascular growth and remodeling to ensure effective exchange between maternal blood and the fetal circulatory system [1]. The majority of the essential exchange of solutes (oxygen, nutrients, hormones, antibodies) takes place across a thin layer of epithelial cells of the terminal villous trees, which contains a network of fetoplacental capillaries. This dense capillary network develops from the elongation and ramification of pre-existing blood vessels formed by vasculogenesis (new vessel formation) by approximately 6 weeks of gestation. From 25 weeks post-conception until term, villous vascular growth switches from branching angiogenesis to non-branching angiogenesis generating coiled capillary structures that reside within the extremities of the fetoplacental vascular trees [2].

Formation of a proper vascular tree in the early placenta is crucial to ensure optimal blood volume loading, preventing placental insufficiency and adverse fetal outcomes. Structural villous microvascular network abnormalities are common features of gestational disorders associated with perinatal morbidity and mortality such as in pre-eclampsia and fetal growth restriction [3, 4]. To determine the etiology of these alterations, it is crucial to understand the mechanisms underlying fetoplacental vasculogenesis and regulation.

Studying fetoplacental vascular development is challenging, mainly due to ethical considerations and inaccessibility of the tissue, particularly in early pregnancy. Current knowledge on villous vascular network development and function comes from observations of human placenta explants obtained at different stages of gestation [5, 6], ultrasound examinations [7] and *in silico* modelling [8, 9]. However, these methods cannot provide a direct measurement of the physical and chemical cues occurring during placental vasculogenesis and angiogenesis, such as mechanical shear stress and associated mechanotransduction. Developing vessels *in vivo* are exposed to constant mechanical stimuli including fluid shear stress induced by interstitial flow (IF) and intraluminal blood flow [10]. Several studies have documented the effects of flow-induced shear stress on vascular morphogenesis, demonstrating flow-induced regulation of angiogenic sprouting and lumen formation by endothelial cell migration, alignment, and apical deformation [11–14]. Hemodynamic forces trigger vessel remodeling through cellular rearrangement, and regulate vascular permeability via modulation of endothelial adherens and tight junctions [15–18]. The role of hemodynamics during pregnancy remains poorly understood since *in vivo* experimentation has mainly focused on umbilical circulation [19], lacking proper characterization of the placental micro-hemodynamics occurring at the fetoplacental interface. Although the placenta at term constitutes a valid tool for *ex vivo* perfusion studies [20], an understanding of early placental vascular circulations is limited, requiring the employment of biomimetic models.

Tissue-specific 3D microvascular networks have recently contributed to our understanding of flow-induced vascular growth and remodeling, by overcoming the limitations of simpler monolayer systems [14, 16, 21, 22]. For instance, interstitial flow has been shown to enhance vessel formation, function and longevity in brain-specific microvasculature comprised of endothelial cells, pericytes, and astrocytes [23]. Despite the development of novel *in vitro* placental models (reviewed in [24]), at present, no models have been used to investigate the effect of flow-associated placental vascular development. Previously, our 3D model of terminal villi microvasculature was established using a triculture of stromal and endothelial cells [25]. This model demonstrated that placental pericytes contribute to growth restriction, which was shown to be largely dependent on VEGF and angiopoietin/Tie2 signaling [25]. Fibroblasts, on the other hand, contributed to increased vasculogenesis; however, a limitation of our former model was the use of non-specific fibroblasts. To build on our former study, placental fibroblasts are now included in a completely placental-derived human 3D vascular model. Our results indicate that growth and remodeling of fetal microvessels, as well as endothelial barrier function (permeability to solutes) and extravascular matrix properties (diffusivity, stiffness, and matrix proteins), are strongly regulated by increasing shear stress. Computational fluid dynamics predicts luminal and interstitial shear stresses, which are highly heterogenous within a single microtissue. Flow-conditioned fetal microvessels exhibit pro-inflammatory profiles and early connection of vessel networks that remain perfusable for weeks in culture.

Overall, fluid dynamics strongly regulate the development of fetoplacental vessels, suggesting that poor or restrictive flow conditions can negatively impact vasculogenesis. Our findings serve as a basis for further research into the mechanisms of placental vascular defects in pregnancy-related disorders related to flow insufficiency.

## Material and methods

### Cell culture

Human umbilical vein endothelial cells (HUVEC) were purchased from Lonza and cultured in endothelial media (VascuLife, Lifeline cell systems) on T75 or T150 flasks coated with 50 μg/ml rat tail collagen I (Roche). HUVEC were transduced to stably express cytoplasmic RFP (LentiBrite RFP Control Lentiviral Biosensor, Millipore Sigma Aldrich), and were used between passages 6 and 9. Primary human placental fibroblasts were purchased from ATCC, cultured in fibroblast media on uncoated T75 flasks (FibroLife, Lifeline cell systems) and used between passages 4 and 8. Unlabeled and GFP-labeled primary human placental microvascular pericytes were acquired from Angioproteomie, cultured in pericyte growth medium according to manufacturer’s protocols and used between passage 6 and 9. All cells were cultured under normal conditions at 37 °C and 5% CO_2_ in a standard incubator, dissociations were carried out using TrypLE Express (Gibco), and media was completely refreshed every other day.

### Device fabrication

Devices were fabricated as previously described [25]. Briefly, PDMS (SYLGARD^TM^ 184 Silicone Elastomer Kit, Dow) was mixed at a 10:1 elastomer to cross-linker ratio according to the manufacturer’s protocols, degassed, and poured onto a mold. Following further degassing, PDMS was placed in a 60° C oven overnight and single devices were cut, punched using 1– and 2-mm diameter biopsy punches for the gel and media ports, respectively, and air-plasma bonded (Harrick systems) to clean glass slides. Assembled devices were then incubated at 60° C overnight to restore the native hydrophobic state. All devices were sterilized under ultraviolet light for at least 30 min prior to cell seeding.

### Device seeding and microvessels formation under interstitial flow

Fibrinogen derived from bovine plasma (Sigma) was reconstituted in phosphate-buffered saline (PBS) to a working concentration of 7 mg/ml before use. Thrombin (Sigma) stock solution (100 U/ml in 0.1% w/v bovine serum albumin solution) was diluted to a 4 U/ml working solution in cold VascuLife medium. Endothelial cells and stromal cells were cultured until near-confluence prior to detachment and re-suspended in thrombin, separately, to concentrations of 24 million endothelial cells/ml and 2.4 million stromal cells/ml (a combination of fibroblasts and pericytes). Cell suspensions were mixed 1:1 by volume and then combined with fibrinogen solution to produce a final concentration of 6 million endothelial cells/ml and 0.6 million stromal cells/ml, in a 10:1 ratio within fibrin (3.5 mg/ml). The cell-gel mixture was injected into the device central channel and allowed to polymerize for 15–20 min at 37 °C in a humidified chamber. VascuLife media was added to each media channel (150 uL total) and devices were initially cultured under static conditions. An intermittent interstitial flow (IF) across the hydrogel channel was generated 48 hours after seeding by adding 3D-printed tailored media reservoirs (Form3 printer, Formlabs). Equal total volumes of media were differentially added to the reservoirs compartments to generate three hydrostatic pressure gradients (ΔP): 0, 3– and 7-mm H_2_O. The pressure gradients were re-stablished with fresh media every day for 5 days following seeding. From day 8 onwards, with the vessels fully perfusable, hydrostatic pressure gradients were restored every other day.

### Imaging and vessel morphology quantification

All images were acquired on a Stellaris 8 confocal microscope using LAS X software (Leica). Confocal z-stacks were obtained for all time-points and used to quantify the morphology of the microvascular networks cultured under different ΔP, as previously described [25]. Briefly, a custom macro (ImageJ, NIH) was generated to process the images as follows: projection of maximum intensity of the RFP channel in the z-direction, Gaussian filter smoothing, removal of outliers and conversion to binary images. The analyze particles and 2D skeletonize built-in ImageJ plugins were finally performed to determine branch density, network connectivity and effective diameter (vessel area/total length).

### Permeability measurements

Microvessels cultured under different flow conditions were perfused at day 7 or 14 with 70 kDa FITC-labelled dextran (Merck) by the application of a hydrostatic pressure drop across the central gel channel. Briefly, media was removed and replaced by adding 40 ul of fluorescent perfusate solution (FITC 0.1 mg/ml) into one channel. After perfusion into and through the microvessels, convective flow was stopped by applying an equal volume of dextran solution in the opposite channel. Following stabilization (∼2–3 min), time-lapse confocal images were captured (3 × 5 min intervals), from which permeability measurements were made, as reported previously [25].

### Cytokine release

Supernatants were collected from 3 device reservoirs per ΔP condition of two or three independent experiments at days 3, 5 and 7 and were frozen until use. Concentration of VEGF, Ang-2 and MCP-1 were measured from individual samples following the manufacturer’s protocols of respective Quantikine ELISA Kits (R&D systems, DVE00, DANG20 and DCP00). Cytokine expression profiles of 2D cultures were assessed using a human angiogenesis antibody array (Abcam). Supernatants from lung fibroblasts, placental fibroblasts and pericytes were collected 48 hours after seeding and processed according to the manufactureŕs instructions. Cytokine profiles on the array membranes were detected by chemiluminescence using the Fusion FX Spectra (Vilber, France). The relative (semi-quantitative) expression of intensity was normalized to the positive controls.

### Fluid dynamics characterization by bead-tracking

Evaluation of fluid velocity of static and flow-conditioned vessels was assessed at day 7 through perfusion of 2.0 µm fluorescent beads (Fluorescent blue latex beads, Sigma Aldrich) into the microvascular networks using reservoirs with applied pressure gradients of 5 mm H_2_O. Time-lapse images (1024×1024 pixels at a resolution of 0.64 μm/pixel with a 0.06 sec frame rate) were acquired for 25 sec using a fluorescent microscope (Thunder Imager – DMi8 microscope, Leica). Tracking of each individual bead (distance over time) was done using the particle tracking plugin – *TrackMate* (NIH ImageJ) [26]. First, individual beads are detected based on size through the “LoG detector” method with an “Estimated blob diameter” of 5 µm, and an intensity threshold of 2 AU. Then, the “simple LAP Tracker” method with “Linking max distance” of 15 µm, “Gap-closing max distance” of 15 µm, and “Gap-closing max frame gap” of 2 frames, is used to identify the same object over time. Upon application of a filter for “Track displacement” to easily remove non-moving objects, the function “Analysis” is used to obtain the mean speeds of the beads.

### Computational fluid dynamics simulation

In order to estimate the range of fluid velocities and shear stresses generated by the application of pressure-induced flows within microvessels and surrounding gel domains, binary masks were obtained from confocal microscopy images and converted to drawing exchange format (dxf) in Adobe Illustrator 2021. These .dxf files were imported into COMSOL multiphysics software (Version 6.0). Using the laminar flow physics module with porous domain enabled physics setting, the spatial profile of luminal and interstitial fluid velocities and shear stresses were computed for 70 Pa applied pressure (a pressure drop, ΔP, of 7 mm H_2_O) between the inlet and outlet of four microfluidic chips containing placental microvessels. A computationally efficient physics-controlled extra fine mesh was applied to the model, resulting in an overall computational time of ∼2 hours.

### Diffusivity measurements and initial fluid velocity tracking

Diffusivity was assessed in the extravascular space (adjacent but outside of vessels observable in images) for each flow condition using FRAP measurements on a Stellaris 8 confocal microscope (Leica). Devices were perfused with 70 kDa FITC labelled dextran and incubated at 37 °C for 12 hours to allow total diffusion throughout the gel matrix (extravascular space). Small regions (30 µm Ø) within the matrix were bleached and, immediately after photobleaching, time-lapse images were recorded every 0.4 sec to capture the fluorescence recovery. Analysis of diffusivity was performed using the *Matlab frap_analysis* plugin [27], as previously described [28]. Similarly, we determine the initial IF velocity (day 2 after seeding), using FRAP as done previously [16]. Briefly, pressure gradients were generated across the gel and cell containing devices by adding FITC dextran to the corresponding reservoir media volumes. Following dextran saturation, a region of 30 µm Ø was bleached and consecutive time-lapse images were collected to observe the fluorescence recovery following the direction of fluid flow. The IF velocity was estimated using the *Matlab frap_analysis* plugin to track the movement of the centroid of the bleached region.

### Microtissue stiffness assessment

Mechanical resistance of fetal placental vascularized microtissues cultured under different flow conditions was measured on day 7 using a Chiaro Nanoindenter (Optics 11, Amsterdam, Netherlands), as previously described [28]. Briefly, gels were cut and extracted from the devices, placed in a petri dish and fully submerged in VascuLife media. Nanoindentation measurements were performed using a spherical probe tip with a radius of 25 μm and a stiffness of 0.027 N/M. After proper calibration of the probe, 12 μm depth indentations were applied to the gels. The effective Young’s modulus values were derived from load-indentation curves by fitting to the standard Hertz model and assuming a Poisson’s ratio of 0.5, using the manufacturer’s data analysis plug-in. An average of ∼80 measurements were performed per condition, for two independent experiments.

### Flow cytometry

Characterization of stromal cell population was performed at day 3 and 7 after seeding by flow cytometry. To extract the cells from the devices, the gels were resected and digested in a solution of Accutase (Gibco) and 50 FU/ml Nattokinase (Japan Bioscience Ltd) for 15-20 min at 37° C. An average of 6 gels was pooled for each time point. Single cells were then stained with the pericyte marker CD140b PerCP-Cy™5.5 (BD Bioscience) for 2 hours at 4° C, washed with PBS, analyzed on a BD LSR II and later processed using FlowJo v.10.8.1 software. The stromal cell population was gated using CD140b^+^ and by exclusion of the RFP^+^ cell cluster (expressed only in HUVEC).

### Immunostaining

On day 7, devices were washed with PBS and fixed with 4% paraformaldehyde for 15 min. Samples were then permeabilized and blocked for non-specific binding with 0.1% v/v Triton X-100, 5% w/v BSA and 1% v/v serum (same source of secondary antibody) in PBS overnight at 4° C on an orbital shaker. Primary antibodies were diluted in 0.5% w/v BSA PBS and added to samples overnight at 4° C on an orbital shaker. Primary antibodies used in this study are S100A4 (FSP-1) (1:100, Abcam), Collagen I (1:200, Abcam), Fibronectin (1:200, Abcam), and Laminin (1:200, Abcam). The following day, samples were washed with 0.1% v/v Triton X-100 PBS and then incubated with the appropriate secondary antibody (1:200, Alexa Fluor 488 or 647, Invitrogen) and DAPI counterstain diluted in PBS overnight at 4° C on an orbital shaker. Samples were washed with PBS and stored at 4° C before imaging. To ensure effective staining and washing of unbound reagents, all washes and incubations steps were performed while applying a pressure gradient across the gel.

### Mass spectrometric characterization of matrix derived proteins

Tryptic digestion of hydrogel/tissue samples for their for mass spectrometry (MS) analysis was performed at day 7 after seeding as described by L.A. Sawicki et al. [29]. Two experiments were conducted, with each experiment having ≥ 2 replicates per static– or flow-conditioned devices. Samples were first decellularized to remove cellular structures and protein contribution. Briefly, extracted gel matrices were washed with PBS and wash buffer (100 mM Na_2_HPO_4_, 2 mM MgCl_2_, 2 mM EGTA) prior incubation to a lysis buffer (8 mM Na_2_HPO_4_, 1% NP-40) for 90 min at 37 °C. Following additional washes (300 mM KCl, 10 mM Na_2_HPO_4_), decellularized gels were degraded with collagenase (Gibco, 50 U/mL in HBSS) for 1 h at 37° C to release all retained proteins. Gels were then dissolved by vigorous pipetting and stored at –80°C prior to lyophilization (Labconco FreeZone 2.5). Lyophilized samples were reconstituted in 25 mM NH₄HCO₃, reduced with DTT and Cysteine residues were alkylated by the addition of iodoacetamide [29]. Proteins were digested over night by the addition of trypsin (Promega) and samples were acidified by using formic acid according to [29]. Digested samples were applied to spin column concentrators (Corning) with a 10 kDa cut off in order to remove trypsin and collagenase. Mixed peptides were subjected to a reverse phase clean-up step (OASIS HLB 96-well µElution Plate, Waters). Peptides were dried and reconstituted in 10 µl of 400 mM Hepes/NaOH, pH 8.5 and reacted for 1 h at room temperature with 80 µg of TMT18plex (Thermo Scientific) dissolved in 4 µl of acetonitrile. Peptides were subjected to a reverse phase clean-up step prior their analysis by LC-MS/MS on an Orbitrap Fusion Lumos mass spectrometer (Thermo Scientific).

To this end, peptides were separated using an Ultimate 3000 nano RSLC system (Dionex) equipped with a trapping cartridge (Precolumn C18 PepMap100, 5 mm, 300 μm i.d., 5 μm, 100 Å) and an analytical column (Acclaim PepMap 100. 75 × 50 cm C18, 3 mm, 100 Å) connected to a nanospray-Flex ion source. The peptides were loaded onto the trap column at 30 µl per min using solvent A (0.1% formic acid) and eluted using a gradient from 2 to 80% Solvent B (0.1% formic acid in acetonitrile) over 2 h at 0.3 µl per min (all solvents were of LC-MS grade). The Orbitrap Fusion Lumos was operated in positive ion mode with a spray voltage of 2.2 kV and capillary temperature of 275° C. Full scan MS spectra with a mass range of 375–1.500 m/z were acquired in profile mode using a resolution of 120.000 with a maximum injection time of 50 ms, AGC operated in standard mode and a RF lens setting of 30%. Fragmentation was triggered for 3 s cycle time for peptide like features with charge states of 2–7 on the MS scan (data-dependent acquisition). Precursors were isolated using the quadrupole with a window of 0.7 m/z and fragmented with a normalized collision energy of 34%. Fragment mass spectra were acquired in profile mode and a resolution of 30,000. Maximum injection time was set to 94 ms and AGC target to custom. The dynamic exclusion was set to 60 s.

Acquired data were analyzed using FragPipe [30] and a Uniprot Homo sapiens database (UP000005640, ID9606 with 20594 entries, October 26^th^ 2022) including common contaminants. The following modifications were considered: Carbamidomethyl (C, fixed), TMT18plex (K, fixed), Acetyl (N-term, variable), Oxidation (M, variable) and TMT18plex (N-term, variable). The mass error tolerance for full scan MS spectra was set to 10 ppm and for MS/MS spectra to 0.02 Da. A maximum of 2 missed cleavages were allowed. A minimum of 2 unique peptides with a peptide length of at least seven amino acids and a false discovery rate below 0.01 were required on the peptide and protein level [31].

### MS data processing

The raw output files of FragPipe (protein.tsv – files, [30]) were processed using the R programming language (ISBN 3-900051-07-0). Contaminants were filtered out and only proteins that were quantified with at least two unique peptides were considered for the analysis. 333 proteins passed the quality control filters. Log2 transformed raw TMT reporter ion intensities were first cleaned for batch effects using the ‘removeBatchEffects’ function of the limma package [32] and further normalized using the vsn package (variance stabilization normalization, [33]). Proteins were tested for differential expression using the limma package. The replicate information was added as a factor in the design matrix given as an argument to the ‘lmFit’ function of limma. A protein was annotated as a hit with a false discovery rate (fdr) smaller than 5 % and a fold-change of at least 30 % and as a candidate with a fdr below 5 % with no fold-change threshold. Hit and candidate proteins were clustered into 2 clusters (method kmeans) based on the Euclidean distance between normalized TMT intensities divided by the 0 mm H_2_O data point.

### Statistics

Statistical significance was analyzed using OriginPro v.9.85 performing one-way ANOVA followed by the post-hoc Tukey’s test for multiple comparison. Differences were considered statistically significant with p <0.05. Data shown here are from at least 2 independent experiments with n ≥ 3 devices, with at least 3 measurements per device, unless otherwise specified.

## Results

### 1. Interstitial flow enables placental microvasculature network formation and function

We previously presented a 3D *in vitro* model of placental terminal villi microvessels capable of recapitulating several aspects of placental vasculopathies [25]. Now, to better approximate the physiological cellular composition of placental fetal tissue, we sourced primary placental fibroblasts that were integrated together with endothelial cells (HUVEC) and placental pericytes to generate microvascular tissues on-chip. Cells were cultured with an endothelial to stromal cell ratio of 10:1 in a 3.5 mg/ml fibrin gel, as opposed to the 5:1 ratio and 3 mg/ml used previously [25], within a single-gel channel device (Fig. 1A). These differences in culture are in-part attributed to the different secretome of stromal cells isolated from placental and lung tissues, which show dissimilarities in angiogenic profile (Fig. S1). In particular, inflammatory molecules including PDGF-BB and MCP-1 are increased for placental fibroblasts in contrast to lung fibroblasts. The formation of connected microvascular networks (with this new tri-culture) occurs over approximately 1 week (Fig. 1B); however, networks are fully perfusable only when an interstitial flow (IF) is applied across the hydrogel through the addition of a 3D printed media reservoir (shown in Fig. 1A top right). Placental microvessels cultured under static conditions self-assemble by day 4; however, they were not perfusable, showing loss of connectivity and vessel pruning by day 7 (Fig. S2).

**Figure. 1.**
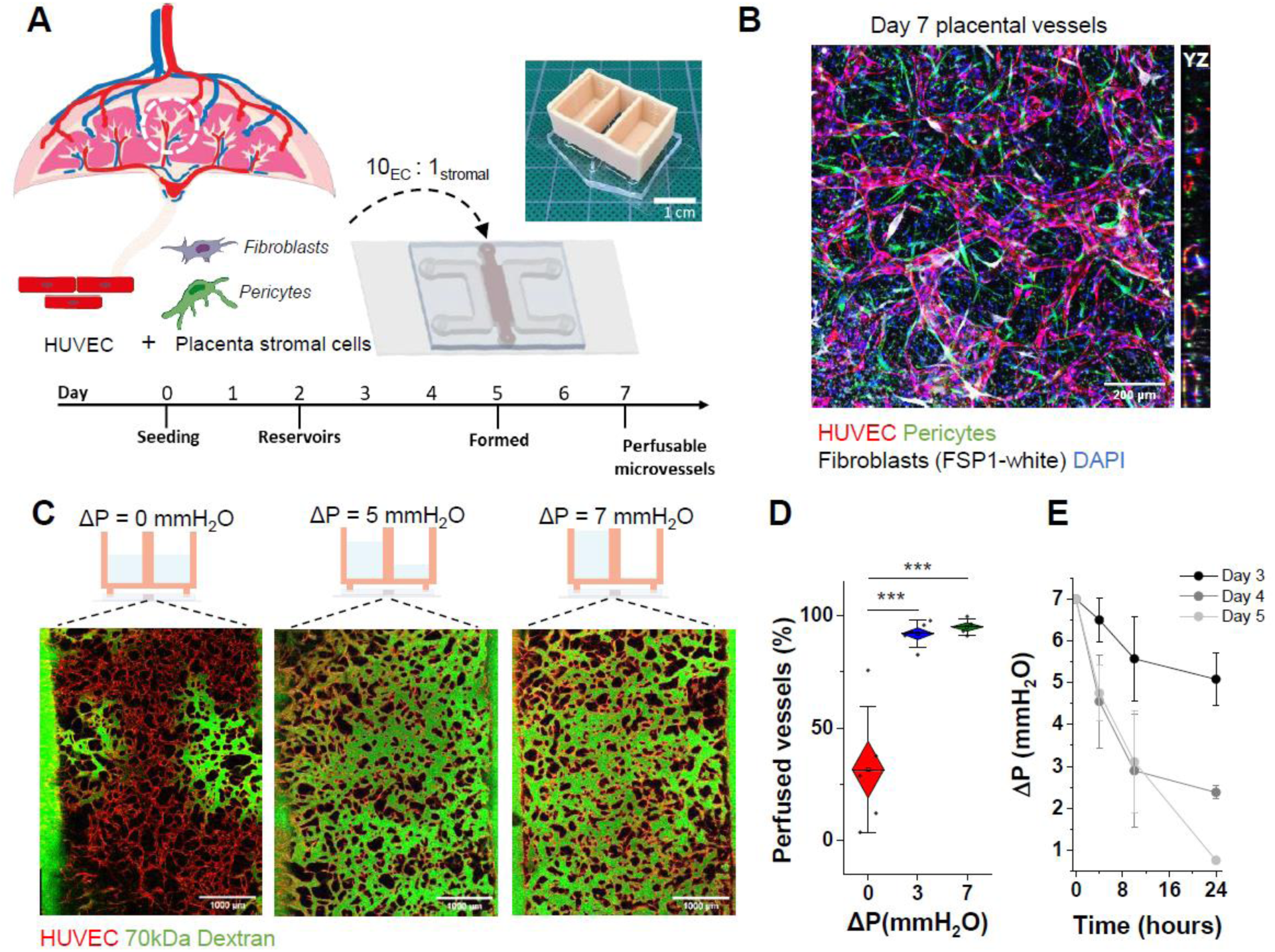
Interstitial flow promotes network perfusion in placental microvessels. A) Schematic demonstrating the culture timeline and protocol. B) Immunofluorescence image of tri-culture placental vessels with orthogonal view showing open vascular lumen. C) Interstitial flow conditions (depicted graphically) promote complete vascular bed perfusion. Perfusion is shown by FITC labelled dextran introduced into the RFP-labelled vessels at day 7. Scale bar is 1000 µm. D) Percentage of perfused vessels is quantified across static and interstitial flow conditioned vessels at day 7. Shown is box plots with outer edge as standard error and bars with standard deviation. Significance is measured by One-way ANOVA and indicated *** < 0.001 for Tukey means comparison test. E) Pressure drop measured over time across different days in culture. Shown is mean and standard deviation of 3 separate experiments with ≥6 devices.

Pressures gradients (ΔP) representative of static (0 mm H_2_O), low (3 mm H_2_O or ∼30 Pa) and high flow (7 mm H_2_O or ∼70 Pa) culture conditions were applied to microvessels (Fig. 1C), with mean interstitial fluid flow velocities measured as 0.13 ± 0.06 µm/s, 0.30 ± 0.12 µm/s and 1.23 ± 0.32 µm/s, respectively at day 2 (Fig. S3). On day 7, perfusion with FITC-dextran shows complete vessel connectivity in devices cultured under flow, whereas partial perfusion was observed in its absence (Fig. 1C), quantified as perfused vessel area (Fig. 1D).

Vessels form over several days in culture to produce open perfusable lumen (of ∼40 µm in diameter at day 7), hence there is a shift from interstitial to luminal flow. This transition was measured by the fluid volumetric drop (every ≈4 hours). The pressure gradient decays slowly at earlier timepoints, but demonstrates a quick drop over 24 hours by day 5. This suggests the presence of open-lumen and a switch from interstitial to primarily intraluminal flow (Fig. 1E).

### 2. Interstitial flow promotes early placental vasculogenesis

Placental microvessels become increasingly perfusable when subjected to flow, thus morphologic and barrier properties were assessed. Development of the vessels was monitored at specific timepoints by confocal microscopy under static, low– and high-flow conditions (Fig. 2A). In early culture (day 3), if-conditioned vessels showed increased network connectivity (Fig. 2B), reduced branch density (Fig. 2C) and larger vessel diameters (Fig. 2D), which were significant for high-flow conditions. At day 7, connectivity remained unchanged between flow and static conditions, although values of connectivity were increased over time for static cultured vessels. At this later time point, branch density was reduced in all cases compared to day 3 under the same conditions (Fig. 2C). Although diameter continues to increase over time (from day 3 to 7) across both conditions, flow-conditioned microvessels are significantly larger in diameter compared to static cultured vessels, for both low and high-flow (Fig. 2D).

**Figure. 2.**
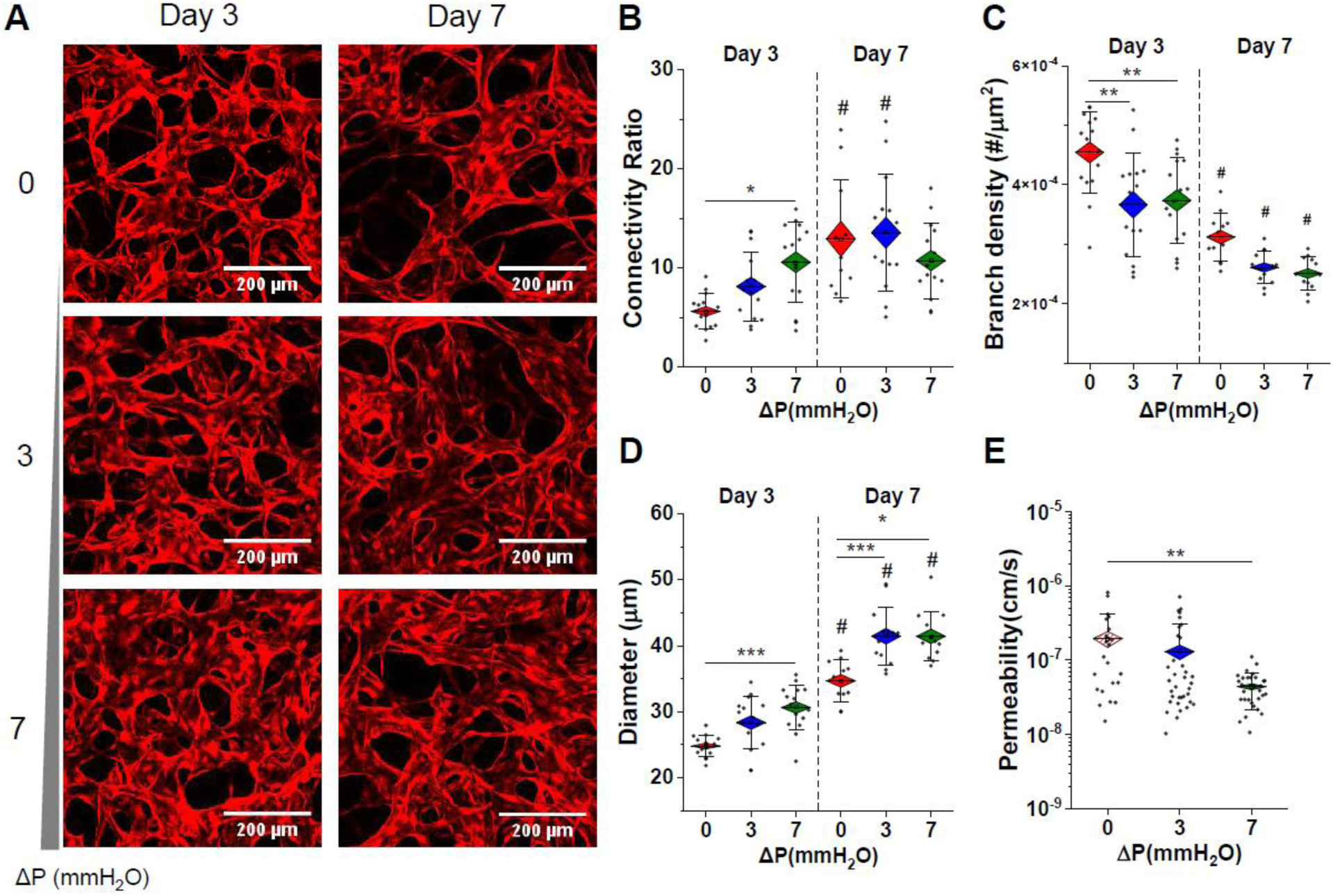
Interstitial flow promotes vessel connectivity and barrier function. A) Confocal images of microvessels (HUVEC with cytoplasmic RFP) at day 3 and 7 of culture under static and flow conditions. Morphology is measured by segmentation of images of RFP-cytoplasmic labeled HUVEC. Morphologic parameters were measured across days 3 and 7 for B-D. B) Connectivity is quantified as a ratio of vessel junctions to endpoints. C) Branch density is the number of branches per area. D) Effective diameter is measured as vascular area coverage over length. E) Permeability to 70kDa dextran is shown for static and flow conditions at day 7. Hashed lines for the static condition indicate that very few vessels were perfused and thus measured. Significance is measured by One-way ANOVA and indicated by *<0.05, **<0.01, and *** < 0.001 for Tukey means comparison test and # represents significance across days.

Next, endothelial barrier function was examined by perfusion on day 7 with 70 kDa FITC-dextran. Vessels cultured under high flow conditions, in comparison to static culture, resulted in significantly decreased permeability values (increased barrier function) (Fig. 2E), indicating that an intermittently applied pressure gradient is sufficient to enhance vascular stability.

### 3. Interstitial flow enhances inflammatory signaling in developing placental vessels

Vessel formation and remodeling is intimately linked to inflammation and growth factor signaling, yet little is known about signaling in human fetoplacental vasculogenesis. Thus, cytokine expression was examined from static and flow-conditioned placental microvessels. Inflammatory cytokines, including interleukin 8 (IL8) and monocyte chemoattractant protein-1 (MCP-1), and pro-angiogenic factors, including vascular endothelial growth factor (VEGF) and Angiopoietin 2 (Ang-2), were quantified by ELISA (Fig. 3). Generally, flow-conditioning (low and high flow) resulted in increased inflammatory signaling across all timepoints, particularly for IL8 (Fig. 3A), whereas MCP1 levels (Fig. 3B) dropped over time (although this was shown for both flow and static conditions). Ang-2 is upregulated in the case of flow-conditioned vessels (Fig. 3C), yet VEGF levels are significantly increased in statically cultured vessels over time (Fig. 3D).

**Figure. 3.**
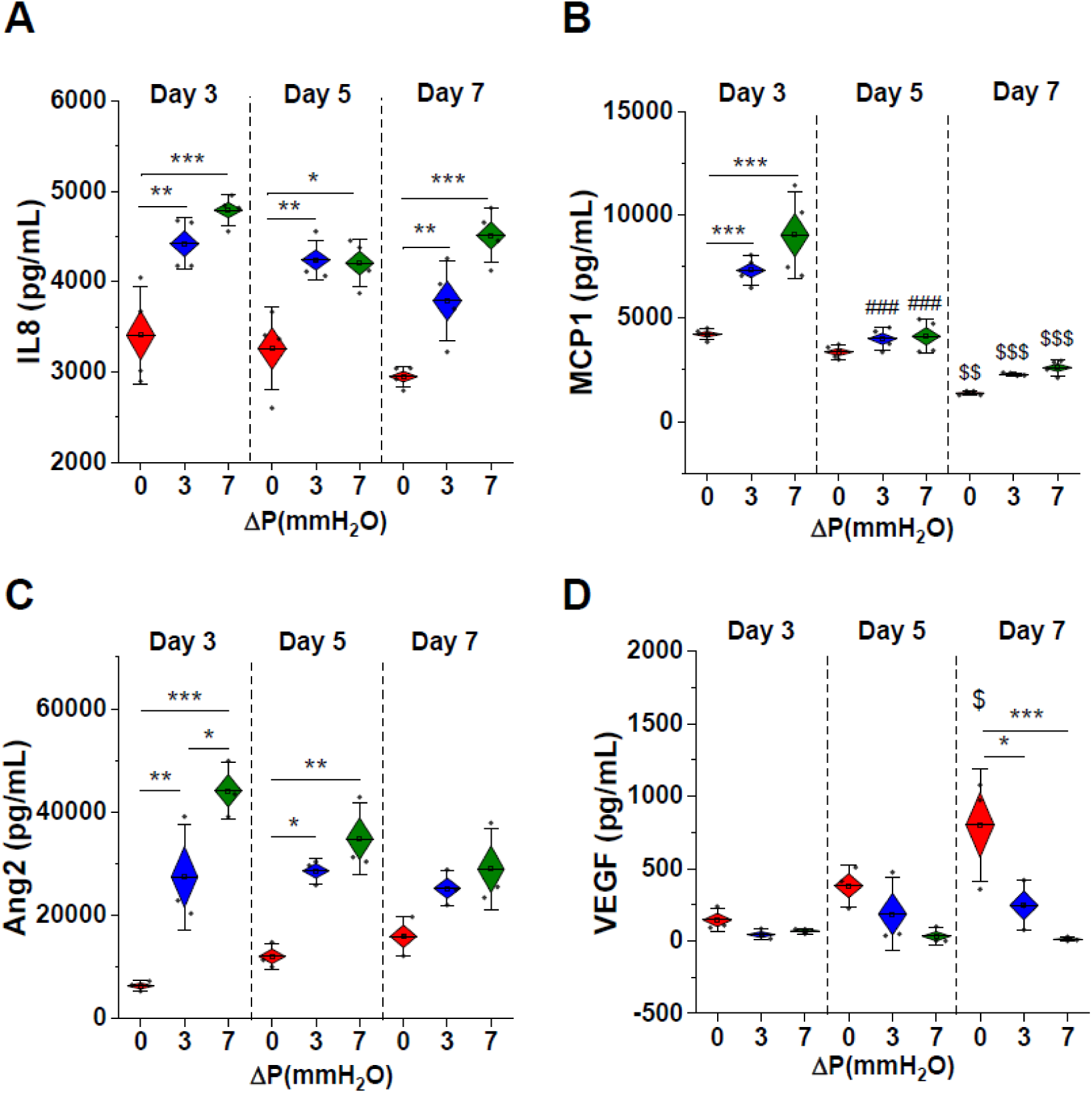
Interstitial flow impacts inflammatory and angiogenic signaling at early timepoints in vessel development. Concentrations of inflammatory molecules A) IL8 and B) MCP1 were measured by ELISA across various time points from 3D microvessels. Angiogenic signaling molecules C) Ang2 and D) VEGFa were significantly different from statically cultured vessels at early and later time points, respectively. Significance is measured by One-way ANOVA and indicated by *<0.05, **<0.01, and *** < 0.001 for Tukey means comparison test; # represents significance across day 5 and day 3 and $ across day 7 and day 3.

### 4. Flow sustains placental microvascular stability

We examined whether flow-conditioning could maintain structure and functionality in long-term culture of placental microvessels (> 7 days). Confocal images acquired at 14 days after seeding demonstrated distinct differences in vessel coverage between static and flow conditions (Fig. 4A). Microvessels maintained under static culture exhibited significantly reduced area coverage (Fig. 4B) and correspondingly narrow effective diameters (Fig. 4C). Branch density and length remained unchanged between static and flow conditions (Fig. 4D, E).

**Figure. 4.**
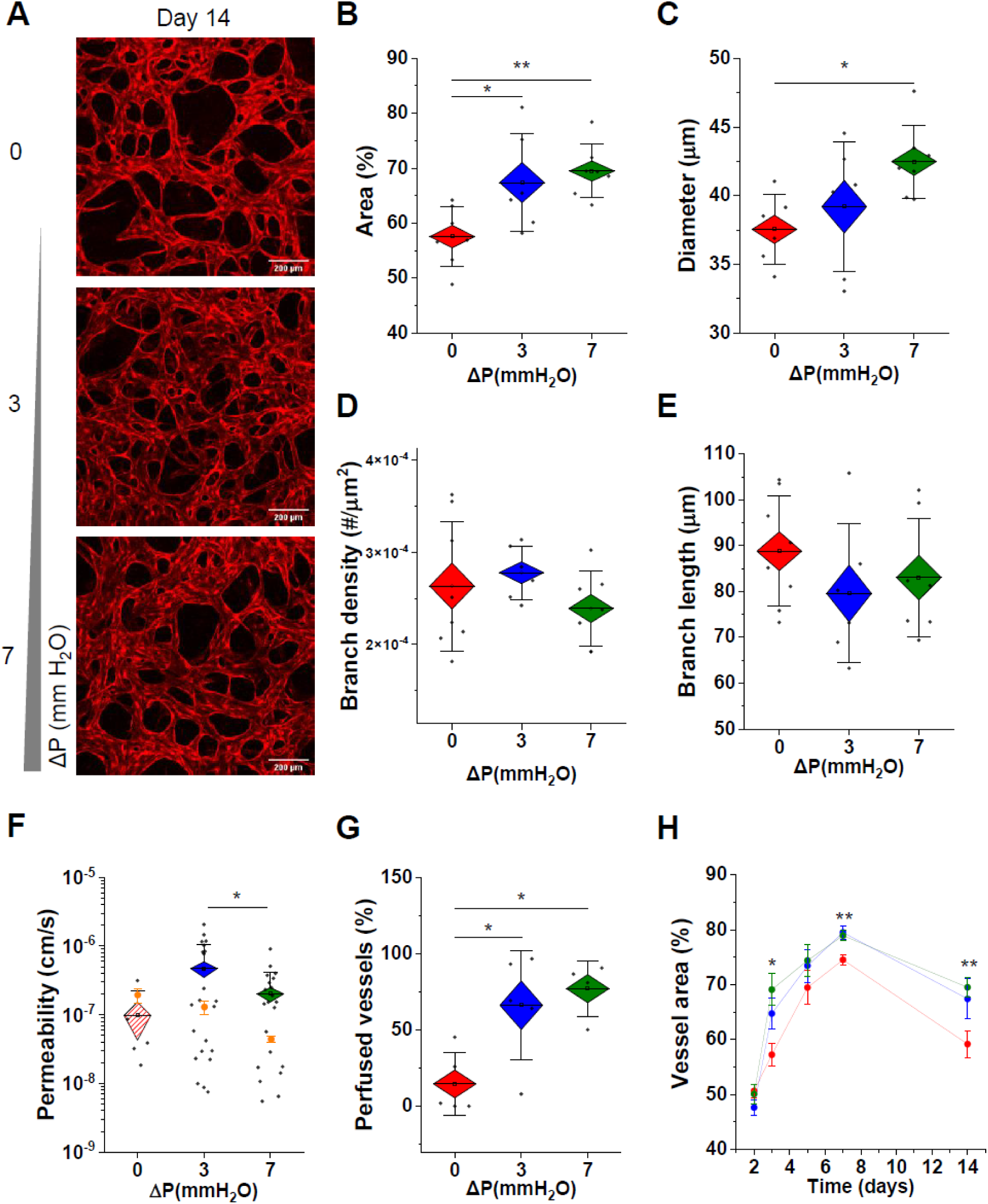
Flow promotes long-term stability of placental microvessels. A) Confocal images demonstrate differences in vascular density between static and flow-conditioned vessels at day 14 in culture. Both vessel B) area coverage and C) effective diameter are significantly increased by flow-conditioning. D) Branch density and E) length were not significantly impacted by flow. F) Vascular permeability is measured for fully perfused flow-conditioned vessels and the few perfusable vessels that remain in the static condition. Flow sustains vessel perfusion capacity after 14 days in culture (G) and higher area coverage H) overtime than in static condition. Significance is measured by One-way ANOVA and indicated by *<0.05 and **<0.01 for Tukey means comparison test.

Perfusion with 70 kDa FITC-dextran at day 14 (Fig. S4) demonstrated reduced leakiness in vessels cultured under high-compared to low-flow conditions. Permeability of statically cultured vessels was lower overall (increased barrier function); however, only a small fraction of vessels was perfused in this case (see Fig. 4G). Over time, vessel area and barrier function decrease in comparison to earlier timepoints (see Fig. 4H and orange dots from day 7 in Fig. 4F). Overall, flow-conditioning results in increased vascular coverage and allows for culture of connected vessels past 3-4 weeks (Fig. S5).

### 5. Characterization of velocity and shear stress distributions in placental microvessels

First, as a proxy for fluid velocity measurements, fluorescent beads were introduced into the microvessels at day 7 and tracked by time-lapse microscopy (Fig. S6A-B). Mean velocities were significantly increased for flow-conditioned vessels (Fig. S6C), corresponding to an increased effective diameter (Fig. 2C).

Next, to accurately map velocity and shear stress distributions within our microvessels, large sections of 70 kDa FITC-dextran-perfused devices were imaged and converted into binary masks for import as vectors into computational fluid dynamics (CFD) software. The computational modeling pipeline is outlined in Figure 5A. Using COMSOL, both the vessels and extravascular regions were modeled as distinct, yet integrated, domains. Employing the laminar flow physics module for flow through porous media, microvessel networks are treated as open pores filled with cell culture media (density = 998.2 kg/m^3^, viscosity = 9.4e^-4^ Pa*s). The extravascular matrix (EVM) is treated as a porous domain with fibrin gel (density = 985 kg/m^3^, viscosity = 1e^-2^ Pa*s) having a porosity of 0.3 and hydraulic permeability k=1e^-13^ m^2^ [35, 36]. In order to replicate the maximum pressure condition employed in the experiment, a simulated pressure gradient of 70 Pa (7 mm H_2_O) was applied across the gel region. From these simulations, both velocity and shear stress distributions are predicted for the combined vessels and EVM domains (Fig. 5 B, C). Estimated velocities and shear stresses for the complete tissue (vessels and gel combined) were lower overall than the vessel regions alone. Moreover, the gel regions demonstrate orders of magnitude lower values than the combined tissue (Table 1). CFD predicts heterogeneous flow patterns develop in these microtissues, which are contingent upon the changing interstitial to luminal flows at earlier time points (Fig. 1E).

**Table. 1.** Quantitative analysis of simulation outcomes. Surface averaging is applied to estimate the velocities and shear stresses (mean ± SD) within combined vessels + gel (tissue), vessels only and gel only regions.

|  |  | Tissue | Vessel region | Gel region |
| --- | --- | --- | --- | --- |
| Velocity<br>(mm/s) | S1 | 11.22 | 15.39 | 0.89 |
|  | S2 | 17.78 | 21.54 | 0.90 |
|  | S3 | 9.64 | 13.83 | 0.73 |
|  | S4 | 11.54 | 15.04 | 0.89 |
|  | Mean | 12.55 | 16.45 | 0.85 |
|  | ± SD | 3.11 | 3.00 | 0.07 |
|  |  | Tissue | Vessel region | Gel region |
| Shear<br>stress (Pa) | S1 | 5.09e<sup>-2</sup> | 29.60e<sup>-2</sup> | 1.2e<sup>-3</sup> |
|  | S2 | 7.50e<sup>-2</sup> | 38.30e<sup>-2</sup> | 1.4e<sup>-3</sup> |
|  | S3 | 5.06e<sup>-2</sup> | 31.18e<sup>-2</sup> | 1.0e<sup>-3</sup> |
|  | S4 | 5.30e<sup>-2</sup> | 30.91e<sup>-2</sup> | 1.6e<sup>-3</sup> |
|  | Mean | 5.74e<sup>-2</sup> | 32.50e<sup>-2</sup> | 1.3e<sup>-3</sup> |
|  | ± SD | 0.0102 | 0.0340 | 0.0002 |

**Figure. 5.**
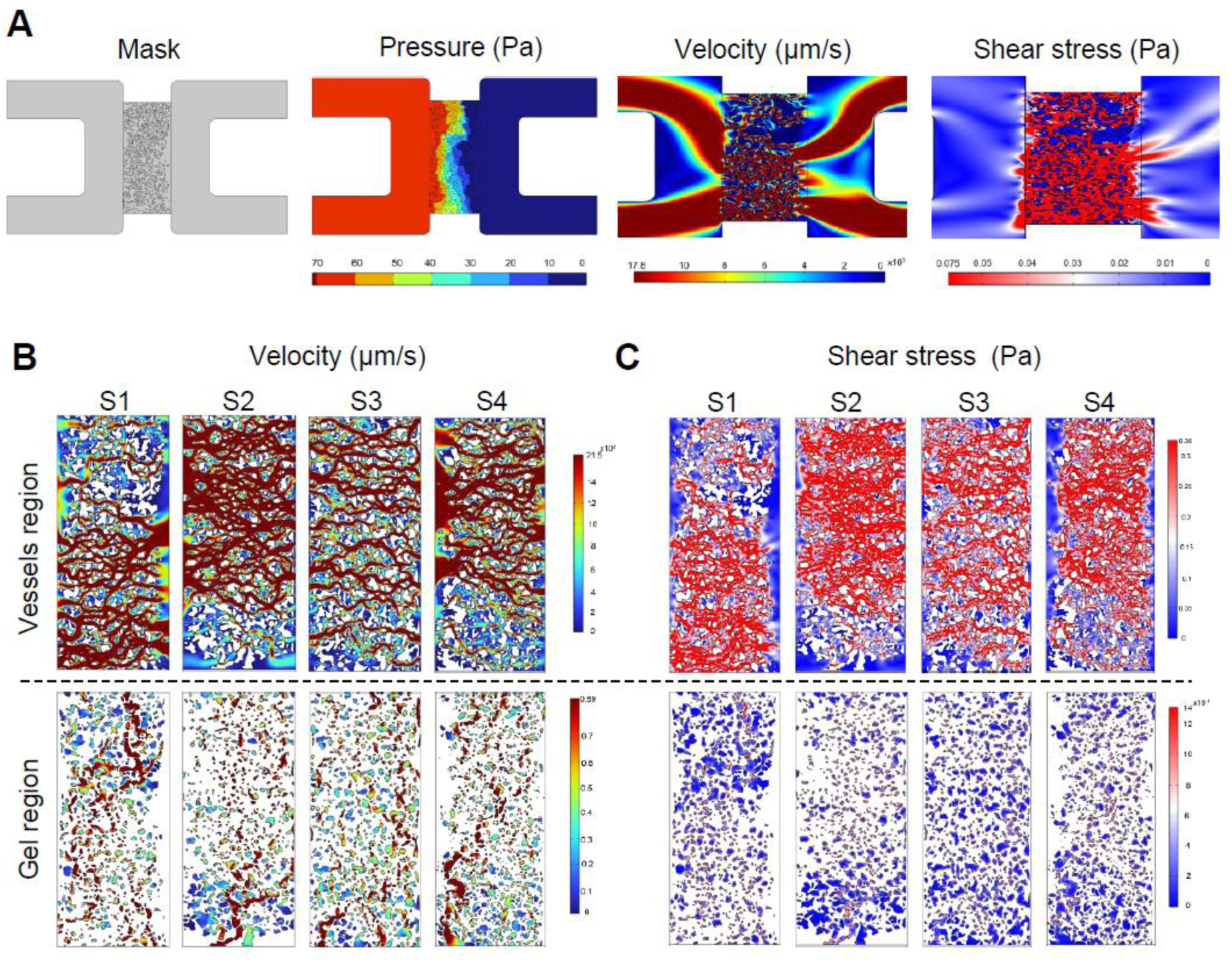
CFD simulation predictions of fetoplacental microvascular tissue. A) Pipeline for computational predictions of velocities and shear stresses within microvessels embedded in porous gel. Shown are 4 independent predictions from 7 mm H_2_O condition for B) velocity and C) shear stress distributions within microvessels and gel regions.

### 6. Flow-conditioning alters biophysical properties of the placental microvascular tissue

The EVM supplies mechanical and chemical cues that contribute to vascular network formation and barrier integrity [37]. We previously demonstrated significant changes in tissue stiffness and diffusivity in response to co-culture with specific fibroblasts [28]. Here, we assessed whether flow conditioning alone can drive these tissue-level changes. First, we examined diffusivity in extravascular regions from vessels cultured under static or flow conditions at day 7, as assessed by FRAP measurements (Fig. 6A). Flow-conditioning, at high flow, results in significantly reduced diffusivity in extravascular regions (Fig. 6B).

**Figure. 6.**
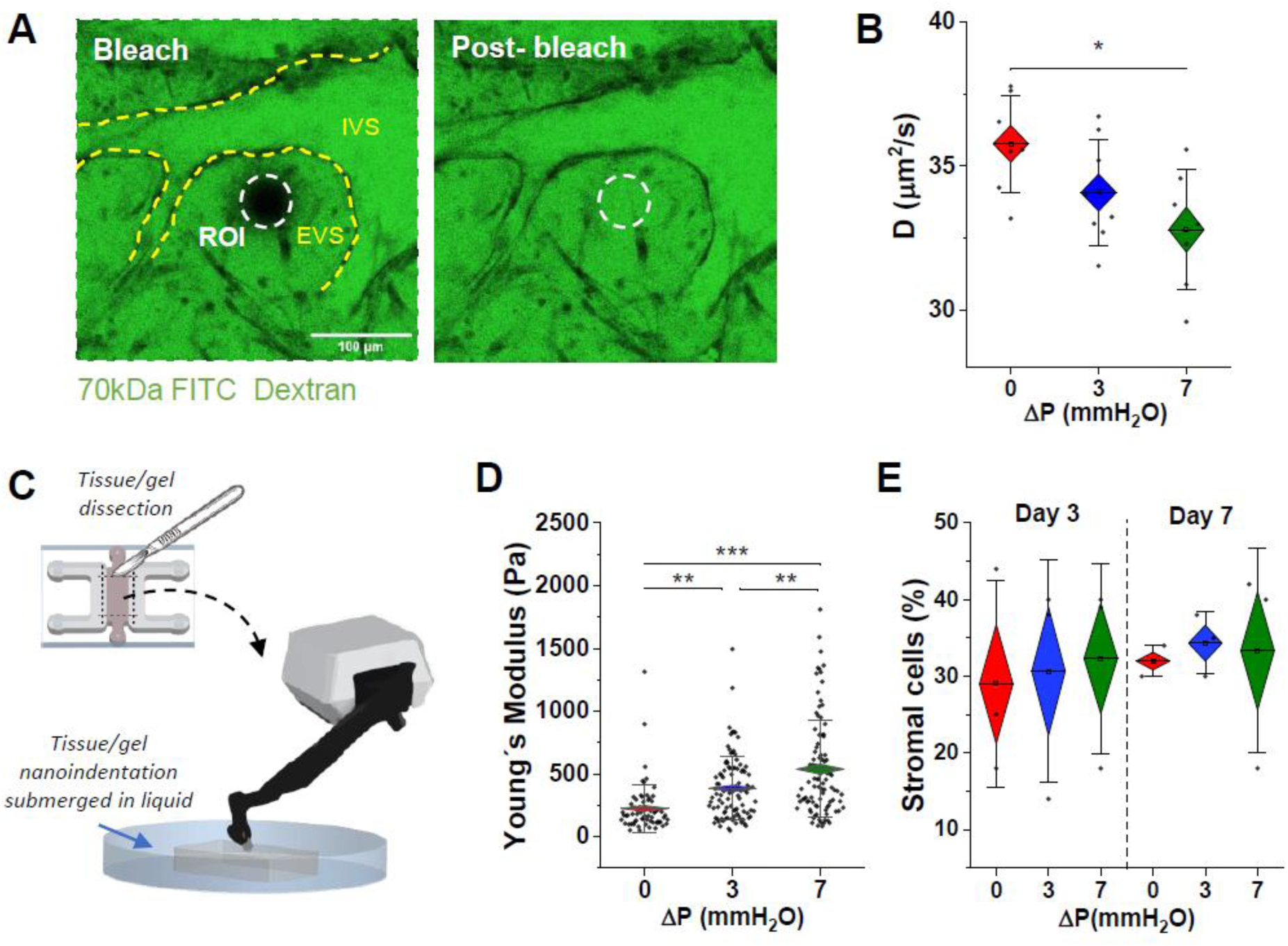
Interstitial flow alters extravascular remodeling and tissue-level stiffness. A) FRAP measurements performed on extravascular regions. ROI is bleached region of interest, EVS and IVS are extravascular and intravascular space respectively. B) Diffusivity measurements calculated from FRAP experiments for static and flow-conditioned microvessels at day 7. C) Schematic representation of nanoindentation on microtissues. D) Stiffness measurements performed across static and flow-conditioned samples. E) Populations of stromal cells from microvessels corresponding to different culture times. Significance is measured by One-way ANOVA and indicated by *<0.05, **<0.01, and *** < 0.001 for Tukey means comparison test.

Next, we assessed the impact of flow-conditioning on microvessel tissue stiffness. Apparent Younǵs moduli were assessed by nanoindentation at day 7 by exposing the tissue as shown in the schematic in Fig. 6C. Microvascular tissues cultured under flow are significantly stiffer than static ones (Fig. 6D), with stiffness dependent on the magnitude of flow (7 versus 3 mm H_2_O).

Since EVM is dynamic, constantly undergoing remodeling and able to activate cell proliferation and tissue morphogenesis [38], we assessed whether changes in tissue-level properties are associated with changes in the stromal cell population over time. To this aim, cells were extracted from the hydrogel matrix of static and flow conditioned devices 7 days after seeding to compare the cell composition over time. Cells were analyzed by flow cytometry (Fig. S7) across days; however, no differences in the percentage of stromal cells were found (Fig. 6E).

### 7. Flow promotes significant EVM protein deposition and remodeling

Tissue-level stiffness increase corresponded with decreased diffusivity in extravascular regions of flow-conditioned placental vessels. Since there are many identified proteins in the developing placenta, we next examined the impact of flow on protein production. After 7 days in culture, microvessels were fixed and stained for collagen I, laminin and fibronectin (Fig. 7A). In comparison to static conditions, all proteins assessed are more significantly expressed under flow-conditions (Fig. 7B).

**Figure. 7.**
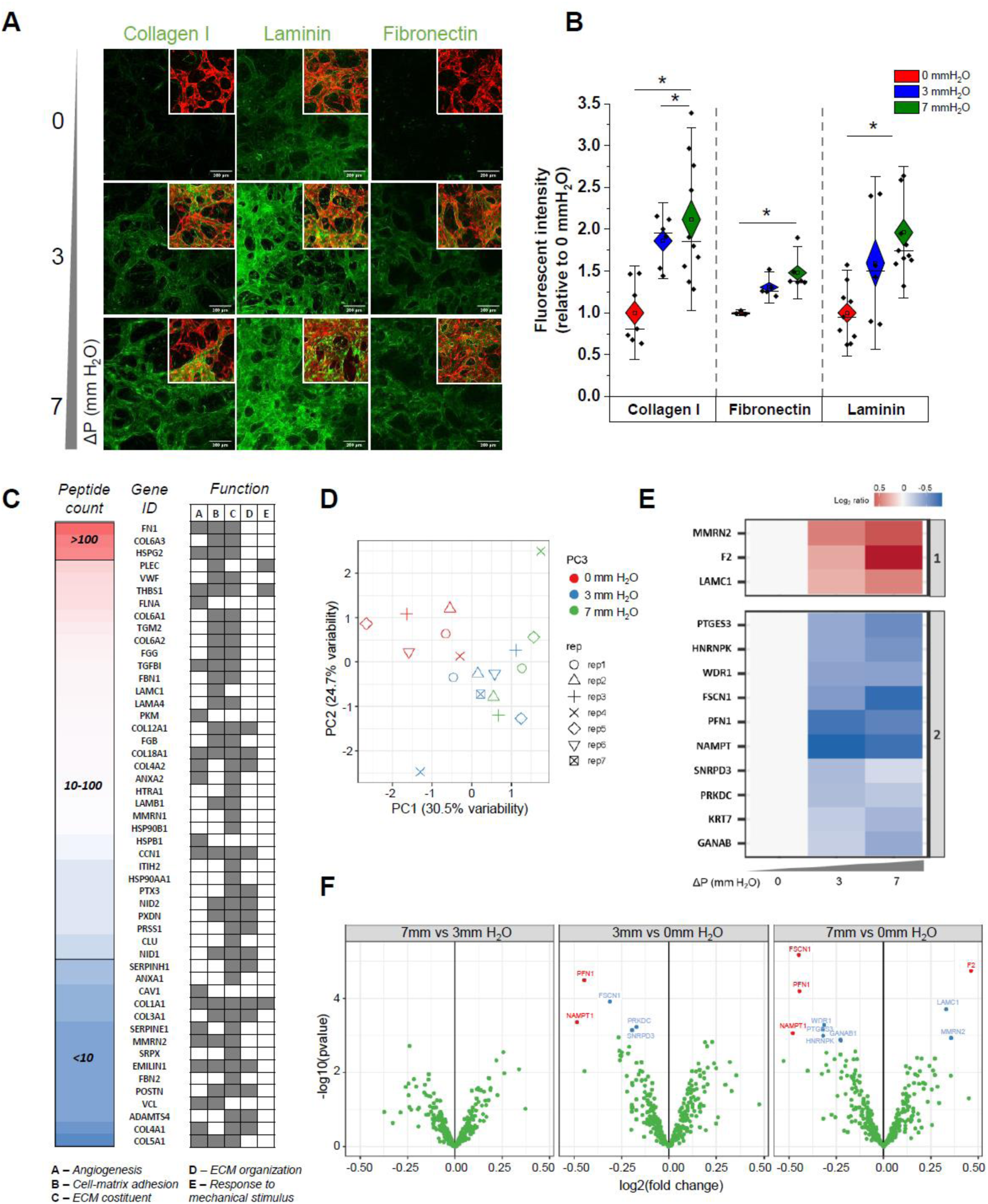
Flow induces changes in EVM protein deposition and composition. A) Confocal images at day 7 of static and flow-conditioned microvessels (red-labeling shown in the corner image insets) immunolabeled with collagen I, laminin and fibronectin (green-labeling). B) Fluorescence intensity of collagen I, fibronectin and laminin was measured and normalized to vessel area. Significance is measured by One-way ANOVA and indicated by *<0.05 for Tukey means comparison test. C) EVM protein composition based on the total peptide count after 7 days of culture, and ranked in a decreasing order of abundance (50 most abundant proteins). Proteins are categorized according to 5 ontology terms selected for their relevance to vascularization and matrix remodeling. D) PCA performed on mass spectrometry-based proteomics normalized data obtained from flow-conditioned samples for seven days. E) K-means clustering and heatmap of the 13 hit and candidate proteins, showing differential expression across the static and flow conditions (shown as the Log.2 ratio of protein abundance in each 3 and 7mm H_2_O sample relative to the average protein abundance in 0mm H_2_O samples). F) Volcano plots of the p values vs. the log2 protein abundance differences between flow and static conditions. Red dots, hits: FDR <0.05, FC >30%. Blue dots, candidates: FDR <0.05, no FC threshold. p values are calculated from moderated t-test (Limma).

To assess the EVM more broadly, we performed mass spectrometry analysis to measure compositional changes of the matrix under static and flow-conditions for vessels after 7 days in culture. Specifically, individual samples from each condition were subjected to tandem mass tag (TMT) labeling, followed by multiplexing, and ultimately measured in a mass spectrometry-based experiment. As a result, we identified a total of 804 proteins, of which 333 were quantified (Data S1), with many showing relevance to vascularization, EVM composition and organization, as classified by gene ontology (Fig. 7C). For each protein, the fold change in expression between conditions was calculated and ratios of protein expression were obtained (by dividing the flow conditioned samples by the corresponding static counterparts). Principle component analysis showed that static and flow-conditioned sample replicates were clustered, consistently with their experimental condition (Fig. 7D). Thus, we identified 13 proteins with a fdr below 0.05, which were clustered into two different groups based on their expression similarities and behaviors under different conditions (Fig. 7E). No significant differences were observed between flow conditions. However, both the 3– and 7-mm H_2_O conditions exhibited notable variations in protein expression when compared to the static counterpart as shown in the volcano plots analysis (Fig. 7F). In particular, we observed that flow-conditioned 7 mm H_2_O tissue exhibited increased expression of EVM components and known regulators of matrix stability such as prothrombin (F2, identified as a hit), laminin (LAMC1, candidate) and multimerin 2 (MMRN2, candidate). Conversely, we observed a significant decrease in the expression levels of the NAD biosynthetic enzyme nicotinamide phosphoribosyltransferase (NAMPT, hit), as well as proteins associated with actin dynamics and cytoskeleton organization, namely profilin (PFN1, hit) and fascin (FSCN1, hit), in the presence of flow. These findings suggest that flow conditions can modulate the expression of key factors involved in EVM integrity and cellular cytoskeletal dynamics.

## Discussion

Limited access to human placentae, particularly at early stages of pregnancy, impedes the elucidation of mechanisms associated with fetoplacental vascular development. To circumvent this challenge, this study employs a 3D placental-specific microvascular tissue on-chip. Building on our previous model [25], we generated an all-placental vascular microtissue and incorporated a flow reservoir, allowing us to characterize vasculogenesis and extravascular remodeling processes in response to flow conditions. Incorporating placental fibroblasts, as opposed to co-culture with lung fibroblasts as done previously [25], our model required parameter adjustment in order to achieve perfusable vessels. In this tri-culture, placental fibroblasts associate with microvessels, as do pericytes, as also shown in our previous model [25]. Both studies have shown that stromal cells have a clear role in contributing to fetoplacental vascular growth and remodeling.

Besides what is known about the stroma, little is known about placental hemodynamics in early pregnancy, particularly in the villous vasculature. Ultrasound imaging has shown that blood flow velocity waveforms of the umbilical artery, and its branches, change with advancing gestation and correlate with the development of the placental villous trees and capillary networks [39]. Despite the presence of a complete vascular network within the villi, it is believed that the chorionic circulation is not fully established until the end of the first trimester. This flow is progressively established in the third month of gestation, together with the perfusion of the maternal blood into the intervillous space [40]. The precise mechanisms behind the formation and remodeling of the villous vascular network remain unknown. Nevertheless, we hypothesized that hemodynamic forces and associated signaling could play a role in directing vasculogenesis. Previous work has shown that interstitial flow promotes early vessel connectivity in 3D vasculature on-chip [16]. Herein, static, low and high interstitial flow velocities ranging from ∼0.1-1.2 µm/s (day 2 measurements) were applied via reservoirs (Fig. S3). To the best of our knowledge, there are no reports on human fetoplacental interstitial fluid velocity; however, our measurements fall within the physiological range found in most soft tissues (from 0.1 to 4.0 μm/s) [41, 42]. For these placental-like vessels, interstitial flow applied at early timepoints is necessary to establish fully perfusable vascular networks, which transition to luminal flow by day 5 after seeding (Fig. 1E). Consistent with our observations, the flow conditions employed in our experiments effectively induce vasculogenesis. This is evident from the shear stress experienced by the cells within the gel and the Peclet number (Pe), which were quantified using the equations described in [13]. Our assessment revealed that the 3 and 7 mmH_2_O gradients generated shear stresses of 1.32e^-5^ and 3.75e^-5^ Pa, respectively. Additionally, by considering the average interstitial fluid velocity (0.30 µm/s and 1.23 µm/s), the length of the central channel in the device (3 mm), and the diffusivity of a 70 kDa solute in the gel as measured by FRAP (D = 34.5 μm^2^/s), we determined that the interstitial fluid conditions corresponding to 3 and 7 mmH_2_O yielded Pe values of approximately 26 and 107, respectively. These Pe values exceed the reported threshold (Pe > 10, [34]) required to trigger a vasculogenic response. Flow-conditioning has a clear impact on vessel morphology, resulting in earlier connectivity, reduced branch density and larger vessel diameters (Fig. 2). Importantly, endothelial barrier function significantly improved in flow-conditioned vessels (Fig. 2E), compared to static cultures. Other studies conducted in perfused placental models have also demonstrated that shear stress (20 dyne/cm^2^ or 2 Pa) promotes vasodilatation (nitric oxide release) and decreases vascular resistance [43, 44]. Our results demonstrate the necessity for flow and a magnitude-dependence to ensure development of perfusable fetoplacental-like vessels with improved barrier function. In line with this, vascular abnormalities were found in fetal growth restriction-affected placentae [45], which are known to be characterized by high vascular resistance and fetoplacental hypoperfusion. While speculative, our findings may implicate the need for flow early in fetal vessel development to prevent gestational complications.

Flow induces changes in vascular function and remodeling by promoting inflammatory and angiogenic responses [46, 47]; however, insight into flow-induced signaling in the placenta remains limited. Pro-inflammatory chemokines, including IL-8 and MCP-1 are associated with angiogenesis, as evidenced by various studies [48, 49]. Sustained levels of these chemokines are produced by the placenta to stimulate the immune response against potential infections during pregnancy [50, 51], but it remains unclear what role, if any, these chemokines play in the development of placental vasculature. Our findings indicate that IL-8 and MCP-1 are strongly influenced by flow, aligning with previous reports of their regulation by shear stress [52–54]. The human placenta has also been found to be a significant source of locally produced angiogenic factors which include members of the VEGF family, FGF family, and angiopoietins, among others [55]. We analyzed the levels of Ang-2 in our fetoplacental vessels, which is highly expressed in early gestation and promotes vascular remodeling in the presence of VEGF [56, 57]. Our findings indicate increased Ang-2 expression as a result of flow-conditioning at early timepoints (Fig. 3D), which contrasts previous research showing a decrease in shear-stress dependent expression of Ang-2 in experiments with flow-exposed HUVEC monolayers [58, 59]. It is worth noting that previous studies investigating Ang-2 have primarily focused on laminar flow conditions (resulting in wall shear stress, WSS, >6 dyne/cm^2^ or 0.6 Pa), whereas other studies have demonstrated that Ang-2 expression is upregulated at lower shear stress levels (1 dyne/cm^2^) [60]. Notably, the expression of Ang-2 in fibroblasts has been previously found to elicit vascular growth, while concurrently maintaining a balance in the quiescent action exerted by Ang-1, as produced by placental pericytes [25]. These observations support our findings and highlight the need for further research to fully understand the complex relationship between shear stress and Ang-2 expression. In particular, VEGF and its receptors are expressed in trophoblasts and villous fetal vessels at early developmental stages, suggesting their involvement in the initiation and progression of vasculogenesis [61]. Herein, we observed elevated VEGF levels at later stages (day 7) within static cultures, proposed as a compensatory mechanism to counteract the absence of flow. In line with this hypothesis, elevated VEGF expression was found in placentae from pregnancies with fetal growth restriction, suggesting that a decrease in fetal-maternal blood circulation during placentation enhances the expression of the angiogenic factor [62].

Vessel stabilization is driven by hemodynamic forces through regression and pruning of branches exposed to low blood flow and by maintenance of vessel connections above a shear flow threshold [63]. After two weeks of culture under intermittent flow, placental-like microvessels were perfusable and functional (maintained a relatively low permeability) (Fig. 4 and Fig. S4), consistent with observations in a brain 3D microvascular system [23]. Although vessel branching did not differ, flow significantly preserved vessel diameter and prevented vascular narrowing, as opposed to the vessel constriction observed in static cultures (Fig. 4H), which decreased from ̴74.5% to ̴59% area coverage from day 7 to 14. Previous observations in HUVEC and lung co-cultures have also shown that continuous flow (at a low WSS of <10 dyne/cm^2^ or 1 Pa) maintains a stable vascular diameter in formed vessels [16]. At this late stage of culture, very few static-cultured vessels were perfusable, and interestingly those few vessels maintained a high barrier function (Fig. 4F). However, day 14 flow-conditioned vessels exhibited a slight decrease of endothelial barrier capacity with respect to day 7 (Fig. 4F, layered data plot in orange) that we attribute to the limited stimuli experienced by the vasculature upon reestablishment of the hydrostatic pressure (every other day) from day 7 onward. In addition, flow-conditioning was essential in maintaining microvessels connectivity over 4 weeks in culture (Fig. S5), highlighting the importance of implementing flow for long-term *in vitro* microvessel studies.

Luminal flow in the vessels was estimated by particle tracking, which demonstrated a significant increase in the mean velocity in flow-conditioned vessels by day 7 (Fig. S6). To obtain a more accurate prediction, vascular network flow was modeled in COMSOL (Fig. 5). Heterogeneous flow distributions were apparent in the networks, which depend on the formation of open lumens in the media channels. Shear stress for flow-conditioned vessels was predicted to be 0.32 Pa (3.2 dyne/cm^2^), whereas shear stress within the EVM was 0.001Pa (0.01 dyne/cm^2^). CFD results deviated from experimental bead measurements by approximately one order of magnitude. Note that velocities and shear stress predictions are obtained using 2D segmented projections and do not account for the exact spatial complexity of 3D vessels, nor the wall effects or size-dependent effects of beads. It was reported previously that the presence of common additives in the media [64], tracer beads size, and their properties [65], such as deformability, contribute to elastic no-slip boundaries at the fluid interfaces influence the velocities.

We postulated that differences in vascular morphogenesis and long-term stability, observed in the presence of flow, could be due to extravascular rearrangements and changes in the EVM, as previously reported [66]. Recent findings showed that flow-conditioning reduces proteolytic activity of cathepsins prolonging *in vitro* microvessel stability [67]. Flow-conditioned vessels, as opposed to the static ones, exhibited significantly reduced extravascular diffusivity and overall increased matrix/tissue rigidity (Fig. 6). Despite evidence showing that stromal cells can contribute to tissue stiffening [28], the flow-dependent increase in stiffness is not attributable to alterations in stromal cell population, since they remain stable (̴30% of the total cell population) over time (Fig. 6E). Examination of proteins in the EVM by immunostaining revealed that flow promotes increased deposition of collagen I, fibronectin, and laminin (Fig.7A-B). Several studies have reported the effect of shear stress in regulating matrix deposition and remodeling [68–70], but none in the context of placental tissue. Collagens, fibronectin, and laminins are abundant in the placental stroma and basement membrane, and their expression increases with advancing gestation to support the developing tissue structure and function [71]. Low expression of these EVM proteins, observed in the absence of flow, could explain their increased destabilization over time. A recent study reported that endothelial cells plated on matrices derived from villous stromal fibroblasts where fetal growth restriction was clinically observed (reduced expression of collagen I and fibronectin) exhibited impaired proliferation and migration, suggesting that matrix composition is crucial for fetoplacental vascular development [72]. A more in-depth examination of the gel/tissue composition utilizing mass spectrometry, revealed an abundance of EVM-related proteins deposited by the cells over the culture period (Fig. 7C). As expected, tissue protein enrichment is influenced by the presence of the flow, revealing changes in protein expression associated with EVM organization and vascular homeostasis. Consistent with the immunofluorescence findings, MS analysis revealed enrichment in laminin in response to flow, indicating flow-induced adaptation and alterations in cell-matrix adhesion, which ultimately resulted in matrix remodeling. In addition, multimerin 2, an EVM molecule that enhances vascular stability and regulates permeability by stabilizing endothelial junctions [73, 74], was found enriched in flow-conditioned tissues suggesting a protective function in maintaining vessel function and stability under mechanical shear stress. The substantial increase of prothrombin, which serves as a precursor to thrombin and facilitates fibrin formation [75], provides further evidence of flow-induced enhancements in EVM rigidity and subsequent matrix stabilization. Flow conditions also resulted in the depletion of nicotinamide phosphoribosyltransferase (NAMPT), an essential coenzyme that plays a critical role in energy production, DNA repair, and signaling pathways. The overexpression of NAMPT has been associated with inflammatory processes and the development of various human conditions, including acute lung injury, atherosclerosis, and cancer [76]. While direct evidence of the specific effects of flow on NAMPT expression is lacking, our findings suggest that the protein is mechanosensitive, indicating a potential protective role in response to flow mechanical forces. Moreover, flow has an impact on actin dynamics as indicated by the reduced levels of profilin and fascin in flow-conditioned tissue. Based on previous reports [77–79], we propose that the depletion of these actin-bundling proteins can alter the organization of the cytoskeleton and hinder cell migration. Consequently, once the perfusable vascularized tissue is formed, the absence or reduction of cellular movement plays a role in ensuring the functionality and stability of the fetoplacental vascular barrier.

To the best of our knowledge, this study is the first to examine the effects of flow on the formation of fetoplacental microvessels and modulation of their extravascular matrix. Although intermittent flow only partially reproduces the hemodynamic forces *in vivo* [19, 80], the predicted shear stress generated in our system is sufficient for inducing perfusable, branched fetal-vascular networks that are stable for several weeks in culture. Our tri-culture system is more physiologic than previous methods; but does lack the branched villous structure and critical epithelial layer (trophoblasts) that forms a barrier between the maternal blood and fetal capillaries. Since trophoblasts are mechanosensitive to shear stress and release angiogenic factors [81], this model will be useful for exploring their role in fetal villous vascular development.

## Acknowledgments

We would like to acknowledge the Flow Cytometry Unit of the Center for Genomic Regulation for consultation in data acquisition and analysis, Akinola Akinbote (Haase lab) for helping with the initial set up of the nanoindentation experiments, and Roberto Paoli (EMBL Barcelona) for helping with reservoir fabrication.

## Author contributions

Conceptualization, M.C. and K.H.; Methodology, M.C., K.H., S.E., P.P., P.H., F.S., V.B.; Experimentation, M.C., S.E., P.P., P.H.; Validation: M.C., K.H., P.P., P.H., F.S.; Writing – original draft, M.C.; Writing—review & editing: M.C., K.H., S.E., P.P., P.H., F.S.; Funding Acquisition, K.H.; Supervision, K.H.

## Data and materials availability

All data needed to evaluate the conclusions in the paper are present in the paper and/or the Supplementary Materials.

