## Supplementary material for "Flow in fetoplacental microvessels in vitro enhances perfusion, barrier function, and matrix stability"

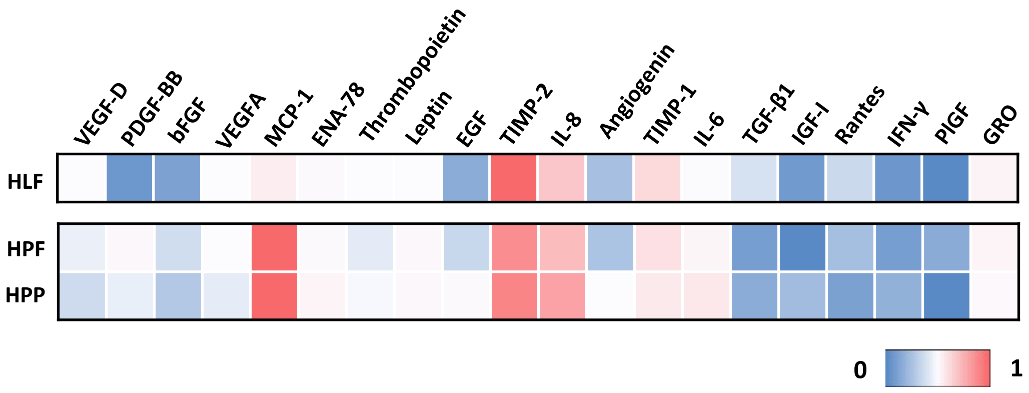


**Figure S1.** Cytokine expression of stromal cells in 2D culture. An array of cytokines are shown by relative expression levels for human lung fibroblasts (HLF), human placental fibroblasts (HPF), and human placental pericytes (HPP).

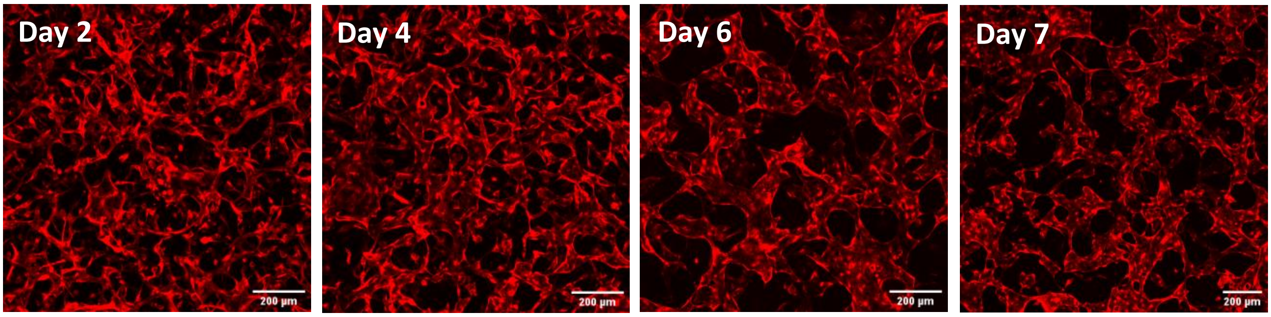


**Figure S2.** Fluorescent images demonstrating microvessel development in the absence of media reservoirs. Vessel connect and form without flow and reduced media volume; however, the connectivity is low as seen by day 6 and 7 and they are not perfusable. HUVEC are shown in red (cytoplasmic label).

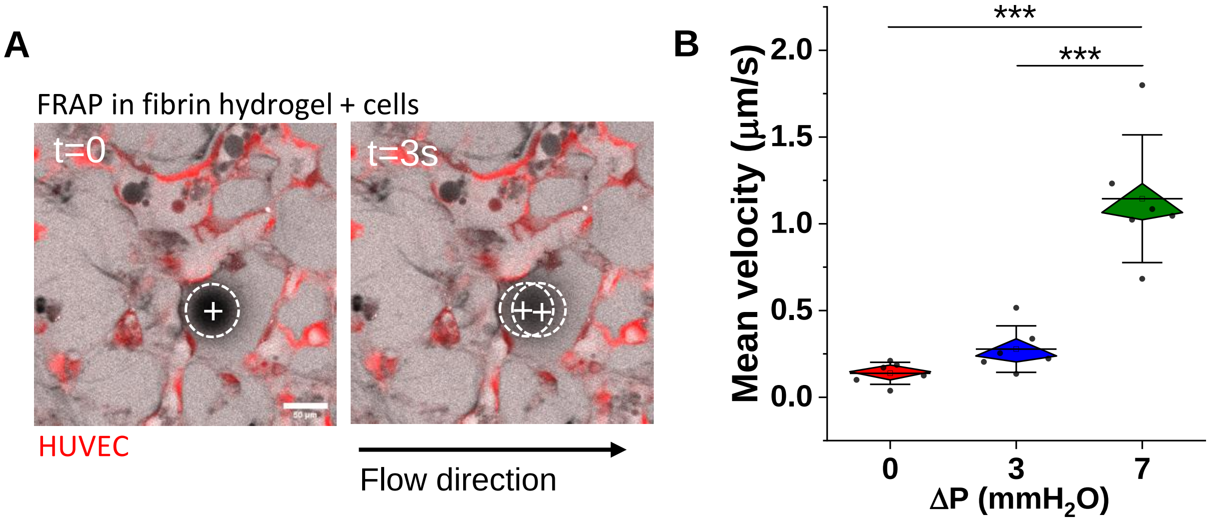


**Figure S3.** Assessment of interstitial flow velocity at day 2. A) FRAP measurements were performed in fibrin hydrogels with cells (tri-culture) to measure B) mean interstitial velocity for different fluid volume change (pressure difference across the gel). At this stage, vessel are not yet formed.

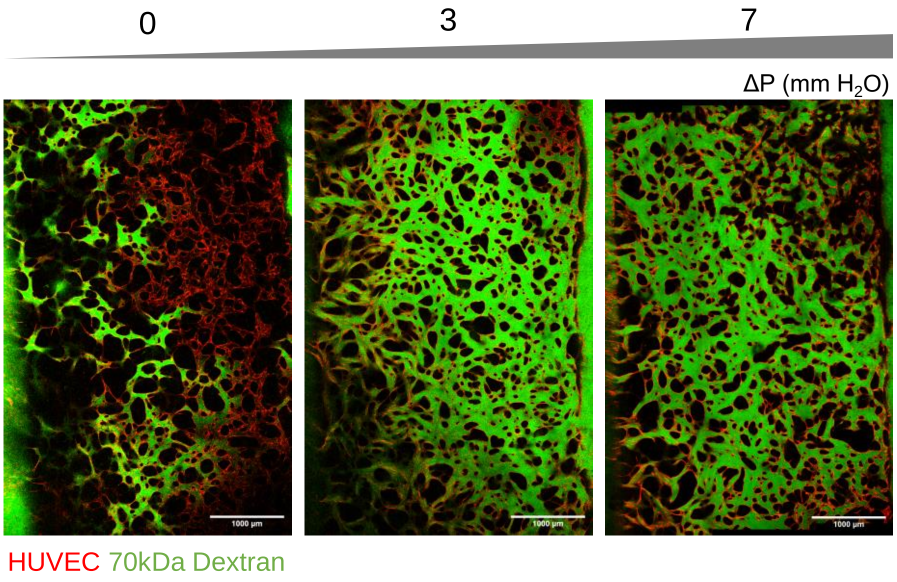


**Figure S4.** Long-term culture with flow maintains perfusion capacity of microvessels. Shown are overview images of example microvascular beds cultured under static and flow conditions demonstrating the complete perfusion for flow-conditioned vessels and only partial perfusion for static cultured placental vessels at day 14.

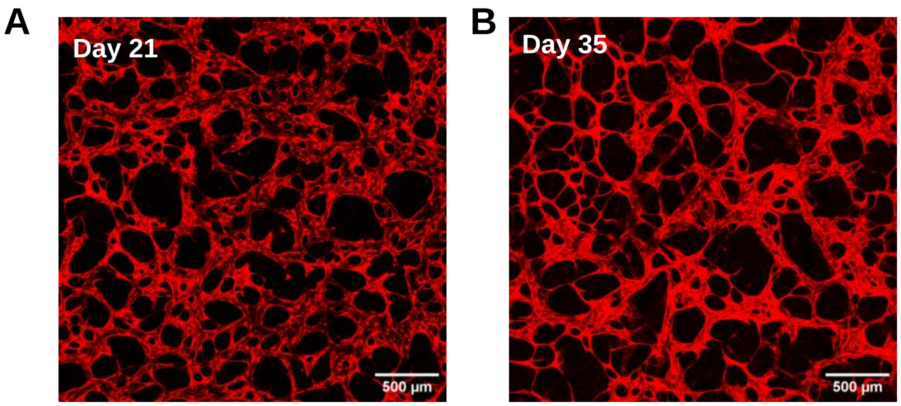


**Figure S5.** Intermittent flow-conditioning Flow-conditioning enables long-term stability of vessels to A) 3 weeks and B) past 4 weeks in culture. Beyond 4 weeks vessels remain connected but begin to narrow in diameter. HUVEC are shown in red (cytoplasmic label).

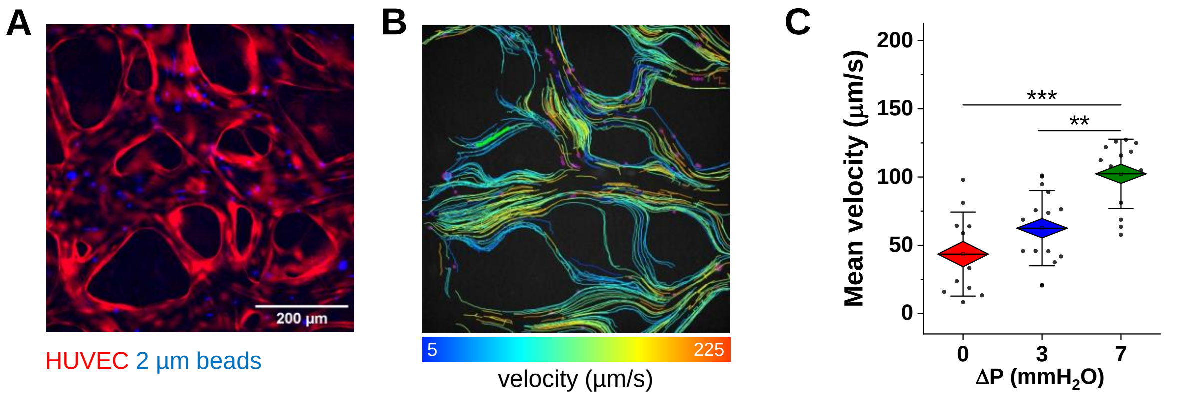


**Figure S6.** Characterization of luminal flow reveals heterogeneous patterns. A) Representative image of fluorescent tracer beads in lumen in day 7 vessels cultured under flow conditions (7 mm$H_2O$). B) Example of tracked paths from beads in A. C) Quantification of mean bead velocities from measurements in static and flow-conditioned microvessels subjected to a ΔP of 5 mm$H_2O$. Significance is measured by One-way ANOVA and indicated by **<0.01, and *** < 0.001 for Tukey means comparison test

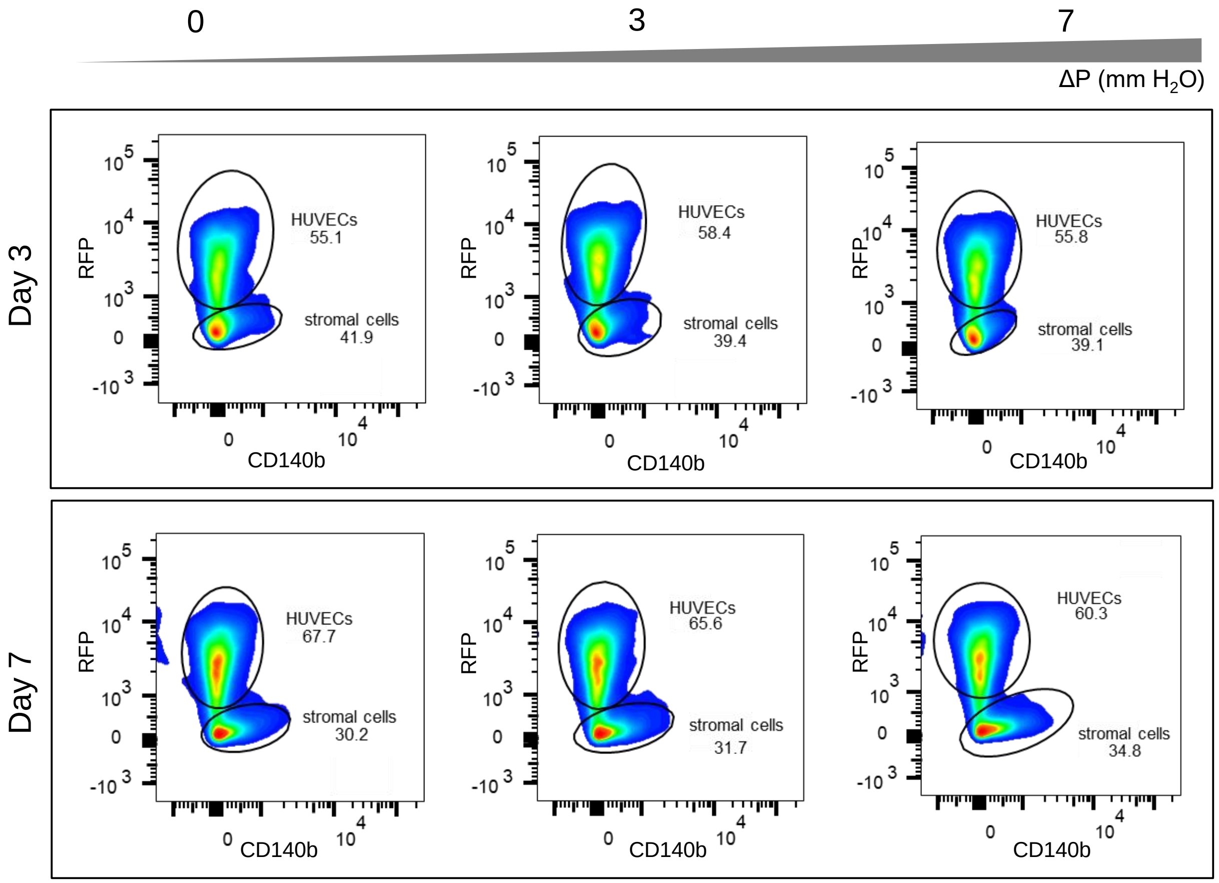


**Figure S7.** Flow cytometry of endothelial and stromal populations over time. Density plots are shown for static and flow-conditioned vessels at days 3 and 7. Stromal cell population was gated based on $RFP^-$ (expressed only in HUVEC) and CD140b (pericyte marker) expression. This figure represents one experiment, however 4 independent experiments were performed.

**Supplementary data**

**Data S1. List of identified protein and relative Limma analysis results**

| Gene | Protein ID | logFC | AveExpr | comparison | pvalue | fdr |
|---|---|---|---|---|---|---|
| FSCN1 | Q16658 | -0.311239042 | 17.22349762 | 3 m$H_2O$ vs 0 mm $H_2O$ | 0.000122123 | 0.020333517 |
| PRKDC | P78527 | -0.170359196 | 19.38685821 | 3 m$H_2O$ vs 0 mm $H_2O$ | 0.000595418 | 0.048257709 |
| SNRPD3 | P62318 | -0.193517279 | 19.04018698 | 3 m$H_2O$ vs 0 mm $H_2O$ | 0.00072459 | 0.048257709 |
| LAMC1 | P11047 | 0.33022331 | 21.18750318 | 7 m$H_2O$ vs 0 mm $H_2O$ | 0.000198976 | 0.016564752 |
| WDR1 | O75083 | -0.314267506 | 18.94923163 | 7 m$H_2O$ vs 0 mm $H_2O$ | 0.000520806 | 0.034685653 |
| PTGES3 | Q15185 | -0.322230437 | 20.03155888 | 7 m$H_2O$ vs 0 mm $H_2O$ | 0.000678983 | 0.037683573 |
| HNRNPK | P61978 | -0.319956645 | 19.95715687 | 7 m$H_2O$ vs 0 mm $H_2O$ | 0.001022726 | 0.041346123 |
| MMRN2 | Q9H8L6 | 0.356334211 | 19.6353503 | 7 m$H_2O$ vs 0 mm $H_2O$ | 0.001189025 | 0.041346123 |
| GANAB | Q14697 | -0.22719015 | 19.37791789 | 7 m$H_2O$ vs 0 mm $H_2O$ | 0.001304175 | 0.041346123 |
| KRT7 | P08729 | -0.225702466 | 19.43973796 | 7 m$H_2O$ vs 0 mm $H_2O$ | 0.001365788 | 0.041346123 |
| PFN1 | P07737 | -0.445904779 | 17.42183319 | 3 m$H_2O$ vs 0 mm $H_2O$ | 3.21E-05 | 0.010685022 |
| NAMPT | P43490 | -0.484561988 | 15.42557579 | 3 m$H_2O$ vs 0 mm $H_2O$ | 0.000444396 | 0.048257709 |
| FSCN1 | Q16658 | -0.448521817 | 17.22349762 | 7 m$H_2O$ vs 0 mm $H_2O$ | 6.77E-06 | 0.002252928 |
| F2 | P00734 | 0.460961056 | 16.10657822 | 7 m$H_2O$ vs 0 mm $H_2O$ | 1.80E-05 | 0.00299408 |
| PFN1 | P07737 | -0.445102778 | 17.42183319 | 7 m$H_2O$ vs 0 mm $H_2O$ | 6.39E-05 | 0.007097386 |
| NAMPT | P43490 | -0.478777915 | 15.42557579 | 7 m$H_2O$ vs 0 mm $H_2O$ | 0.000863648 | 0.041084968 |
| WDR1 | O75083 | -0.265809268 | 18.94923163 | 3 m$H_2O$ vs 0 mm $H_2O$ | 0.00113256 | 0.062857079 |
| TMED2 | Q15363 | 0.230057385 | 17.21365208 | 3 m$H_2O$ vs 0 mm $H_2O$ | 0.001514619 | 0.064038381 |
| RPL31 | P62899 | 0.192198321 | 20.04161035 | 3 m$H_2O$ vs 0 mm $H_2O$ | 0.00153846 | 0.064038381 |
| IKBIP | Q70UQ0 | 0.201276041 | 16.34942842 | 3 m$H_2O$ vs 0 mm $H_2O$ | 0.001984753 | 0.068107507 |
| P4HB | P07237 | -0.208783845 | 19.25928557 | 3 m$H_2O$ vs 0 mm $H_2O$ | 0.00204527 | 0.068107507 |
| DHX9 | Q08211 | -0.261672967 | 16.0006753 | 3 m$H_2O$ vs 0 mm $H_2O$ | 0.002616897 | 0.075741114 |
| PTGES3 | Q15185 | -0.247804013 | 20.03155888 | 3 m$H_2O$ vs 0 mm $H_2O$ | 0.002879286 | 0.075741114 |
| MDH2 | P40926 | -0.214276749 | 18.21421021 | 3 m$H_2O$ vs 0 mm $H_2O$ | 0.003048225 | 0.075741114 |
| NONO | Q15233 | -0.251033949 | 19.26168669 | 3 m$H_2O$ vs 0 mm $H_2O$ | 0.003473754 | 0.075741114 |
| HSP90B1 | P14625 | -0.25544991 | 20.36585658 | 3 m$H_2O$ vs 0 mm $H_2O$ | 0.003514477 | 0.075741114 |
| HNRNPK | P61978 | -0.250395114 | 19.95715687 | 3 m$H_2O$ vs 0 mm $H_2O$ | 0.003639213 | 0.075741114 |
| HNRNPA2B1 | P22626 | -0.258959563 | 18.53624742 | 3 m$H_2O$ vs 0 mm $H_2O$ | 0.004681953 | 0.083704152 |
| VCAN | P13611 | -0.14419221 | 19.33335978 | 3 m$H_2O$ vs 0 mm $H_2O$ | 0.005162574 | 0.083704152 |
| BCAP31 | P51572 | 0.318831663 | 19.52713253 | 3 m$H_2O$ vs 0 mm $H_2O$ | 0.00520154 | 0.083704152 |
| PFKP | Q01813 | -0.143275956 | 20.72340758 | 3 m$H_2O$ vs 0 mm $H_2O$ | 0.005462831 | 0.083704152 |
| RAN | P62826 | -0.170320053 | 20.33795383 | 3 m$H_2O$ vs 0 mm $H_2O$ | 0.005566116 | 0.083704152 |
| RPL7A | P62424 | 0.238847995 | 20.68475958 | 3 m$H_2O$ vs 0 mm $H_2O$ | 0.005746101 | 0.083704152 |
| CKAP4 | Q07065 | 0.156320408 | 20.5524488 | 3 m$H_2O$ vs 0 mm $H_2O$ | 0.005781368 | 0.083704152 |
| F2 | P00734 | 0.20543168 | 16.10657822 | 3 m$H_2O$ vs 0 mm $H_2O$ | 0.007435729 | 0.102436971 |
| COX4I1 | P13073 | 0.251123214 | 18.4693388 | 3 m$H_2O$ vs 0 mm $H_2O$ | 0.007690463 | 0.102436971 |
| COX5B | P10606 | 0.163862747 | 18.89368926 | 3 m$H_2O$ vs 0 mm $H_2O$ | 0.008494673 | 0.102864098 |
| EHD2 | Q9NZN4 | -0.15150935 | 17.32935895 | 3 m$H_2O$ vs 0 mm $H_2O$ | 0.009112048 | 0.102864098 |

| Gene | Protein ID | logFC | AveExpr | comparison | pvalue | fdr |
|---|---|---|---|---|---|---|
| ITIH2 | P19823 | -0.444385226 | 21.56628132 | 3 m$H_2O$ vs 0 mm $H_2O$ | 0.009289891 | 0.102864098 |
| CSRP1 | P21291 | 0.123023858 | 18.18444219 | 3 m$H_2O$ vs 0 mm $H_2O$ | 0.009305483 | 0.102864098 |
| ALDOA | P04075 | 0.14476581 | 20.97535175 | 3 m$H_2O$ vs 0 mm $H_2O$ | 0.009511934 | 0.102864098 |
| CS | O75390 | -0.139831151 | 18.86813435 | 3 m$H_2O$ vs 0 mm $H_2O$ | 0.009575937 | 0.102864098 |
| FN1 | P02751 | 0.185701999 | 25.93912475 | 3 m$H_2O$ vs 0 mm $H_2O$ | 0.010047903 | 0.104560993 |
| HSP90AB1 | P08238 | -0.159862642 | 20.68956491 | 3 m$H_2O$ vs 0 mm $H_2O$ | 0.011245681 | 0.109850696 |
| EMILIN1 | Q9Y6C2 | 0.166543931 | 18.74132285 | 3 m$H_2O$ vs 0 mm $H_2O$ | 0.011953282 | 0.109850696 |
| STOM | P27105 | 0.351391044 | 18.22513008 | 3 m$H_2O$ vs 0 mm $H_2O$ | 0.012054837 | 0.109850696 |
| ETF1 | P62495 | -0.222774406 | 16.60031114 | 3 m$H_2O$ vs 0 mm $H_2O$ | 0.012407121 | 0.109850696 |
| PFKL | P17858 | -0.222925291 | 19.53869474 | 3 m$H_2O$ vs 0 mm $H_2O$ | 0.012592304 | 0.109850696 |
| RPS17 | P08708 | 0.16629904 | 19.20866766 | 3 m$H_2O$ vs 0 mm $H_2O$ | 0.012685526 | 0.109850696 |
| EEF2 | P13639 | -0.109307578 | 21.48552521 | 3 m$H_2O$ vs 0 mm $H_2O$ | 0.012865397 | 0.109850696 |
| LMAN1 | P49257 | 0.187883355 | 19.07369945 | 3 m$H_2O$ vs 0 mm $H_2O$ | 0.014075197 | 0.117176011 |
| PHGDH | O43175 | -0.122361146 | 16.66674826 | 3 m$H_2O$ vs 0 mm $H_2O$ | 0.015854023 | 0.124142483 |
| PRDX5 | P30044 | -0.172619611 | 16.63267706 | 3 m$H_2O$ vs 0 mm $H_2O$ | 0.01602267 | 0.124142483 |
| KRT7 | P08729 | -0.143096831 | 19.43973796 | 3 m$H_2O$ vs 0 mm $H_2O$ | 0.016075628 | 0.124142483 |
| ATP2A2 | P16615 | 0.117420456 | 19.13691058 | 3 m$H_2O$ vs 0 mm $H_2O$ | 0.016403211 | 0.124142483 |
| VDAC3 | Q9Y277 | -0.143631303 | 18.97904053 | 3 m$H_2O$ vs 0 mm $H_2O$ | 0.016793494 | 0.124271859 |
| CAD | P27708 | -0.193070608 | 15.11351192 | 3 m$H_2O$ vs 0 mm $H_2O$ | 0.019201027 | 0.136298531 |
| TUBB6 | Q9BUF5 | -0.087607678 | 21.29011268 | 3 m$H_2O$ vs 0 mm $H_2O$ | 0.019382999 | 0.136298531 |
| RPN1 | P04843 | 0.109642817 | 20.58979474 | 3 m$H_2O$ vs 0 mm $H_2O$ | 0.019646635 | 0.136298531 |
| VIM | P08670 | 0.143840901 | 23.56319756 | 3 m$H_2O$ vs 0 mm $H_2O$ | 0.0204391 | 0.138874102 |
| SERPINH1 | P50454 | 0.204038409 | 20.57627706 | 3 m$H_2O$ vs 0 mm $H_2O$ | 0.020912175 | 0.138874102 |
| COL1A2 | P08123 | 0.297162237 | 16.18730792 | 3 m$H_2O$ vs 0 mm $H_2O$ | 0.021446719 | 0.138874102 |
| ALDH18A1 | P54886 | -0.19126924 | 15.2913844 | 3 m$H_2O$ vs 0 mm $H_2O$ | 0.021793012 | 0.138874102 |
| HSPG2 | P98160 | 0.120814343 | 24.74423208 | 3 m$H_2O$ vs 0 mm $H_2O$ | 0.022404182 | 0.138874102 |
| EIF2AK2 | P19525 | -0.125507664 | 15.71603506 | 3 m$H_2O$ vs 0 mm $H_2O$ | 0.022520125 | 0.138874102 |
| RPS19 | P39019 | 0.184493949 | 16.54266348 | 3 m$H_2O$ vs 0 mm $H_2O$ | 0.024676333 | 0.14940398 |
| RPL10A | P62906 | 0.106289465 | 19.96141059 | 3 m$H_2O$ vs 0 mm $H_2O$ | 0.026857129 | 0.159704 |
| LAMC1 | P11047 | 0.148337594 | 21.18750318 | 3 m$H_2O$ vs 0 mm $H_2O$ | 0.027665674 | 0.160609709 |
| CCT4 | P50991 | -0.090158857 | 18.88952591 | 3 m$H_2O$ vs 0 mm $H_2O$ | 0.028978665 | 0.160609709 |
| COL6A2 | P12110 | 0.120648281 | 23.03754647 | 3 m$H_2O$ vs 0 mm $H_2O$ | 0.029086936 | 0.160609709 |
| LAMA4 | Q16363 | 0.10058626 | 21.38083312 | 3 m$H_2O$ vs 0 mm $H_2O$ | 0.029117765 | 0.160609709 |
| RPL13 | P26373 | 0.132839871 | 19.8746121 | 3 m$H_2O$ vs 0 mm $H_2O$ | 0.029420998 | 0.160609709 |
| NID1 | P14543 | 0.118332222 | 20.1895549 | 3 m$H_2O$ vs 0 mm $H_2O$ | 0.031757889 | 0.16867092 |
| POSTN | Q15063 | 0.326562426 | 16.30898796 | 3 m$H_2O$ vs 0 mm $H_2O$ | 0.031910715 | 0.16867092 |
| CALR | P27797 | -0.221103924 | 18.69992242 | 3 m$H_2O$ vs 0 mm $H_2O$ | 0.033172958 | 0.171341812 |
| MMRN2 | Q9H8L6 | 0.190621022 | 19.6353503 | 3 m$H_2O$ vs 0 mm $H_2O$ | 0.033445099 | 0.171341812 |
| ARPC2 | O15144 | -0.12576386 | 18.22115014 | 3 m$H_2O$ vs 0 mm $H_2O$ | 0.03396781 | 0.17138304 |
| COL6A3 | P12111 | 0.116288672 | 25.25110137 | 3 m$H_2O$ vs 0 mm $H_2O$ | 0.035573033 | 0.176803282 |
| BANF1 | O75531 | 0.09802552 | 19.40338113 | 3 m$H_2O$ vs 0 mm $H_2O$ | 0.03730266 | 0.182021868 |

| Gene | Protein ID | logFC | AveExpr | comparison | pvalue | fdr |
|---|---|---|---|---|---|---|
| ACOT7 | O00154 | -0.097919957 | 15.53290167 | 3 m$H_2O$ vs 0 mm $H_2O$ | 0.037716243 | 0.182021868 |
| HSD17B10 | Q99714 | -0.104272406 | 18.62299534 | 3 m$H_2O$ vs 0 mm $H_2O$ | 0.038608214 | 0.183664787 |
| GANAB | Q14697 | -0.116471703 | 19.37791789 | 3 m$H_2O$ vs 0 mm $H_2O$ | 0.042267946 | 0.198242618 |
| RPS13 | P62277 | -0.101006165 | 18.74926133 | 3 m$H_2O$ vs 0 mm $H_2O$ | 0.043536723 | 0.201357344 |
| LRRC59 | Q96AG4 | 0.130592418 | 17.79277083 | 3 m$H_2O$ vs 0 mm $H_2O$ | 0.045237483 | 0.206357285 |
| AHCY | P23526 | 0.117407247 | 18.04335454 | 3 m$H_2O$ vs 0 mm $H_2O$ | 0.048417966 | 0.217068431 |
| NSF | P46459 | -0.129827331 | 14.86161002 | 3 m$H_2O$ vs 0 mm $H_2O$ | 0.048889286 | 0.217068431 |
| EEF1G | P26641 | -0.076894277 | 19.6645041 | 3 m$H_2O$ vs 0 mm $H_2O$ | 0.05009107 | 0.218607039 |
| KIF5B | P33176 | 0.15482228 | 14.94681864 | 3 m$H_2O$ vs 0 mm $H_2O$ | 0.050548775 | 0.218607039 |
| RPL6 | Q02878 | 0.084077867 | 21.67881898 | 3 m$H_2O$ vs 0 mm $H_2O$ | 0.051582209 | 0.220216355 |
| RPS12 | P25398 | -0.171352591 | 16.01206618 | 3 m$H_2O$ vs 0 mm $H_2O$ | 0.053525098 | 0.225618451 |
| MYH9 | P35579 | 0.140092864 | 25.82383901 | 3 m$H_2O$ vs 0 mm $H_2O$ | 0.055816885 | 0.232012321 |
| CAVIN1 | Q6NZI2 | -0.112309221 | 19.48843453 | 3 m$H_2O$ vs 0 mm $H_2O$ | 0.056435429 | 0.232012321 |
| ARPC1B | O15143 | -0.114020392 | 19.59422932 | 3 m$H_2O$ vs 0 mm $H_2O$ | 0.058720466 | 0.238462382 |
| COL12A1 | Q99715 | 0.114739349 | 19.23200628 | 3 m$H_2O$ vs 0 mm $H_2O$ | 0.060538701 | 0.242884188 |
| BAG2 | O95816 | 0.107600201 | 16.49986499 | 3 m$H_2O$ vs 0 mm $H_2O$ | 0.063878838 | 0.253233967 |
| DECR1 | Q16698 | -0.097880346 | 19.03934709 | 3 m$H_2O$ vs 0 mm $H_2O$ | 0.066068577 | 0.258833368 |
| ATP5F1B | P06576 | 0.078441085 | 21.08547363 | 3 m$H_2O$ vs 0 mm $H_2O$ | 0.067914414 | 0.262970929 |
| PHB1 | P35232 | 0.169004833 | 19.87316936 | 3 m$H_2O$ vs 0 mm $H_2O$ | 0.071657997 | 0.267887014 |
| LGALS1 | P09382 | -0.10690805 | 19.93191209 | 3 m$H_2O$ vs 0 mm $H_2O$ | 0.072747543 | 0.267887014 |
| RPL27A | P46776 | 0.472902216 | 18.54518335 | 3 m$H_2O$ vs 0 mm $H_2O$ | 0.072766472 | 0.267887014 |
| H1-0 | P07305 | 0.126866472 | 18.2800493 | 3 m$H_2O$ vs 0 mm $H_2O$ | 0.07288755 | 0.267887014 |
| DSTN | P60981 | -0.099060711 | 18.25003562 | 3 m$H_2O$ vs 0 mm $H_2O$ | 0.073206361 | 0.267887014 |
| TCP1 | P17987 | -0.195761166 | 19.08413477 | 3 m$H_2O$ vs 0 mm $H_2O$ | 0.074631378 | 0.27013314 |
| PSMD13 | Q9UNM6 | -0.079963497 | 18.87776101 | 3 m$H_2O$ vs 0 mm $H_2O$ | 0.077111154 | 0.272419282 |
| DLAT | P10515 | 0.164214981 | 17.13024169 | 3 m$H_2O$ vs 0 mm $H_2O$ | 0.078344744 | 0.272419282 |
| COL4A2 | P08572 | 0.184364095 | 21.58486103 | 3 m$H_2O$ vs 0 mm $H_2O$ | 0.078510513 | 0.272419282 |
| G6PD | P11413 | -0.139546556 | 19.4860753 | 3 m$H_2O$ vs 0 mm $H_2O$ | 0.078964959 | 0.272419282 |
| RPL28 | P46779 | 0.09677449 | 21.24799089 | 3 m$H_2O$ vs 0 mm $H_2O$ | 0.079353364 | 0.272419282 |
| ATP5F1D | P30049 | 0.126373848 | 19.44075806 | 3 m$H_2O$ vs 0 mm $H_2O$ | 0.085271521 | 0.274855736 |
| PCBP1 | Q15365 | -0.069735905 | 18.89923787 | 3 m$H_2O$ vs 0 mm $H_2O$ | 0.085400722 | 0.274855736 |
| TUBB | P07437 | -0.089830013 | 22.27281811 | 3 m$H_2O$ vs 0 mm $H_2O$ | 0.086451428 | 0.274855736 |
| DDX5 | P17844 | -0.081462705 | 18.93492511 | 3 m$H_2O$ vs 0 mm $H_2O$ | 0.087095763 | 0.274855736 |
| EIF4A3 | P38919 | -0.19646764 | 17.58417509 | 3 m$H_2O$ vs 0 mm $H_2O$ | 0.08716353 | 0.274855736 |
| MARCKS | P29966 | 0.163197793 | 16.38719431 | 3 m$H_2O$ vs 0 mm $H_2O$ | 0.087244481 | 0.274855736 |
| HADHA | P40939 | -0.088696684 | 18.98316112 | 3 m$H_2O$ vs 0 mm $H_2O$ | 0.087600595 | 0.274855736 |
| HSPD1 | P10809 | -0.086450301 | 21.18631581 | 3 m$H_2O$ vs 0 mm $H_2O$ | 0.087806138 | 0.274855736 |
| PRXL2A | Q9BRX8 | -0.190339346 | 18.71078175 | 3 m$H_2O$ vs 0 mm $H_2O$ | 0.088052845 | 0.274855736 |
| RPL32 | P62910 | 0.069807439 | 20.21761555 | 3 m$H_2O$ vs 0 mm $H_2O$ | 0.088773601 | 0.274855736 |
| NT5E | P21589 | 0.099734508 | 18.06619473 | 3 m$H_2O$ vs 0 mm $H_2O$ | 0.089142401 | 0.274855736 |
| RPS3A | P61247 | 0.06576206 | 21.14483502 | 3 m$H_2O$ vs 0 mm $H_2O$ | 0.094925554 | 0.290001922 |

| Gene | Protein ID | logFC | AveExpr | comparison | pvalue | fdr |
|---|---|---|---|---|---|---|
| ECI2 | O75521 | -0.095930725 | 16.270599 | 3 m$H_2O$ vs 0 mm $H_2O$ | 0.096804921 | 0.293054897 |
| FLNB | O75369 | -0.080047692 | 20.90348132 | 3 m$H_2O$ vs 0 mm $H_2O$ | 0.099148462 | 0.297445386 |
| CFL1 | P23528 | -0.171430521 | 18.71089364 | 3 m$H_2O$ vs 0 mm $H_2O$ | 0.102830621 | 0.30573747 |
| RPL14 | P50914 | 0.100188608 | 18.62701412 | 3 m$H_2O$ vs 0 mm $H_2O$ | 0.105138342 | 0.309832459 |
| CYB5R1 | Q9UHQ9 | -0.084171707 | 17.76261391 | 3 m$H_2O$ vs 0 mm $H_2O$ | 0.106513707 | 0.311132144 |
| BGN | P21810 | 0.134313252 | 17.72622742 | 3 m$H_2O$ vs 0 mm $H_2O$ | 0.107962493 | 0.312621828 |
| RPL4 | P36578 | 0.057847967 | 20.17098489 | 3 m$H_2O$ vs 0 mm $H_2O$ | 0.110516883 | 0.316369214 |
| HSPE1 | P61604 | -0.102231289 | 17.78317038 | 3 m$H_2O$ vs 0 mm $H_2O$ | 0.111333653 | 0.316369214 |
| PTX3 | P26022 | -0.195767356 | 20.76887638 | 3 m$H_2O$ vs 0 mm $H_2O$ | 0.11232824 | 0.316369214 |
| MACF1 | Q9UPN3 | 0.086950279 | 19.0594487 | 3 m$H_2O$ vs 0 mm $H_2O$ | 0.113056866 | 0.316369214 |
| RPS23 | P62266 | 0.167561031 | 19.05334112 | 3 m$H_2O$ vs 0 mm $H_2O$ | 0.114200585 | 0.316906623 |
| ATP5F1A | P25705 | 0.084692743 | 20.00502696 | 3 m$H_2O$ vs 0 mm $H_2O$ | 0.116949473 | 0.321852681 |
| HNRNPU | Q00839 | -0.113339239 | 19.03295246 | 3 m$H_2O$ vs 0 mm $H_2O$ | 0.118179362 | 0.322571538 |
| SAMM50 | Q9Y512 | -0.115082907 | 14.15564128 | 3 m$H_2O$ vs 0 mm $H_2O$ | 0.119894364 | 0.324592058 |
| TGFBI | Q15582 | 0.078276718 | 22.50578879 | 3 m$H_2O$ vs 0 mm $H_2O$ | 0.123089193 | 0.330554041 |
| RPS18 | P62269 | 0.069783115 | 21.59125597 | 3 m$H_2O$ vs 0 mm $H_2O$ | 0.128167064 | 0.339121201 |
| HSPA8 | P11142 | -0.07672588 | 22.30402347 | 3 m$H_2O$ vs 0 mm $H_2O$ | 0.12831613 | 0.339121201 |
| CCN1 | O00622 | 0.093454388 | 20.29624494 | 3 m$H_2O$ vs 0 mm $H_2O$ | 0.135881506 | 0.356287728 |
| ARPC4 | P59998 | -0.139874083 | 18.84161516 | 3 m$H_2O$ vs 0 mm $H_2O$ | 0.138930968 | 0.361437597 |
| LAMB1 | P07942 | 0.079585543 | 21.8352099 | 3 m$H_2O$ vs 0 mm $H_2O$ | 0.142306745 | 0.362730595 |
| RPS20 | P60866 | 0.120275408 | 19.95181265 | 3 m$H_2O$ vs 0 mm $H_2O$ | 0.143428565 | 0.362730595 |
| COL5A1 | P20908 | 0.087604701 | 19.65823476 | 3 m$H_2O$ vs 0 mm $H_2O$ | 0.143503562 | 0.362730595 |
| COL18A1 | P39060 | 0.083230184 | 21.76400023 | 3 m$H_2O$ vs 0 mm $H_2O$ | 0.143785101 | 0.362730595 |
| STOML2 | Q9UJZ1 | 0.065980931 | 19.8083535 | 3 m$H_2O$ vs 0 mm $H_2O$ | 0.146068348 | 0.365719999 |
| RPS11 | P62280 | 0.086040792 | 20.10277157 | 3 m$H_2O$ vs 0 mm $H_2O$ | 0.148982863 | 0.370144607 |
| TGM2 | P21980 | 0.069944117 | 22.41037841 | 3 m$H_2O$ vs 0 mm $H_2O$ | 0.152158736 | 0.370144607 |
| PSMD12 | O00232 | -0.100697599 | 16.03269934 | 3 m$H_2O$ vs 0 mm $H_2O$ | 0.152211294 | 0.370144607 |
| TMED10 | P49755 | 0.14654482 | 16.93071406 | 3 m$H_2O$ vs 0 mm $H_2O$ | 0.152281715 | 0.370144607 |
| DHCR7 | Q9UBM7 | -0.085587405 | 16.32955993 | 3 m$H_2O$ vs 0 mm $H_2O$ | 0.155717658 | 0.373601969 |
| GAPDH | P04406 | -0.087403286 | 20.98767695 | 3 m$H_2O$ vs 0 mm $H_2O$ | 0.157386416 | 0.373601969 |
| HADHB | P55084 | -0.067885074 | 19.70348331 | 3 m$H_2O$ vs 0 mm $H_2O$ | 0.159206983 | 0.373601969 |
| SHMT2 | P34897 | -0.084672815 | 17.5884267 | 3 m$H_2O$ vs 0 mm $H_2O$ | 0.161220878 | 0.373601969 |
| PAPSS2 | O95340 | 0.095239565 | 19.62764135 | 3 m$H_2O$ vs 0 mm $H_2O$ | 0.161264444 | 0.373601969 |
| FGG | P02679 | -0.093823241 | 25.08264222 | 3 m$H_2O$ vs 0 mm $H_2O$ | 0.161419551 | 0.373601969 |
| PECAM1 | P16284 | 0.259588655 | 15.97467544 | 3 m$H_2O$ vs 0 mm $H_2O$ | 0.161557608 | 0.373601969 |
| SERPINE1 | P05121 | 0.08413256 | 20.70463064 | 3 m$H_2O$ vs 0 mm $H_2O$ | 0.164480924 | 0.377738949 |
| COL6A1 | P12109 | 0.086721019 | 22.09386598 | 3 m$H_2O$ vs 0 mm $H_2O$ | 0.16603296 | 0.378691615 |
| ATP5PD | O75947 | 0.082492701 | 17.79180696 | 3 m$H_2O$ vs 0 mm $H_2O$ | 0.169824749 | 0.384705044 |
| DNAJB4 | Q9UDY4 | 0.071777352 | 17.32499825 | 3 m$H_2O$ vs 0 mm $H_2O$ | 0.173538695 | 0.390462065 |
| HSD17B12 | Q53GQ0 | 0.098902299 | 17.61672755 | 3 m$H_2O$ vs 0 mm $H_2O$ | 0.180004892 | 0.398781231 |
| TUFM | P49411 | -0.057571621 | 19.92690738 | 3 m$H_2O$ vs 0 mm $H_2O$ | 0.180429847 | 0.398781231 |

| Gene | Protein ID | logFC | AveExpr | comparison | pvalue | fdr |
|---|---|---|---|---|---|---|
| CCT5 | P48643 | -0.068287281 | 18.65229805 | 3 m$H_2O$ vs 0 mm $H_2O$ | 0.180828727 | 0.398781231 |
| RUVBL2 | Q9Y230 | -0.070168359 | 18.15268106 | 3 m$H_2O$ vs 0 mm $H_2O$ | 0.182389257 | 0.399576463 |
| KRT9 | P35527 | -0.171053169 | 19.0598067 | 3 m$H_2O$ vs 0 mm $H_2O$ | 0.184159564 | 0.399696626 |
| PLOD1 | Q02809 | -0.088133551 | 18.19084371 | 3 m$H_2O$ vs 0 mm $H_2O$ | 0.18543399 | 0.399696626 |
| GLS | O94925 | -0.0981345 | 15.35759825 | 3 m$H_2O$ vs 0 mm $H_2O$ | 0.186684998 | 0.399696626 |
| ANXA2 | P07355 | -0.114875175 | 21.4349227 | 3 m$H_2O$ vs 0 mm $H_2O$ | 0.187245266 | 0.399696626 |
| SRPX | P78539 | -0.07373863 | 19.32751409 | 3 m$H_2O$ vs 0 mm $H_2O$ | 0.19215709 | 0.406662031 |
| RPL21 | P46778 | 0.062989122 | 19.01435715 | 3 m$H_2O$ vs 0 mm $H_2O$ | 0.193787474 | 0.406662031 |
| ATP6V0D1 | P61421 | 0.092272364 | 17.34164973 | 3 m$H_2O$ vs 0 mm $H_2O$ | 0.194171961 | 0.406662031 |
| NDUFA10 | O95299 | 0.089243708 | 16.62857077 | 3 m$H_2O$ vs 0 mm $H_2O$ | 0.198385104 | 0.412888997 |
| PRSS23 | O95084 | 0.07494852 | 19.72187531 | 3 m$H_2O$ vs 0 mm $H_2O$ | 0.204209462 | 0.419284392 |
| DYNC1H1 | Q14204 | -0.048531839 | 21.33031749 | 3 m$H_2O$ vs 0 mm $H_2O$ | 0.204350689 | 0.419284392 |
| ANXA1 | P04083 | -0.086850456 | 19.10152145 | 3 m$H_2O$ vs 0 mm $H_2O$ | 0.2063985 | 0.419284392 |
| PHB2 | Q99623 | 0.11147002 | 20.07531942 | 3 m$H_2O$ vs 0 mm $H_2O$ | 0.207440878 | 0.419284392 |
| CLU | P10909 | -0.079516612 | 20.64891502 | 3 m$H_2O$ vs 0 mm $H_2O$ | 0.207753528 | 0.419284392 |
| RPS8 | P62241 | -0.072859874 | 20.45506954 | 3 m$H_2O$ vs 0 mm $H_2O$ | 0.219179879 | 0.43968012 |
| MFAP2 | P55001 | -0.130876344 | 18.75574408 | 3 m$H_2O$ vs 0 mm $H_2O$ | 0.221819482 | 0.440909805 |
| EIF4G1 | Q04637 | -0.059679304 | 17.58873533 | 3 m$H_2O$ vs 0 mm $H_2O$ | 0.222440982 | 0.440909805 |
| CAPZA1 | P52907 | -0.103080419 | 16.07243134 | 3 m$H_2O$ vs 0 mm $H_2O$ | 0.225073903 | 0.442501588 |
| ACTN1 | P12814 | -0.048884448 | 21.41145854 | 3 m$H_2O$ vs 0 mm $H_2O$ | 0.225901711 | 0.442501588 |
| XRCC6 | P12956 | -0.088313075 | 17.85962569 | 3 m$H_2O$ vs 0 mm $H_2O$ | 0.227704739 | 0.443425018 |
| LARS1 | Q9P2J5 | -0.086446635 | 16.82750163 | 3 m$H_2O$ vs 0 mm $H_2O$ | 0.231794637 | 0.448765198 |
| H4C16 | P62805 | 0.251227775 | 23.19715251 | 3 m$H_2O$ vs 0 mm $H_2O$ | 0.239192896 | 0.460411759 |
| SPTAN1 | Q13813 | -0.115932134 | 18.24571041 | 3 m$H_2O$ vs 0 mm $H_2O$ | 0.242649309 | 0.464380574 |
| RPL22 | P35268 | -0.0818337 | 19.68733867 | 3 m$H_2O$ vs 0 mm $H_2O$ | 0.247580493 | 0.47111031 |
| NDUFA2 | O43678 | -0.058404696 | 17.93233932 | 3 m$H_2O$ vs 0 mm $H_2O$ | 0.251236172 | 0.475350257 |
| CYB5R3 | P00387 | 0.041025833 | 19.72778346 | 3 m$H_2O$ vs 0 mm $H_2O$ | 0.259335461 | 0.487902307 |
| HTRA3 | P83110 | -0.100865631 | 17.52469239 | 3 m$H_2O$ vs 0 mm $H_2O$ | 0.262338807 | 0.490779902 |
| FLNC | Q14315 | 0.05047834 | 20.94363974 | 3 m$H_2O$ vs 0 mm $H_2O$ | 0.266374125 | 0.495545159 |
| MYH10 | P35580 | 0.072403839 | 23.55126614 | 3 m$H_2O$ vs 0 mm $H_2O$ | 0.274372582 | 0.507589277 |
| MAP1B | P46821 | 0.117762331 | 15.38233639 | 3 m$H_2O$ vs 0 mm $H_2O$ | 0.277080946 | 0.509767707 |
| IQGAP1 | P46940 | -0.066012199 | 18.57483499 | 3 m$H_2O$ vs 0 mm $H_2O$ | 0.285048712 | 0.519563437 |
| CAPZB | P47756 | -0.041437298 | 18.76183816 | 3 m$H_2O$ vs 0 mm $H_2O$ | 0.286234455 | 0.519563437 |
| EIF3E | P60228 | -0.050115949 | 17.00277122 | 3 m$H_2O$ vs 0 mm $H_2O$ | 0.287086104 | 0.519563437 |
| RPL35A | P18077 | 0.04623465 | 19.59741322 | 3 m$H_2O$ vs 0 mm $H_2O$ | 0.293170148 | 0.527706267 |
| VAPA | Q9P0L0 | 0.165252511 | 16.66742819 | 3 m$H_2O$ vs 0 mm $H_2O$ | 0.296993702 | 0.530349147 |
| PDHB | P11177 | -0.070340707 | 16.73766943 | 3 m$H_2O$ vs 0 mm $H_2O$ | 0.297823695 | 0.530349147 |
| ADAMTS4 | O75173 | 0.084035455 | 16.22315078 | 3 m$H_2O$ vs 0 mm $H_2O$ | 0.307436739 | 0.544555501 |
| ATL3 | Q6DD88 | -0.041714503 | 19.24931374 | 3 m$H_2O$ vs 0 mm $H_2O$ | 0.314675681 | 0.549037268 |
| MTCH2 | Q9Y6C9 | 0.063855595 | 19.00153265 | 3 m$H_2O$ vs 0 mm $H_2O$ | 0.315252511 | 0.549037268 |
| MYADM | Q96S97 | 0.060759801 | 17.86105498 | 3 m$H_2O$ vs 0 mm $H_2O$ | 0.316578077 | 0.549037268 |

| Gene | Protein ID | logFC | AveExpr | comparison | pvalue | fdr |
|---|---|---|---|---|---|---|
| ACSL3 | O95573 | 0.043899288 | 18.8979977 | 3 m$H_2O$ vs 0 mm $H_2O$ | 0.31659734 | 0.549037268 |
| FASN | P49327 | -0.044052708 | 19.88841937 | 3 m$H_2O$ vs 0 mm $H_2O$ | 0.318210789 | 0.549037268 |
| RRBP1 | Q9P2E9 | 0.096007839 | 16.81832112 | 3 m$H_2O$ vs 0 mm $H_2O$ | 0.320333114 | 0.549850139 |
| MYL6 | P60660 | 0.083742255 | 21.82604227 | 3 m$H_2O$ vs 0 mm $H_2O$ | 0.324411052 | 0.553994258 |
| TMEM43 | Q9BTV4 | -0.080572931 | 18.65996781 | 3 m$H_2O$ vs 0 mm $H_2O$ | 0.326219267 | 0.554239876 |
| CAV1 | Q03135 | 0.053365194 | 21.4061969 | 3 m$H_2O$ vs 0 mm $H_2O$ | 0.328280279 | 0.55491032 |
| MFGE8 | Q08431 | 0.055222574 | 17.96391857 | 3 m$H_2O$ vs 0 mm $H_2O$ | 0.330976474 | 0.556642251 |
| RPL7 | P18124 | 0.034705041 | 21.07465763 | 3 m$H_2O$ vs 0 mm $H_2O$ | 0.341978405 | 0.572255321 |
| NDUFA13 | Q9P0J0 | -0.065771306 | 17.52794809 | 3 m$H_2O$ vs 0 mm $H_2O$ | 0.344258839 | 0.573190967 |
| COPA | P53621 | -0.047969134 | 18.80077329 | 3 m$H_2O$ vs 0 mm $H_2O$ | 0.347710532 | 0.574906416 |
| KRT10 | P13645 | -0.198923425 | 21.96807875 | 3 m$H_2O$ vs 0 mm $H_2O$ | 0.34874203 | 0.574906416 |
| FGB | P02675 | -0.053129586 | 26.64713393 | 3 m$H_2O$ vs 0 mm $H_2O$ | 0.355536371 | 0.58145344 |
| H1-5 | P16401 | 0.195245905 | 22.04840624 | 3 m$H_2O$ vs 0 mm $H_2O$ | 0.358347788 | 0.58145344 |
| RACK1 | P63244 | 0.038212529 | 20.05788219 | 3 m$H_2O$ vs 0 mm $H_2O$ | 0.359340774 | 0.58145344 |
| RPS4X | P62701 | 0.033993939 | 20.11765589 | 3 m$H_2O$ vs 0 mm $H_2O$ | 0.359697924 | 0.58145344 |
| RPL27 | P61353 | 0.057226296 | 19.40289322 | 3 m$H_2O$ vs 0 mm $H_2O$ | 0.366082702 | 0.588915651 |
| CCT8 | P50990 | -0.049677543 | 19.07398562 | 3 m$H_2O$ vs 0 mm $H_2O$ | 0.36942841 | 0.591440675 |
| VCL | P18206 | 0.095987202 | 16.93515267 | 3 m$H_2O$ vs 0 mm $H_2O$ | 0.374927874 | 0.59737312 |
| TAGLN | Q01995 | 0.063325559 | 18.81489542 | 3 m$H_2O$ vs 0 mm $H_2O$ | 0.386217426 | 0.61243049 |
| TLN1 | Q9Y490 | -0.038385209 | 19.12623259 | 3 m$H_2O$ vs 0 mm $H_2O$ | 0.390413996 | 0.616150999 |
| TMEM33 | P57088 | 0.113871294 | 15.21113383 | 3 m$H_2O$ vs 0 mm $H_2O$ | 0.39362356 | 0.617290694 |
| CCT3 | P49368 | -0.041994034 | 20.32312181 | 3 m$H_2O$ vs 0 mm $H_2O$ | 0.394843597 | 0.617290694 |
| MSN | P26038 | -0.044422739 | 18.77231066 | 3 m$H_2O$ vs 0 mm $H_2O$ | 0.400134805 | 0.622639673 |
| RPL23 | P62829 | 0.029398613 | 19.69082598 | 3 m$H_2O$ vs 0 mm $H_2O$ | 0.403375204 | 0.623050862 |
| PDLIM5 | Q96HC4 | -0.095273431 | 17.8332416 | 3 m$H_2O$ vs 0 mm $H_2O$ | 0.4041411 | 0.623050862 |
| DDB1 | Q16531 | -0.04182991 | 16.99934053 | 3 m$H_2O$ vs 0 mm $H_2O$ | 0.410293809 | 0.625097498 |
| IMPDH2 | P12268 | -0.040763273 | 17.60641556 | 3 m$H_2O$ vs 0 mm $H_2O$ | 0.410689322 | 0.625097498 |
| GARS1 | P41250 | -0.037829993 | 17.42784741 | 3 m$H_2O$ vs 0 mm $H_2O$ | 0.411100157 | 0.625097498 |
| ACTN4 | O43707 | -0.039137368 | 20.91981811 | 3 m$H_2O$ vs 0 mm $H_2O$ | 0.413909823 | 0.62650896 |
| MMRN1 | Q13201 | 0.054219232 | 21.09043115 | 3 m$H_2O$ vs 0 mm $H_2O$ | 0.417202908 | 0.628636056 |
| GNAI2 | P04899 | -0.04020003 | 19.49499447 | 3 m$H_2O$ vs 0 mm $H_2O$ | 0.423924127 | 0.63588619 |
| CLTC | Q00610 | -0.068026836 | 20.30142585 | 3 m$H_2O$ vs 0 mm $H_2O$ | 0.426293724 | 0.63657314 |
| SPTBN1 | Q01082 | -0.04482962 | 18.2872298 | 3 m$H_2O$ vs 0 mm $H_2O$ | 0.447626559 | 0.662148365 |
| SSBP1 | Q04837 | -0.036811508 | 18.41703518 | 3 m$H_2O$ vs 0 mm $H_2O$ | 0.449251212 | 0.662148365 |
| COL3A1 | P02461 | 0.052116702 | 18.87600336 | 3 m$H_2O$ vs 0 mm $H_2O$ | 0.449385978 | 0.662148365 |
| RPS7 | P62081 | 0.058020527 | 18.6908975 | 3 m$H_2O$ vs 0 mm $H_2O$ | 0.451972984 | 0.663026448 |
| HSD17B4 | P51659 | -0.071082946 | 15.898217 | 3 m$H_2O$ vs 0 mm $H_2O$ | 0.457539379 | 0.666761153 |
| TOMM40 | O96008 | 0.080736262 | 16.40672496 | 3 m$H_2O$ vs 0 mm $H_2O$ | 0.458523436 | 0.666761153 |
| PSMD2 | Q13200 | -0.039204052 | 17.68793105 | 3 m$H_2O$ vs 0 mm $H_2O$ | 0.460650477 | 0.666941778 |
| PKM | P14618 | -0.036377207 | 21.11943098 | 3 m$H_2O$ vs 0 mm $H_2O$ | 0.466363268 | 0.672289906 |
| RPL18A | Q02543 | 0.026987199 | 20.57824937 | 3 m$H_2O$ vs 0 mm $H_2O$ | 0.470387167 | 0.675167787 |

| Gene | Protein ID | logFC | AveExpr | comparison | pvalue | fdr |
|---|---|---|---|---|---|---|
| NDUFB10 | O96000 | 0.048272744 | 17.39845643 | 3 m$H_2O$ vs 0 mm $H_2O$ | 0.474380848 | 0.677048993 |
| ACTR3 | P61158 | -0.048882457 | 18.92023602 | 3 m$H_2O$ vs 0 mm $H_2O$ | 0.475764157 | 0.677048993 |
| MACROH2A1 | O75367 | 0.059437373 | 18.22980609 | 3 m$H_2O$ vs 0 mm $H_2O$ | 0.48081804 | 0.681329393 |
| VASP | P50552 | 0.05273767 | 16.99263325 | 3 m$H_2O$ vs 0 mm $H_2O$ | 0.483230784 | 0.681846826 |
| S100A11 | P31949 | 0.031750159 | 19.68174633 | 3 m$H_2O$ vs 0 mm $H_2O$ | 0.495461594 | 0.696154898 |
| EEF1D | P29692 | 0.023535683 | 17.18896539 | 3 m$H_2O$ vs 0 mm $H_2O$ | 0.502158651 | 0.698397297 |
| CTSB | P07858 | -0.042514066 | 18.18104717 | 3 m$H_2O$ vs 0 mm $H_2O$ | 0.50364219 | 0.698397297 |
| FBN1 | P35555 | -0.03982393 | 21.77901541 | 3 m$H_2O$ vs 0 mm $H_2O$ | 0.505351052 | 0.698397297 |
| RPL3 | P39023 | 0.025967118 | 21.79984935 | 3 m$H_2O$ vs 0 mm $H_2O$ | 0.508044273 | 0.698397297 |
| RPL15 | P61313 | 0.039249202 | 19.97908186 | 3 m$H_2O$ vs 0 mm $H_2O$ | 0.514706623 | 0.698397297 |
| RPL34 | P49207 | 0.034988306 | 19.23488626 | 3 m$H_2O$ vs 0 mm $H_2O$ | 0.515412069 | 0.698397297 |
| RPL18 | Q07020 | -0.031663629 | 21.23440702 | 3 m$H_2O$ vs 0 mm $H_2O$ | 0.516495975 | 0.698397297 |
| PXDN | Q92626 | -0.035759396 | 20.11908894 | 3 m$H_2O$ vs 0 mm $H_2O$ | 0.516929766 | 0.698397297 |
| RPS3 | P23396 | -0.023992069 | 21.84558542 | 3 m$H_2O$ vs 0 mm $H_2O$ | 0.516960314 | 0.698397297 |
| SRPX2 | O60687 | -0.057251271 | 17.80297146 | 3 m$H_2O$ vs 0 mm $H_2O$ | 0.519292654 | 0.698397297 |
| HSPA5 | P11021 | -0.039486554 | 21.89509302 | 3 m$H_2O$ vs 0 mm $H_2O$ | 0.521775972 | 0.698397297 |
| FLNA | P21333 | 0.024820672 | 22.91262799 | 3 m$H_2O$ vs 0 mm $H_2O$ | 0.522225006 | 0.698397297 |
| PSMC5 | P62195 | 0.030715959 | 16.40322065 | 3 m$H_2O$ vs 0 mm $H_2O$ | 0.524896854 | 0.699162609 |
| PRDX6 | P30041 | -0.046250745 | 16.40196639 | 3 m$H_2O$ vs 0 mm $H_2O$ | 0.532659283 | 0.706675463 |
| RPS9 | P46781 | -0.024218099 | 21.30621062 | 3 m$H_2O$ vs 0 mm $H_2O$ | 0.539862965 | 0.711082517 |
| LMNB1 | P20700 | -0.041909444 | 20.31075579 | 3 m$H_2O$ vs 0 mm $H_2O$ | 0.540251882 | 0.711082517 |
| RSL1D1 | O76021 | -0.052588587 | 14.99516928 | 3 m$H_2O$ vs 0 mm $H_2O$ | 0.542655548 | 0.711434242 |
| KRT8 | P05787 | -0.129443726 | 22.31188608 | 3 m$H_2O$ vs 0 mm $H_2O$ | 0.545196249 | 0.711507447 |
| HSPB1 | P04792 | -0.028059494 | 20.58486216 | 3 m$H_2O$ vs 0 mm $H_2O$ | 0.546984704 | 0.711507447 |
| RPL19 | P84098 | -0.036585449 | 19.74579175 | 3 m$H_2O$ vs 0 mm $H_2O$ | 0.550961626 | 0.71191025 |
| RAB7A | P51149 | -0.031842291 | 17.51936984 | 3 m$H_2O$ vs 0 mm $H_2O$ | 0.554389478 | 0.71191025 |
| RPS16 | P62249 | 0.033171299 | 21.02085815 | 3 m$H_2O$ vs 0 mm $H_2O$ | 0.554673804 | 0.71191025 |
| PLEC | Q15149 | 0.023833585 | 23.08702195 | 3 m$H_2O$ vs 0 mm $H_2O$ | 0.555845841 | 0.71191025 |
| RPL8 | P62917 | 0.070699057 | 20.07922488 | 3 m$H_2O$ vs 0 mm $H_2O$ | 0.564341729 | 0.720022207 |
| PLOD2 | O00469 | 0.023712655 | 18.32332173 | 3 m$H_2O$ vs 0 mm $H_2O$ | 0.566807894 | 0.720408507 |
| AHNAK | Q09666 | 0.045196224 | 20.16252586 | 3 m$H_2O$ vs 0 mm $H_2O$ | 0.573114628 | 0.725654643 |
| IMMT | Q16891 | 0.031477503 | 20.00559592 | 3 m$H_2O$ vs 0 mm $H_2O$ | 0.579105596 | 0.730462741 |
| DARS1 | P14868 | 0.023248343 | 18.31722998 | 3 m$H_2O$ vs 0 mm $H_2O$ | 0.583646958 | 0.73341297 |
| LMNB2 | Q03252 | -0.03423055 | 20.43563345 | 3 m$H_2O$ vs 0 mm $H_2O$ | 0.587703464 | 0.735734036 |
| COL1A1 | P02452 | 0.022341057 | 19.50614173 | 3 m$H_2O$ vs 0 mm $H_2O$ | 0.594665994 | 0.741662082 |
| RPL23A | P62750 | 0.049525376 | 20.24011738 | 3 m$H_2O$ vs 0 mm $H_2O$ | 0.603502327 | 0.74987416 |
| C5 | P01031 | -0.032877509 | 18.82768108 | 3 m$H_2O$ vs 0 mm $H_2O$ | 0.610472046 | 0.755714466 |
| VWF | P04275 | -0.042706816 | 22.88791985 | 3 m$H_2O$ vs 0 mm $H_2O$ | 0.615922651 | 0.759070131 |
| SEC11A | P67812 | 0.022477537 | 18.655312 | 3 m$H_2O$ vs 0 mm $H_2O$ | 0.618185801 | 0.759070131 |
| RPS2 | P15880 | -0.024370438 | 20.34911658 | 3 m$H_2O$ vs 0 mm $H_2O$ | 0.620600804 | 0.759070131 |
| NID2 | Q14112 | 0.029922435 | 20.71553975 | 3 m$H_2O$ vs 0 mm $H_2O$ | 0.622300738 | 0.759070131 |

| Gene | Protein ID | logFC | AveExpr | comparison | pvalue | fdr |
|---|---|---|---|---|---|---|
| ACLY | P53396 | 0.021450497 | 20.50835424 | 3 m$H_2O$ vs 0 mm $H_2O$ | 0.63067985 | 0.766483176 |
| HTRA1 | Q92743 | 0.041907202 | 19.88510114 | 3 m$H_2O$ vs 0 mm $H_2O$ | 0.63694301 | 0.771280081 |
| UGP2 | Q16851 | -0.021936418 | 16.28042693 | 3 m$H_2O$ vs 0 mm $H_2O$ | 0.639637456 | 0.771736496 |
| VDAC2 | P45880 | 0.01862387 | 21.04702323 | 3 m$H_2O$ vs 0 mm $H_2O$ | 0.646875281 | 0.777619589 |
| EIF4G2 | P78344 | 0.022334919 | 18.67469545 | 3 m$H_2O$ vs 0 mm $H_2O$ | 0.64980445 | 0.777619589 |
| VDAC1 | P21796 | 0.033361945 | 20.43149002 | 3 m$H_2O$ vs 0 mm $H_2O$ | 0.651519115 | 0.777619589 |
| MVP | Q14764 | 0.018842337 | 22.37678295 | 3 m$H_2O$ vs 0 mm $H_2O$ | 0.660805074 | 0.785886035 |
| RPS25 | P62851 | 0.027436995 | 18.6137467 | 3 m$H_2O$ vs 0 mm $H_2O$ | 0.679159045 | 0.804839723 |
| SEC61B | P60468 | -0.024577909 | 17.67294288 | 3 m$H_2O$ vs 0 mm $H_2O$ | 0.684844053 | 0.808698829 |
| RPS15A | P62244 | -0.019086185 | 16.81721761 | 3 m$H_2O$ vs 0 mm $H_2O$ | 0.687941387 | 0.809485801 |
| RPL17 | P18621 | 0.077211846 | 15.43673871 | 3 m$H_2O$ vs 0 mm $H_2O$ | 0.695979311 | 0.816060249 |
| MYO1B | O43795 | -0.063202636 | 15.60251924 | 3 m$H_2O$ vs 0 mm $H_2O$ | 0.699528832 | 0.817344215 |
| RPL30 | P62888 | -0.025929406 | 16.60151038 | 3 m$H_2O$ vs 0 mm $H_2O$ | 0.724289117 | 0.843315651 |
| AARS1 | P49588 | -0.023458117 | 16.12354083 | 3 m$H_2O$ vs 0 mm $H_2O$ | 0.72943637 | 0.844249465 |
| TPM3 | P06753 | -0.035228293 | 19.40677325 | 3 m$H_2O$ vs 0 mm $H_2O$ | 0.7301617 | 0.844249465 |
| NDUFS2 | O75306 | 0.020510631 | 15.97505329 | 3 m$H_2O$ vs 0 mm $H_2O$ | 0.736235525 | 0.847454163 |
| PRDX1 | Q06830 | -0.028171773 | 20.74985169 | 3 m$H_2O$ vs 0 mm $H_2O$ | 0.738023145 | 0.847454163 |
| EIF4A1 | P60842 | -0.013231306 | 21.20872477 | 3 m$H_2O$ vs 0 mm $H_2O$ | 0.759199361 | 0.864931575 |
| SND1 | Q7KZF4 | -0.01617554 | 17.04539574 | 3 m$H_2O$ vs 0 mm $H_2O$ | 0.760511175 | 0.864931575 |
| MOGS | Q13724 | 0.018328826 | 16.31339399 | 3 m$H_2O$ vs 0 mm $H_2O$ | 0.761267556 | 0.864931575 |
| RUVBL1 | Q9Y265 | -0.041739044 | 14.1186164 | 3 m$H_2O$ vs 0 mm $H_2O$ | 0.765868293 | 0.864931575 |
| RPL11 | P62913 | -0.016909091 | 20.19794449 | 3 m$H_2O$ vs 0 mm $H_2O$ | 0.766230675 | 0.864931575 |
| TINAGL1 | Q9GZM7 | -0.01914155 | 19.31518041 | 3 m$H_2O$ vs 0 mm $H_2O$ | 0.77108508 | 0.867470715 |
| KRT18 | P05783 | -0.027079318 | 21.504805 | 3 m$H_2O$ vs 0 mm $H_2O$ | 0.776896611 | 0.871065897 |
| THBS1 | P07996 | 0.012255608 | 22.95722555 | 3 m$H_2O$ vs 0 mm $H_2O$ | 0.782024368 | 0.873872868 |
| CALD1 | Q05682 | 0.019143692 | 19.9361574 | 3 m$H_2O$ vs 0 mm $H_2O$ | 0.785675814 | 0.875016877 |
| GREM1 | O60565 | 0.054816545 | 13.94746047 | 3 m$H_2O$ vs 0 mm $H_2O$ | 0.795296501 | 0.882779116 |
| CCT2 | P78371 | -0.008065723 | 19.69240291 | 3 m$H_2O$ vs 0 mm $H_2O$ | 0.817464991 | 0.904371568 |
| FLOT2 | Q14254 | 0.013526701 | 17.96423756 | 3 m$H_2O$ vs 0 mm $H_2O$ | 0.821034315 | 0.904477208 |
| RPLP2 | P05387 | -0.014523367 | 18.74895696 | 3 m$H_2O$ vs 0 mm $H_2O$ | 0.824520309 | 0.904477208 |
| LETM1 | O95202 | 0.01574963 | 17.41326173 | 3 m$H_2O$ vs 0 mm $H_2O$ | 0.825708922 | 0.904477208 |
| HM13 | Q8TCT9 | -0.017071953 | 17.14646182 | 3 m$H_2O$ vs 0 mm $H_2O$ | 0.830252091 | 0.906471955 |
| GCN1 | Q92616 | -0.014512436 | 16.06870247 | 3 m$H_2O$ vs 0 mm $H_2O$ | 0.839284768 | 0.913339306 |
| LMNA | P02545 | -0.010204513 | 22.81706654 | 3 m$H_2O$ vs 0 mm $H_2O$ | 0.849454822 | 0.921395621 |
| RPL24 | P83731 | 0.019653913 | 15.41834801 | 3 m$H_2O$ vs 0 mm $H_2O$ | 0.854340107 | 0.923685895 |
| GSN | P06396 | -0.02916212 | 18.09615138 | 3 m$H_2O$ vs 0 mm $H_2O$ | 0.858809884 | 0.925513565 |
| ARF4 | P18085 | -0.010748628 | 18.52619066 | 3 m$H_2O$ vs 0 mm $H_2O$ | 0.867215611 | 0.929579305 |
| CHCHD3 | Q9NX63 | -0.007958891 | 20.25383758 | 3 m$H_2O$ vs 0 mm $H_2O$ | 0.868367619 | 0.929579305 |
| STT3A | P46977 | 0.014838498 | 19.16816849 | 3 m$H_2O$ vs 0 mm $H_2O$ | 0.872547158 | 0.929579305 |
| PRDX4 | Q13162 | -0.008185761 | 19.17129669 | 3 m$H_2O$ vs 0 mm $H_2O$ | 0.873748716 | 0.929579305 |
| A2M | P01023 | -0.006607654 | 19.99210831 | 3 m$H_2O$ vs 0 mm $H_2O$ | 0.891538067 | 0.945484638 |

| Gene | Protein ID | logFC | AveExpr | comparison | pvalue | fdr |
|---|---|---|---|---|---|---|
| RPS14 | P62263 | 0.005488145 | 18.62611539 | 3 m$H_2O$ vs 0 mm $H_2O$ | 0.89722862 | 0.948498827 |
| MYO1C | O00159 | 0.006201538 | 19.20310494 | 3 m$H_2O$ vs 0 mm $H_2O$ | 0.901625586 | 0.95013076 |
| LRPPRC | P42704 | -0.007080383 | 19.08842444 | 3 m$H_2O$ vs 0 mm $H_2O$ | 0.904908764 | 0.950582393 |
| C3 | P01024 | -0.008103442 | 18.48488775 | 3 m$H_2O$ vs 0 mm $H_2O$ | 0.912461193 | 0.955215061 |
| VARS1 | P26640 | 0.006779219 | 17.26292586 | 3 m$H_2O$ vs 0 mm $H_2O$ | 0.917661876 | 0.955215061 |
| PSMC2 | P35998 | -0.004310318 | 17.63702096 | 3 m$H_2O$ vs 0 mm $H_2O$ | 0.917924383 | 0.955215061 |
| DDOST | P39656 | -0.005701988 | 18.12022848 | 3 m$H_2O$ vs 0 mm $H_2O$ | 0.931440841 | 0.966261059 |
| IDH2 | P48735 | 0.012978617 | 15.22680513 | 3 m$H_2O$ vs 0 mm $H_2O$ | 0.936985655 | 0.968994482 |
| LTBP1 | Q14766 | 0.00313695 | 19.9748236 | 3 m$H_2O$ vs 0 mm $H_2O$ | 0.961900865 | 0.989388493 |
| CCT6A | P40227 | -0.002663205 | 18.08843594 | 3 m$H_2O$ vs 0 mm $H_2O$ | 0.962976717 | 0.989388493 |
| CANX | P27824 | 0.003627442 | 20.42117326 | 3 m$H_2O$ vs 0 mm $H_2O$ | 0.9656194 | 0.989388493 |
| SLC25A5 | P05141 | 0.001446119 | 21.66997078 | 3 m$H_2O$ vs 0 mm $H_2O$ | 0.975893067 | 0.996847827 |
| HSPA9 | P38646 | 0.002171949 | 20.25499001 | 3 m$H_2O$ vs 0 mm $H_2O$ | 0.983168456 | 0.997519991 |
| RAP1A | P62834 | -0.000919263 | 20.39007357 | 3 m$H_2O$ vs 0 mm $H_2O$ | 0.983335986 | 0.997519991 |
| FBN2 | P35556 | -0.001568103 | 19.4779791 | 3 m$H_2O$ vs 0 mm $H_2O$ | 0.987308653 | 0.997519991 |
| RPS5 | P46782 | 0.000894195 | 21.16789776 | 3 m$H_2O$ vs 0 mm $H_2O$ | 0.990550004 | 0.997519991 |
| FLOT1 | O75955 | -0.000629156 | 18.47467429 | 3 m$H_2O$ vs 0 mm $H_2O$ | 0.991532736 | 0.997519991 |
| RPS6 | P62753 | -0.000315815 | 21.27019737 | 3 m$H_2O$ vs 0 mm $H_2O$ | 0.996881766 | 0.997519991 |
| RAB10 | P61026 | 0.000208406 | 15.89189572 | 3 m$H_2O$ vs 0 mm $H_2O$ | 0.997519991 | 0.997519991 |
| EEF2 | P13639 | -0.150304762 | 21.48552521 | 7 m$H_2O$ vs 0 mm $H_2O$ | 0.002735353 | 0.070367688 |
| PFKL | P17858 | -0.302473215 | 19.53869474 | 7 m$H_2O$ vs 0 mm $H_2O$ | 0.002920708 | 0.070367688 |
| MDH2 | P40926 | -0.227126502 | 18.21421021 | 7 m$H_2O$ vs 0 mm $H_2O$ | 0.003300961 | 0.070367688 |
| CKAP4 | Q07065 | 0.177588665 | 20.5524488 | 7 m$H_2O$ vs 0 mm $H_2O$ | 0.003981329 | 0.070367688 |
| RPS19 | P39019 | 0.270355565 | 16.54266348 | 7 m$H_2O$ vs 0 mm $H_2O$ | 0.004202443 | 0.070367688 |
| AHCY | P23526 | 0.200667902 | 18.04335454 | 7 m$H_2O$ vs 0 mm $H_2O$ | 0.004215043 | 0.070367688 |
| PRKDC | P78527 | -0.139326172 | 19.38685821 | 7 m$H_2O$ vs 0 mm $H_2O$ | 0.004373295 | 0.070367688 |
| NONO | Q15233 | -0.259795141 | 19.26168669 | 7 m$H_2O$ vs 0 mm $H_2O$ | 0.00440175 | 0.070367688 |
| PFKP | Q01813 | -0.15869199 | 20.72340758 | 7 m$H_2O$ vs 0 mm $H_2O$ | 0.004435892 | 0.070367688 |
| VCAN | P13611 | -0.158158955 | 19.33335978 | 7 m$H_2O$ vs 0 mm $H_2O$ | 0.004461221 | 0.070367688 |
| HSP90AB1 | P08238 | -0.197410332 | 20.68956491 | 7 m$H_2O$ vs 0 mm $H_2O$ | 0.0047892 | 0.070367688 |
| ITIH2 | P19823 | -0.529559384 | 21.56628132 | 7 m$H_2O$ vs 0 mm $H_2O$ | 0.004860231 | 0.070367688 |
| SRPX2 | O60687 | -0.30318804 | 17.80297146 | 7 m$H_2O$ vs 0 mm $H_2O$ | 0.006248692 | 0.083734211 |
| HSD17B12 | Q53GQ0 | 0.242348813 | 17.61672755 | 7 m$H_2O$ vs 0 mm $H_2O$ | 0.006623681 | 0.083734211 |
| GAPDH | P04406 | -0.201885074 | 20.98767695 | 7 m$H_2O$ vs 0 mm $H_2O$ | 0.006694127 | 0.083734211 |
| RPL23 | P62829 | 0.117780114 | 19.69082598 | 7 m$H_2O$ vs 0 mm $H_2O$ | 0.00678926 | 0.083734211 |
| LAMA4 | Q16363 | 0.138909178 | 21.38083312 | 7 m$H_2O$ vs 0 mm $H_2O$ | 0.007636023 | 0.090814126 |
| FLNB | O75369 | -0.149161652 | 20.90348132 | 7 m$H_2O$ vs 0 mm $H_2O$ | 0.008817771 | 0.093863744 |
| ADAMTS4 | O75173 | 0.261758592 | 16.22315078 | 7 m$H_2O$ vs 0 mm $H_2O$ | 0.008874865 | 0.093863744 |
| KRT9 | P35527 | -0.4020467 | 19.0598067 | 7 m$H_2O$ vs 0 mm $H_2O$ | 0.009046112 | 0.093863744 |
| ARPC4 | P59998 | -0.291232178 | 18.84161516 | 7 m$H_2O$ vs 0 mm $H_2O$ | 0.009265877 | 0.093863744 |
| FLOT2 | Q14254 | 0.192456588 | 17.96423756 | 7 m$H_2O$ vs 0 mm $H_2O$ | 0.009301812 | 0.093863744 |

| Gene | Protein ID | logFC | AveExpr | comparison | pvalue | fdr |
|---|---|---|---|---|---|---|
| HNRNPU | Q00839 | -0.220789799 | 19.03295246 | 7 m$H_2O$ vs 0 mm $H_2O$ | 0.009644511 | 0.094459474 |
| RPL6 | Q02878 | 0.1264263 | 21.67881898 | 7 m$H_2O$ vs 0 mm $H_2O$ | 0.01024127 | 0.096094935 |
| MFAP2 | P55001 | -0.326157911 | 18.75574408 | 7 m$H_2O$ vs 0 mm $H_2O$ | 0.010791073 | 0.096094935 |
| CYB5R1 | Q9UHQ9 | -0.15471367 | 17.76261391 | 7 m$H_2O$ vs 0 mm $H_2O$ | 0.010949051 | 0.096094935 |
| IKBIP | Q70UQ0 | 0.165485508 | 16.34942842 | 7 m$H_2O$ vs 0 mm $H_2O$ | 0.011000743 | 0.096094935 |
| DLAT | P10515 | 0.270378995 | 17.13024169 | 7 m$H_2O$ vs 0 mm $H_2O$ | 0.011768542 | 0.096094935 |
| CTSB | P07858 | -0.193439681 | 18.18104717 | 7 m$H_2O$ vs 0 mm $H_2O$ | 0.012221361 | 0.096094935 |
| RPS17 | P08708 | 0.179497788 | 19.20866766 | 7 m$H_2O$ vs 0 mm $H_2O$ | 0.012238351 | 0.096094935 |
| COL1A2 | P08123 | 0.354371503 | 16.18730792 | 7 m$H_2O$ vs 0 mm $H_2O$ | 0.012301281 | 0.096094935 |
| TUFM | P49411 | -0.126841512 | 19.92690738 | 7 m$H_2O$ vs 0 mm $H_2O$ | 0.012408655 | 0.096094935 |
| ARPC1B | O15143 | -0.170399591 | 19.59422932 | 7 m$H_2O$ vs 0 mm $H_2O$ | 0.012828617 | 0.097089307 |
| RAN | P62826 | -0.157295144 | 20.33795383 | 7 m$H_2O$ vs 0 mm $H_2O$ | 0.013414873 | 0.098910322 |
| ALDH18A1 | P54886 | -0.223830744 | 15.2913844 | 7 m$H_2O$ vs 0 mm $H_2O$ | 0.013884443 | 0.098910322 |
| PXDN | Q92626 | -0.163960635 | 20.11908894 | 7 m$H_2O$ vs 0 mm $H_2O$ | 0.013960316 | 0.098910322 |
| SLC25A5 | P05141 | -0.138891091 | 21.66997078 | 7 m$H_2O$ vs 0 mm $H_2O$ | 0.016562484 | 0.113943991 |
| ARPC2 | O15144 | -0.156238562 | 18.22115014 | 7 m$H_2O$ vs 0 mm $H_2O$ | 0.016766533 | 0.113943991 |
| THBS1 | P07996 | -0.126073577 | 22.95722555 | 7 m$H_2O$ vs 0 mm $H_2O$ | 0.018173458 | 0.121035232 |
| RPL7A | P62424 | 0.207975716 | 20.68475958 | 7 m$H_2O$ vs 0 mm $H_2O$ | 0.018778776 | 0.122614363 |
| COL12A1 | Q99715 | 0.160248721 | 19.23200628 | 7 m$H_2O$ vs 0 mm $H_2O$ | 0.019150095 | 0.122634264 |
| NDUFA10 | O95299 | 0.187037503 | 16.62857077 | 7 m$H_2O$ vs 0 mm $H_2O$ | 0.020292098 | 0.124476986 |
| SERPINH1 | P50454 | 0.219731716 | 20.57627706 | 7 m$H_2O$ vs 0 mm $H_2O$ | 0.020477906 | 0.124476986 |
| LGALS1 | P09382 | -0.154975664 | 19.93191209 | 7 m$H_2O$ vs 0 mm $H_2O$ | 0.020559262 | 0.124476986 |
| RPL31 | P62899 | 0.133523241 | 20.04161035 | 7 m$H_2O$ vs 0 mm $H_2O$ | 0.022264535 | 0.132394468 |
| MAP1B | P46821 | 0.284773247 | 15.38233639 | 7 m$H_2O$ vs 0 mm $H_2O$ | 0.024098449 | 0.140785676 |
| P4HB | P07237 | -0.146282358 | 19.25928557 | 7 m$H_2O$ vs 0 mm $H_2O$ | 0.025764375 | 0.147923052 |
| ATP2A2 | P16615 | 0.114140792 | 19.13691058 | 7 m$H_2O$ vs 0 mm $H_2O$ | 0.02668841 | 0.150486817 |
| PRDX5 | P30044 | -0.166241905 | 16.63267706 | 7 m$H_2O$ vs 0 mm $H_2O$ | 0.02729912 | 0.150486817 |
| COX4I1 | P13073 | 0.211862954 | 18.4693388 | 7 m$H_2O$ vs 0 mm $H_2O$ | 0.027566654 | 0.150486817 |
| TMED10 | P49755 | 0.254871464 | 16.93071406 | 7 m$H_2O$ vs 0 mm $H_2O$ | 0.028557434 | 0.153381058 |
| SRPX | P78539 | -0.138849266 | 19.32751409 | 7 m$H_2O$ vs 0 mm $H_2O$ | 0.031418721 | 0.164471212 |
| RPS13 | P62277 | -0.116700589 | 18.74926133 | 7 m$H_2O$ vs 0 mm $H_2O$ | 0.031610083 | 0.164471212 |
| CLU | P10909 | -0.151948172 | 20.64891502 | 7 m$H_2O$ vs 0 mm $H_2O$ | 0.03469347 | 0.177737313 |
| A2M | P01023 | 0.11872485 | 19.99210831 | 7 m$H_2O$ vs 0 mm $H_2O$ | 0.036988668 | 0.18277555 |
| RPN1 | P04843 | 0.102299262 | 20.58979474 | 7 m$H_2O$ vs 0 mm $H_2O$ | 0.037547258 | 0.18277555 |
| RPS12 | P25398 | -0.200242335 | 16.01206618 | 7 m$H_2O$ vs 0 mm $H_2O$ | 0.037761429 | 0.18277555 |
| H1-0 | P07305 | 0.161230537 | 18.2800493 | 7 m$H_2O$ vs 0 mm $H_2O$ | 0.037872411 | 0.18277555 |
| DSTN | P60981 | -0.124767037 | 18.25003562 | 7 m$H_2O$ vs 0 mm $H_2O$ | 0.03955697 | 0.188178157 |
| NID1 | P14543 | 0.119966844 | 20.1895549 | 7 m$H_2O$ vs 0 mm $H_2O$ | 0.040514366 | 0.190018083 |
| HSPA9 | P38646 | -0.243192628 | 20.25499001 | 7 m$H_2O$ vs 0 mm $H_2O$ | 0.043197793 | 0.198285636 |
| EEF1G | P26641 | -0.085209245 | 19.6645041 | 7 m$H_2O$ vs 0 mm $H_2O$ | 0.043915269 | 0.198285636 |
| EMILIN1 | Q9Y6C2 | 0.136174546 | 18.74132285 | 7 m$H_2O$ vs 0 mm $H_2O$ | 0.044063475 | 0.198285636 |

| Gene | Protein ID | logFC | AveExpr | comparison | pvalue | fdr |
|---|---|---|---|---|---|---|
| RPL17 | P18621 | 0.450593192 | 15.43673871 | 7 m$H_2O$ vs 0 mm $H_2O$ | 0.049178288 | 0.218351599 |
| SND1 | Q7KZF4 | 0.120296523 | 17.04539574 | 7 m$H_2O$ vs 0 mm $H_2O$ | 0.050585235 | 0.221643201 |
| CS | O75390 | -0.105707637 | 18.86813435 | 7 m$H_2O$ vs 0 mm $H_2O$ | 0.051572208 | 0.223033055 |
| NT5E | P21589 | 0.123570638 | 18.06619473 | 7 m$H_2O$ vs 0 mm $H_2O$ | 0.053670331 | 0.229131027 |
| HNRNPA2B1 | P22626 | -0.171209472 | 18.53624742 | 7 m$H_2O$ vs 0 mm $H_2O$ | 0.055183015 | 0.232606884 |
| UGP2 | Q16851 | 0.101446728 | 16.28042693 | 7 m$H_2O$ vs 0 mm $H_2O$ | 0.059664317 | 0.242603534 |
| SERPINE1 | P05121 | 0.126468506 | 20.70463064 | 7 m$H_2O$ vs 0 mm $H_2O$ | 0.059671195 | 0.242603534 |
| EIF2AK2 | P19525 | -0.107153737 | 15.71603506 | 7 m$H_2O$ vs 0 mm $H_2O$ | 0.05974021 | 0.242603534 |
| TOMM40 | O96008 | 0.232638398 | 16.40672496 | 7 m$H_2O$ vs 0 mm $H_2O$ | 0.061191466 | 0.245503109 |
| EHD2 | Q9NZN4 | -0.108058071 | 17.32935895 | 7 m$H_2O$ vs 0 mm $H_2O$ | 0.062217988 | 0.246649882 |
| ACTN1 | P12814 | -0.083589732 | 21.41145854 | 7 m$H_2O$ vs 0 mm $H_2O$ | 0.063944641 | 0.250512534 |
| LAMB1 | P07942 | 0.110274452 | 21.8352099 | 7 m$H_2O$ vs 0 mm $H_2O$ | 0.064877998 | 0.251213644 |
| POSTN | Q15063 | 0.291984879 | 16.30898796 | 7 m$H_2O$ vs 0 mm $H_2O$ | 0.066241663 | 0.252964001 |
| PRDX4 | Q13162 | -0.108543623 | 19.17129669 | 7 m$H_2O$ vs 0 mm $H_2O$ | 0.066849346 | 0.252964001 |
| COL18A1 | P39060 | 0.114148539 | 21.76400023 | 7 m$H_2O$ vs 0 mm $H_2O$ | 0.068274721 | 0.255454853 |
| IMMT | Q16891 | 0.117339535 | 20.00559592 | 7 m$H_2O$ vs 0 mm $H_2O$ | 0.070003925 | 0.259014521 |
| DNAJB4 | Q9UDY4 | 0.105248943 | 17.32499825 | 7 m$H_2O$ vs 0 mm $H_2O$ | 0.070877335 | 0.259364313 |
| PKM | P14618 | -0.101314315 | 21.11943098 | 7 m$H_2O$ vs 0 mm $H_2O$ | 0.0736197 | 0.266471305 |
| PRDX1 | Q06830 | -0.169159509 | 20.74985169 | 7 m$H_2O$ vs 0 mm $H_2O$ | 0.07844654 | 0.280889222 |
| MYH10 | P35580 | 0.128698406 | 23.55126614 | 7 m$H_2O$ vs 0 mm $H_2O$ | 0.081335112 | 0.288133962 |
| ATP6V0D1 | P61421 | 0.135962692 | 17.34164973 | 7 m$H_2O$ vs 0 mm $H_2O$ | 0.082757304 | 0.290086129 |
| SEC11A | P67812 | -0.087965519 | 18.655312 | 7 m$H_2O$ vs 0 mm $H_2O$ | 0.085600052 | 0.295530857 |
| HSD17B10 | Q99714 | -0.090186766 | 18.62299534 | 7 m$H_2O$ vs 0 mm $H_2O$ | 0.086085565 | 0.295530857 |
| LTBP1 | Q14766 | -0.128015016 | 19.9748236 | 7 m$H_2O$ vs 0 mm $H_2O$ | 0.08718098 | 0.29623741 |
| ACTN4 | O43707 | -0.091434102 | 20.91981811 | 7 m$H_2O$ vs 0 mm $H_2O$ | 0.089274156 | 0.29766496 |
| RPS7 | P62081 | -0.1474962 | 18.6908975 | 7 m$H_2O$ vs 0 mm $H_2O$ | 0.089388877 | 0.29766496 |
| CAPZA1 | P52907 | 0.157309622 | 16.07243134 | 7 m$H_2O$ vs 0 mm $H_2O$ | 0.093284481 | 0.307561705 |
| ETF1 | P62495 | -0.147396022 | 16.60031114 | 7 m$H_2O$ vs 0 mm $H_2O$ | 0.096218206 | 0.309325506 |
| VIM | P08670 | 0.104400867 | 23.56319756 | 7 m$H_2O$ vs 0 mm $H_2O$ | 0.096926609 | 0.309325506 |
| ACOT7 | O00154 | -0.080687011 | 15.53290167 | 7 m$H_2O$ vs 0 mm $H_2O$ | 0.098569553 | 0.309325506 |
| ANXA2 | P07355 | -0.157335654 | 21.4349227 | 7 m$H_2O$ vs 0 mm $H_2O$ | 0.098971733 | 0.309325506 |
| FBN1 | P35555 | -0.110900328 | 21.77901541 | 7 m$H_2O$ vs 0 mm $H_2O$ | 0.09904364 | 0.309325506 |
| ATP5F1B | P06576 | 0.074938174 | 21.08547363 | 7 m$H_2O$ vs 0 mm $H_2O$ | 0.09939288 | 0.309325506 |
| MYH9 | P35579 | 0.125431104 | 25.82383901 | 7 m$H_2O$ vs 0 mm $H_2O$ | 0.102894205 | 0.317257132 |
| CCT4 | P50991 | -0.068197401 | 18.88952591 | 7 m$H_2O$ vs 0 mm $H_2O$ | 0.106642351 | 0.325797273 |
| DDX5 | P17844 | -0.081029591 | 18.93492511 | 7 m$H_2O$ vs 0 mm $H_2O$ | 0.109730445 | 0.332183983 |
| PHB1 | P35232 | 0.157430143 | 19.87316936 | 7 m$H_2O$ vs 0 mm $H_2O$ | 0.112116226 | 0.336348677 |
| FN1 | P02751 | 0.111969793 | 25.93912475 | 7 m$H_2O$ vs 0 mm $H_2O$ | 0.113543466 | 0.336590411 |
| PLOD1 | Q02809 | -0.114503151 | 18.19084371 | 7 m$H_2O$ vs 0 mm $H_2O$ | 0.114218368 | 0.336590411 |
| RPL13 | P26373 | 0.097047326 | 19.8746121 | 7 m$H_2O$ vs 0 mm $H_2O$ | 0.119028611 | 0.345559196 |
| HSPA8 | P11142 | -0.084470809 | 22.30402347 | 7 m$H_2O$ vs 0 mm $H_2O$ | 0.11933726 | 0.345559196 |

| Gene | Protein ID | logFC | AveExpr | comparison | pvalue | fdr |
|---|---|---|---|---|---|---|
| SNRPD3 | P62318 | -0.07742114 | 19.04018698 | 7 m$H_2O$ vs 0 mm $H_2O$ | 0.122975531 | 0.353024585 |
| HSPG2 | P98160 | 0.081795861 | 24.74423208 | 7 m$H_2O$ vs 0 mm $H_2O$ | 0.125289676 | 0.354846877 |
| RPL10A | P62906 | 0.074644575 | 19.96141059 | 7 m$H_2O$ vs 0 mm $H_2O$ | 0.125741536 | 0.354846877 |
| VCL | P18206 | 0.181552941 | 16.93515267 | 7 m$H_2O$ vs 0 mm $H_2O$ | 0.129417052 | 0.362150239 |
| CCT2 | P78371 | -0.059141088 | 19.69240291 | 7 m$H_2O$ vs 0 mm $H_2O$ | 0.131763126 | 0.365642673 |
| RPS14 | P62263 | 0.071660655 | 18.62611539 | 7 m$H_2O$ vs 0 mm $H_2O$ | 0.133249067 | 0.366710242 |
| TGFBI | Q15582 | 0.081080752 | 22.50578879 | 7 m$H_2O$ vs 0 mm $H_2O$ | 0.135190348 | 0.367526 |
| IDH2 | P48735 | -0.274985257 | 15.22680513 | 7 m$H_2O$ vs 0 mm $H_2O$ | 0.135752847 | 0.367526 |
| PCBP1 | Q15365 | -0.063479422 | 18.89923787 | 7 m$H_2O$ vs 0 mm $H_2O$ | 0.137885884 | 0.370290318 |
| RUVBL2 | Q9Y230 | -0.083902279 | 18.15268106 | 7 m$H_2O$ vs 0 mm $H_2O$ | 0.140342605 | 0.373872698 |
| RPL22 | P35268 | -0.113157563 | 19.68733867 | 7 m$H_2O$ vs 0 mm $H_2O$ | 0.142691973 | 0.374250774 |
| GCN1 | Q92616 | -0.11749146 | 16.06870247 | 7 m$H_2O$ vs 0 mm $H_2O$ | 0.142732277 | 0.374250774 |
| HSD17B4 | P51659 | 0.154954138 | 15.898217 | 7 m$H_2O$ vs 0 mm $H_2O$ | 0.14394359 | 0.374478247 |
| XRCC6 | P12956 | -0.114859992 | 17.85962569 | 7 m$H_2O$ vs 0 mm $H_2O$ | 0.148772059 | 0.3840395 |
| EIF4A3 | P38919 | -0.17377562 | 17.58417509 | 7 m$H_2O$ vs 0 mm $H_2O$ | 0.150789116 | 0.38625212 |
| CCT3 | P49368 | -0.077334028 | 20.32312181 | 7 m$H_2O$ vs 0 mm $H_2O$ | 0.154962519 | 0.393912358 |
| STT3A | P46977 | 0.146304506 | 19.16816849 | 7 m$H_2O$ vs 0 mm $H_2O$ | 0.156834832 | 0.395651508 |
| PTX3 | P26022 | -0.183742371 | 20.76887638 | 7 m$H_2O$ vs 0 mm $H_2O$ | 0.159807441 | 0.398033996 |
| MYO1B | O43795 | -0.255395784 | 15.60251924 | 7 m$H_2O$ vs 0 mm $H_2O$ | 0.161288274 | 0.398033996 |
| RPS15A | P62244 | -0.074078456 | 16.81721761 | 7 m$H_2O$ vs 0 mm $H_2O$ | 0.161365134 | 0.398033996 |
| RPL35A | P18077 | 0.066939468 | 19.59741322 | 7 m$H_2O$ vs 0 mm $H_2O$ | 0.163137994 | 0.399448178 |
| RPL11 | P62913 | -0.087314652 | 20.19794449 | 7 m$H_2O$ vs 0 mm $H_2O$ | 0.167924941 | 0.408167923 |
| CALD1 | Q05682 | -0.106500621 | 19.9361574 | 7 m$H_2O$ vs 0 mm $H_2O$ | 0.173879907 | 0.417857867 |
| GREM1 | O60565 | -0.319249958 | 13.94746047 | 7 m$H_2O$ vs 0 mm $H_2O$ | 0.174421152 | 0.417857867 |
| COL6A3 | P12111 | 0.073809618 | 25.25110137 | 7 m$H_2O$ vs 0 mm $H_2O$ | 0.187847538 | 0.429151903 |
| NSF | P46459 | 0.088716958 | 14.86161002 | 7 m$H_2O$ vs 0 mm $H_2O$ | 0.188975365 | 0.429151903 |
| RPS23 | P62266 | 0.146804409 | 19.05334112 | 7 m$H_2O$ vs 0 mm $H_2O$ | 0.189632015 | 0.429151903 |
| LMNA | P02545 | 0.078086004 | 22.81706654 | 7 m$H_2O$ vs 0 mm $H_2O$ | 0.19075365 | 0.429151903 |
| HADHB | P55084 | -0.067231947 | 19.70348331 | 7 m$H_2O$ vs 0 mm $H_2O$ | 0.191035777 | 0.429151903 |
| MARCKS | P29966 | 0.130141398 | 16.38719431 | 7 m$H_2O$ vs 0 mm $H_2O$ | 0.191957083 | 0.429151903 |
| PRSS23 | O95084 | -0.082444363 | 19.72187531 | 7 m$H_2O$ vs 0 mm $H_2O$ | 0.19352176 | 0.429151903 |
| DDB1 | Q16531 | 0.072275622 | 16.99934053 | 7 m$H_2O$ vs 0 mm $H_2O$ | 0.193894405 | 0.429151903 |
| BAG2 | O95816 | 0.077851312 | 16.49986499 | 7 m$H_2O$ vs 0 mm $H_2O$ | 0.194529883 | 0.429151903 |
| COX5B | P10606 | 0.076872208 | 18.89368926 | 7 m$H_2O$ vs 0 mm $H_2O$ | 0.197140552 | 0.429151903 |
| ATP5F1D | P30049 | 0.098808577 | 19.44075806 | 7 m$H_2O$ vs 0 mm $H_2O$ | 0.197234049 | 0.429151903 |
| SEC61B | P60468 | 0.086317113 | 17.67294288 | 7 m$H_2O$ vs 0 mm $H_2O$ | 0.197502581 | 0.429151903 |
| MYO1C | O00159 | 0.071514089 | 19.20310494 | 7 m$H_2O$ vs 0 mm $H_2O$ | 0.198776328 | 0.429151903 |
| TMEM33 | P57088 | 0.186984763 | 15.21113383 | 7 m$H_2O$ vs 0 mm $H_2O$ | 0.199952728 | 0.429151903 |
| HSPD1 | P10809 | -0.067690174 | 21.18631581 | 7 m$H_2O$ vs 0 mm $H_2O$ | 0.200436921 | 0.429151903 |
| IQGAP1 | P46940 | -0.085554887 | 18.57483499 | 7 m$H_2O$ vs 0 mm $H_2O$ | 0.200950706 | 0.429151903 |
| PRXL2A | Q9BRX8 | -0.148615045 | 18.71078175 | 7 m$H_2O$ vs 0 mm $H_2O$ | 0.201994998 | 0.429151903 |

| Gene | Protein ID | logFC | AveExpr | comparison | pvalue | fdr |
|---|---|---|---|---|---|---|
| RPS16 | P62249 | 0.07882234 | 21.02085815 | 7 m$H_2O$ vs 0 mm $H_2O$ | 0.202332879 | 0.429151903 |
| TCP1 | P17987 | 0.142575405 | 19.08413477 | 7 m$H_2O$ vs 0 mm $H_2O$ | 0.210390077 | 0.441233273 |
| RPL8 | P62917 | -0.168897438 | 20.07922488 | 7 m$H_2O$ vs 0 mm $H_2O$ | 0.21067895 | 0.441233273 |
| MACROH2A1 | O75367 | 0.115197584 | 18.22980609 | 7 m$H_2O$ vs 0 mm $H_2O$ | 0.212557596 | 0.442385497 |
| CCN1 | O00622 | 0.082324772 | 20.29624494 | 7 m$H_2O$ vs 0 mm $H_2O$ | 0.214017145 | 0.44265658 |
| RPL18 | Q07020 | -0.066224957 | 21.23440702 | 7 m$H_2O$ vs 0 mm $H_2O$ | 0.216228512 | 0.442734199 |
| RPL19 | P84098 | -0.083265654 | 19.74579175 | 7 m$H_2O$ vs 0 mm $H_2O$ | 0.216713737 | 0.442734199 |
| ATP5F1A | P25705 | 0.068938143 | 20.00502696 | 7 m$H_2O$ vs 0 mm $H_2O$ | 0.224481597 | 0.455807145 |
| ALDOA | P04075 | 0.06387904 | 20.97535175 | 7 m$H_2O$ vs 0 mm $H_2O$ | 0.231794073 | 0.464301621 |
| KRT18 | P05783 | -0.12593207 | 21.504805 | 7 m$H_2O$ vs 0 mm $H_2O$ | 0.232030732 | 0.464301621 |
| TGM2 | P21980 | 0.061557407 | 22.41037841 | 7 m$H_2O$ vs 0 mm $H_2O$ | 0.233724838 | 0.464301621 |
| CALR | P27797 | -0.123924072 | 18.69992242 | 7 m$H_2O$ vs 0 mm $H_2O$ | 0.234242259 | 0.464301621 |
| LRRC59 | Q96AG4 | 0.078149884 | 17.79277083 | 7 m$H_2O$ vs 0 mm $H_2O$ | 0.237609579 | 0.468189288 |
| G6PD | P11413 | -0.096334947 | 19.4860753 | 7 m$H_2O$ vs 0 mm $H_2O$ | 0.241065118 | 0.468507785 |
| CSRP1 | P21291 | 0.052902396 | 18.18444219 | 7 m$H_2O$ vs 0 mm $H_2O$ | 0.241552728 | 0.468507785 |
| ECI2 | O75521 | -0.070404263 | 16.270599 | 7 m$H_2O$ vs 0 mm $H_2O$ | 0.241992009 | 0.468507785 |
| TUBB | P07437 | -0.06307455 | 22.27281811 | 7 m$H_2O$ vs 0 mm $H_2O$ | 0.24576167 | 0.4730557 |
| FASN | P49327 | -0.054839377 | 19.88841937 | 7 m$H_2O$ vs 0 mm $H_2O$ | 0.249805485 | 0.478076015 |
| H4C16 | P62805 | 0.256661006 | 23.19715251 | 7 m$H_2O$ vs 0 mm $H_2O$ | 0.2609539 | 0.494585492 |
| PHGDH | O43175 | -0.055420154 | 16.66674826 | 7 m$H_2O$ vs 0 mm $H_2O$ | 0.261402543 | 0.494585492 |
| RRBP1 | Q9P2E9 | 0.115152821 | 16.81832112 | 7 m$H_2O$ vs 0 mm $H_2O$ | 0.268694488 | 0.505509969 |
| LETM1 | O95202 | 0.086261208 | 17.41326173 | 7 m$H_2O$ vs 0 mm $H_2O$ | 0.272121341 | 0.509080936 |
| COL6A2 | P12110 | 0.060120053 | 23.03754647 | 7 m$H_2O$ vs 0 mm $H_2O$ | 0.274041505 | 0.509809057 |
| ACTR3 | P61158 | -0.079644033 | 18.92023602 | 7 m$H_2O$ vs 0 mm $H_2O$ | 0.284903737 | 0.527071913 |
| COPA | P53621 | -0.058493785 | 18.80077329 | 7 m$H_2O$ vs 0 mm $H_2O$ | 0.288080021 | 0.530003575 |
| CLTC | Q00610 | -0.097951646 | 20.30142585 | 7 m$H_2O$ vs 0 mm $H_2O$ | 0.290268377 | 0.531095438 |
| VWF | P04275 | -0.097039412 | 22.88791985 | 7 m$H_2O$ vs 0 mm $H_2O$ | 0.296176967 | 0.532543242 |
| FBN2 | P35556 | -0.11278738 | 19.4779791 | 7 m$H_2O$ vs 0 mm $H_2O$ | 0.296817887 | 0.532543242 |
| RPS11 | P62280 | 0.06496324 | 20.10277157 | 7 m$H_2O$ vs 0 mm $H_2O$ | 0.299455722 | 0.532543242 |
| PRDX6 | P30041 | -0.083474841 | 16.40196639 | 7 m$H_2O$ vs 0 mm $H_2O$ | 0.300464601 | 0.532543242 |
| TUBB6 | Q9BUF5 | -0.037793211 | 21.29011268 | 7 m$H_2O$ vs 0 mm $H_2O$ | 0.30161828 | 0.532543242 |
| RPS3 | P23396 | -0.041481784 | 21.84558542 | 7 m$H_2O$ vs 0 mm $H_2O$ | 0.302515936 | 0.532543242 |
| PECAM1 | P16284 | 0.200686342 | 15.97467544 | 7 m$H_2O$ vs 0 mm $H_2O$ | 0.303800833 | 0.532543242 |
| LMAN1 | P49257 | 0.075716071 | 19.07369945 | 7 m$H_2O$ vs 0 mm $H_2O$ | 0.30477324 | 0.532543242 |
| RPL28 | P46779 | 0.058094457 | 21.24799089 | 7 m$H_2O$ vs 0 mm $H_2O$ | 0.30545273 | 0.532543242 |
| MACF1 | Q9UPN3 | 0.057889488 | 19.0594487 | 7 m$H_2O$ vs 0 mm $H_2O$ | 0.310239131 | 0.537950534 |
| DARS1 | P14868 | 0.046630891 | 18.31722998 | 7 m$H_2O$ vs 0 mm $H_2O$ | 0.312490757 | 0.537950534 |
| IMPDH2 | P12268 | 0.053943632 | 17.60641556 | 7 m$H_2O$ vs 0 mm $H_2O$ | 0.313400611 | 0.537950534 |
| RAB10 | P61026 | 0.073377813 | 15.89189572 | 7 m$H_2O$ vs 0 mm $H_2O$ | 0.317426015 | 0.541676291 |
| MSN | P26038 | -0.056426314 | 18.77231066 | 7 m$H_2O$ vs 0 mm $H_2O$ | 0.3216992 | 0.541676291 |
| RAP1A | P62834 | -0.047421317 | 20.39007357 | 7 m$H_2O$ vs 0 mm $H_2O$ | 0.324663944 | 0.541676291 |

| Gene | Protein ID | logFC | AveExpr | comparison | pvalue | fdr |
|---|---|---|---|---|---|---|
| VDAC1 | P21796 | 0.079191099 | 20.43149002 | 7 m$H_2O$ vs 0 mm $H_2O$ | 0.325057436 | 0.541676291 |
| VAPA | Q9P0L0 | 0.16673142 | 16.66742819 | 7 m$H_2O$ vs 0 mm $H_2O$ | 0.325284532 | 0.541676291 |
| RPL24 | P83731 | 0.114961648 | 15.41834801 | 7 m$H_2O$ vs 0 mm $H_2O$ | 0.325919315 | 0.541676291 |
| RPS3A | P61247 | 0.039842118 | 21.14483502 | 7 m$H_2O$ vs 0 mm $H_2O$ | 0.326957761 | 0.541676291 |
| S100A11 | P31949 | -0.048913374 | 19.68174633 | 7 m$H_2O$ vs 0 mm $H_2O$ | 0.332199103 | 0.547635155 |
| RPS5 | P46782 | -0.078492002 | 21.16789776 | 7 m$H_2O$ vs 0 mm $H_2O$ | 0.341286134 | 0.557563192 |
| PLEC | Q15149 | 0.041744941 | 23.08702195 | 7 m$H_2O$ vs 0 mm $H_2O$ | 0.341570244 | 0.557563192 |
| CYB5R3 | P00387 | -0.036648368 | 19.72778346 | 7 m$H_2O$ vs 0 mm $H_2O$ | 0.343669199 | 0.558252894 |
| PAPSS2 | O95340 | -0.066887185 | 19.62764135 | 7 m$H_2O$ vs 0 mm $H_2O$ | 0.348092729 | 0.562693586 |
| DHX9 | Q08211 | -0.072633223 | 16.0006753 | 7 m$H_2O$ vs 0 mm $H_2O$ | 0.351908425 | 0.566113554 |
| RPL34 | P49207 | 0.053201626 | 19.23488626 | 7 m$H_2O$ vs 0 mm $H_2O$ | 0.360479324 | 0.576748578 |
| PHB2 | Q99623 | 0.084703762 | 20.07531942 | 7 m$H_2O$ vs 0 mm $H_2O$ | 0.364102444 | 0.576748578 |
| CAVIN1 | Q6NZI2 | -0.053709274 | 19.48843453 | 7 m$H_2O$ vs 0 mm $H_2O$ | 0.366653014 | 0.576748578 |
| ATP5PD | O75947 | 0.056866063 | 17.79180696 | 7 m$H_2O$ vs 0 mm $H_2O$ | 0.366899803 | 0.576748578 |
| KRT8 | P05787 | -0.208826818 | 22.31188608 | 7 m$H_2O$ vs 0 mm $H_2O$ | 0.367179275 | 0.576748578 |
| COL1A1 | P02452 | 0.040559897 | 19.50614173 | 7 m$H_2O$ vs 0 mm $H_2O$ | 0.372730763 | 0.582719925 |
| RPS2 | P15880 | -0.046281829 | 20.34911658 | 7 m$H_2O$ vs 0 mm $H_2O$ | 0.38566633 | 0.600125644 |
| HSP90B1 | P14625 | -0.067668332 | 20.36585658 | 7 m$H_2O$ vs 0 mm $H_2O$ | 0.393358683 | 0.609248565 |
| STOM | P27105 | 0.112827755 | 18.22513008 | 7 m$H_2O$ vs 0 mm $H_2O$ | 0.397003657 | 0.612047305 |
| NDUFS2 | O75306 | 0.055509283 | 15.97505329 | 7 m$H_2O$ vs 0 mm $H_2O$ | 0.40100865 | 0.615372721 |
| KRT10 | P13645 | -0.18968854 | 21.96807875 | 7 m$H_2O$ vs 0 mm $H_2O$ | 0.402986573 | 0.615571233 |
| DHCR7 | Q9UBM7 | -0.051881453 | 16.32955993 | 7 m$H_2O$ vs 0 mm $H_2O$ | 0.409348616 | 0.622423578 |
| C3 | P01024 | 0.065847711 | 18.48488775 | 7 m$H_2O$ vs 0 mm $H_2O$ | 0.411210772 | 0.622423578 |
| ACLY | P53396 | -0.039193745 | 20.50835424 | 7 m$H_2O$ vs 0 mm $H_2O$ | 0.416800523 | 0.626881169 |
| H1-5 | P16401 | 0.183935924 | 22.04840624 | 7 m$H_2O$ vs 0 mm $H_2O$ | 0.417920779 | 0.626881169 |
| TINAGL1 | Q9GZM7 | -0.056293339 | 19.31518041 | 7 m$H_2O$ vs 0 mm $H_2O$ | 0.43002092 | 0.642138862 |
| GNAI2 | P04899 | -0.042307051 | 19.49499447 | 7 m$H_2O$ vs 0 mm $H_2O$ | 0.432386122 | 0.642788298 |
| PDLIM5 | Q96HC4 | -0.095338399 | 17.8332416 | 7 m$H_2O$ vs 0 mm $H_2O$ | 0.435461752 | 0.643390482 |
| ARF4 | P18085 | 0.054264194 | 18.52619066 | 7 m$H_2O$ vs 0 mm $H_2O$ | 0.436655402 | 0.643390482 |
| HSPE1 | P61604 | 0.050854444 | 17.78317038 | 7 m$H_2O$ vs 0 mm $H_2O$ | 0.441818712 | 0.646158902 |
| KIF5B | P33176 | -0.06078917 | 14.94681864 | 7 m$H_2O$ vs 0 mm $H_2O$ | 0.443503198 | 0.646158902 |
| RPS25 | P62851 | 0.054866256 | 18.6137467 | 7 m$H_2O$ vs 0 mm $H_2O$ | 0.444416331 | 0.646158902 |
| HM13 | Q8TCT9 | 0.065498238 | 17.14646182 | 7 m$H_2O$ vs 0 mm $H_2O$ | 0.448041456 | 0.646158902 |
| SPTBN1 | Q01082 | 0.048008355 | 18.2872298 | 7 m$H_2O$ vs 0 mm $H_2O$ | 0.448236355 | 0.646158902 |
| HTRA1 | Q92743 | -0.071981752 | 19.88510114 | 7 m$H_2O$ vs 0 mm $H_2O$ | 0.452832162 | 0.649970302 |
| RPLP2 | P05387 | 0.052648203 | 18.74895696 | 7 m$H_2O$ vs 0 mm $H_2O$ | 0.45815577 | 0.654789148 |
| SPTAN1 | Q13813 | -0.07593602 | 18.24571041 | 7 m$H_2O$ vs 0 mm $H_2O$ | 0.467765578 | 0.663855628 |
| DYNC1H1 | Q14204 | -0.028933361 | 21.33031749 | 7 m$H_2O$ vs 0 mm $H_2O$ | 0.470458226 | 0.663855628 |
| RACK1 | P63244 | 0.032052491 | 20.05788219 | 7 m$H_2O$ vs 0 mm $H_2O$ | 0.470480265 | 0.663855628 |
| EIF4G2 | P78344 | 0.038116362 | 18.67469545 | 7 m$H_2O$ vs 0 mm $H_2O$ | 0.472772528 | 0.664275325 |
| RPL15 | P61313 | -0.046182568 | 19.97908186 | 7 m$H_2O$ vs 0 mm $H_2O$ | 0.475455272 | 0.665237838 |

| Gene | Protein ID | logFC | AveExpr | comparison | pvalue | fdr |
|---|---|---|---|---|---|---|
| HSPA5 | P11021 | -0.045919561 | 21.89509302 | 7 m$H_2O$ vs 0 mm $H_2O$ | 0.487880486 | 0.677141544 |
| ACSL3 | O95573 | -0.032081996 | 18.8979977 | 7 m$H_2O$ vs 0 mm $H_2O$ | 0.490228757 | 0.677141544 |
| VASP | P50552 | -0.055280946 | 16.99263325 | 7 m$H_2O$ vs 0 mm $H_2O$ | 0.492949193 | 0.677141544 |
| FLOT1 | O75955 | 0.043995798 | 18.47467429 | 7 m$H_2O$ vs 0 mm $H_2O$ | 0.493099328 | 0.677141544 |
| MYL6 | P60660 | 0.061667623 | 21.82604227 | 7 m$H_2O$ vs 0 mm $H_2O$ | 0.494130316 | 0.677141544 |
| COL5A1 | P20908 | 0.042021774 | 19.65823476 | 7 m$H_2O$ vs 0 mm $H_2O$ | 0.498030422 | 0.67968906 |
| EIF4A1 | P60842 | -0.030133347 | 21.20872477 | 7 m$H_2O$ vs 0 mm $H_2O$ | 0.517841939 | 0.699774054 |
| SHMT2 | P34897 | 0.04048027 | 17.5884267 | 7 m$H_2O$ vs 0 mm $H_2O$ | 0.518968084 | 0.699774054 |
| HSPB1 | P04792 | -0.032250036 | 20.58486216 | 7 m$H_2O$ vs 0 mm $H_2O$ | 0.519051626 | 0.699774054 |
| RPL18A | Q02543 | -0.025530227 | 20.57824937 | 7 m$H_2O$ vs 0 mm $H_2O$ | 0.523293692 | 0.702648385 |
| TMEM43 | Q9BTV4 | -0.054249899 | 18.65996781 | 7 m$H_2O$ vs 0 mm $H_2O$ | 0.532793217 | 0.712530688 |
| SAMM50 | Q9Y512 | -0.047064634 | 14.15564128 | 7 m$H_2O$ vs 0 mm $H_2O$ | 0.536078369 | 0.714056388 |
| RPL27 | P61353 | 0.041267735 | 19.40289322 | 7 m$H_2O$ vs 0 mm $H_2O$ | 0.539656875 | 0.715959121 |
| RPS8 | P62241 | -0.037661551 | 20.45506954 | 7 m$H_2O$ vs 0 mm $H_2O$ | 0.544242803 | 0.719080573 |
| FLNC | Q14315 | 0.028857922 | 20.94363974 | 7 m$H_2O$ vs 0 mm $H_2O$ | 0.546328483 | 0.719080573 |
| AARS1 | P49588 | 0.043501754 | 16.12354083 | 7 m$H_2O$ vs 0 mm $H_2O$ | 0.551679312 | 0.721782067 |
| VDAC3 | Q9Y277 | -0.034123348 | 18.97904053 | 7 m$H_2O$ vs 0 mm $H_2O$ | 0.552715997 | 0.721782067 |
| LMNB1 | P20700 | -0.042567589 | 20.31075579 | 7 m$H_2O$ vs 0 mm $H_2O$ | 0.561565536 | 0.729718345 |
| BCAP31 | P51572 | 0.060231778 | 19.52713253 | 7 m$H_2O$ vs 0 mm $H_2O$ | 0.56317602 | 0.729718345 |
| LMNB2 | Q03252 | -0.037108654 | 20.43563345 | 7 m$H_2O$ vs 0 mm $H_2O$ | 0.583684789 | 0.743808319 |
| RPS18 | P62269 | 0.025747471 | 21.59125597 | 7 m$H_2O$ vs 0 mm $H_2O$ | 0.585175175 | 0.743808319 |
| RPL27A | P46776 | 0.144424025 | 18.54518335 | 7 m$H_2O$ vs 0 mm $H_2O$ | 0.586996981 | 0.743808319 |
| RSL1D1 | O76021 | -0.050182521 | 14.99516928 | 7 m$H_2O$ vs 0 mm $H_2O$ | 0.587415815 | 0.743808319 |
| RPL30 | P62888 | -0.04284507 | 16.60151038 | 7 m$H_2O$ vs 0 mm $H_2O$ | 0.588074016 | 0.743808319 |
| RPL23A | P62750 | -0.055296049 | 20.24011738 | 7 m$H_2O$ vs 0 mm $H_2O$ | 0.588852518 | 0.743808319 |
| LARS1 | Q9P2J5 | -0.040837856 | 16.82750163 | 7 m$H_2O$ vs 0 mm $H_2O$ | 0.589685875 | 0.743808319 |
| CFL1 | P23528 | -0.056423584 | 18.71089364 | 7 m$H_2O$ vs 0 mm $H_2O$ | 0.598782327 | 0.752195057 |
| GARS1 | P41250 | -0.025609122 | 17.42784741 | 7 m$H_2O$ vs 0 mm $H_2O$ | 0.600852508 | 0.752195057 |
| PDHB | P11177 | 0.036524298 | 16.73766943 | 7 m$H_2O$ vs 0 mm $H_2O$ | 0.608154864 | 0.756599419 |
| GSN | P06396 | -0.090300903 | 18.09615138 | 7 m$H_2O$ vs 0 mm $H_2O$ | 0.609176783 | 0.756599419 |
| C5 | P01031 | 0.035066592 | 18.82768108 | 7 m$H_2O$ vs 0 mm $H_2O$ | 0.612383178 | 0.756599419 |
| HTRA3 | P83110 | 0.047750419 | 17.52469239 | 7 m$H_2O$ vs 0 mm $H_2O$ | 0.613458989 | 0.756599419 |
| CCT6A | P40227 | -0.030127094 | 18.08843594 | 7 m$H_2O$ vs 0 mm $H_2O$ | 0.626039645 | 0.769266427 |
| DDOST | P39656 | 0.034388866 | 18.12022848 | 7 m$H_2O$ vs 0 mm $H_2O$ | 0.630072703 | 0.771375772 |
| CANX | P27824 | -0.043182269 | 20.42117326 | 7 m$H_2O$ vs 0 mm $H_2O$ | 0.633872585 | 0.773185242 |
| LRPPRC | P42704 | -0.029754009 | 19.08842444 | 7 m$H_2O$ vs 0 mm $H_2O$ | 0.641075364 | 0.776562514 |
| MYADM | Q96S97 | 0.029822837 | 17.86105498 | 7 m$H_2O$ vs 0 mm $H_2O$ | 0.64130538 | 0.776562514 |
| NID2 | Q14112 | -0.02986813 | 20.71553975 | 7 m$H_2O$ vs 0 mm $H_2O$ | 0.646368507 | 0.779857656 |
| COL6A1 | P12109 | 0.028680829 | 22.09386598 | 7 m$H_2O$ vs 0 mm $H_2O$ | 0.658004769 | 0.790760264 |
| RAB7A | P51149 | 0.025315625 | 17.51936984 | 7 m$H_2O$ vs 0 mm $H_2O$ | 0.660154214 | 0.790760264 |
| TLN1 | Q9Y490 | -0.020679515 | 19.12623259 | 7 m$H_2O$ vs 0 mm $H_2O$ | 0.662581222 | 0.790822749 |

| Gene | Protein ID | logFC | AveExpr | comparison | pvalue | fdr |
|---|---|---|---|---|---|---|
| PSMC5 | P62195 | 0.022271819 | 16.40322065 | 7 m$H_2O$ vs 0 mm $H_2O$ | 0.665864929 | 0.791903647 |
| RUVBL1 | Q9Y265 | -0.064127417 | 14.1186164 | 7 m$H_2O$ vs 0 mm $H_2O$ | 0.670236254 | 0.793064838 |
| TPM3 | P06753 | 0.046492813 | 19.40677325 | 7 m$H_2O$ vs 0 mm $H_2O$ | 0.671604458 | 0.793064838 |
| AHNAK | Q09666 | 0.034827039 | 20.16252586 | 7 m$H_2O$ vs 0 mm $H_2O$ | 0.684627317 | 0.800111611 |
| COL3A1 | P02461 | -0.029557192 | 18.87600336 | 7 m$H_2O$ vs 0 mm $H_2O$ | 0.686637038 | 0.800111611 |
| CAD | P27708 | -0.031811535 | 15.11351192 | 7 m$H_2O$ vs 0 mm $H_2O$ | 0.687165894 | 0.800111611 |
| RPL32 | P62910 | 0.016731453 | 20.21761555 | 7 m$H_2O$ vs 0 mm $H_2O$ | 0.687182945 | 0.800111611 |
| FGB | P02675 | 0.023923636 | 26.64713393 | 7 m$H_2O$ vs 0 mm $H_2O$ | 0.69390828 | 0.805127029 |
| EIF3E | P60228 | -0.019114418 | 17.00277122 | 7 m$H_2O$ vs 0 mm $H_2O$ | 0.699226204 | 0.808480299 |
| CAV1 | Q03135 | 0.021821379 | 21.4061969 | 7 m$H_2O$ vs 0 mm $H_2O$ | 0.70465897 | 0.811942689 |
| MOGS | Q13724 | 0.023542156 | 16.31339399 | 7 m$H_2O$ vs 0 mm $H_2O$ | 0.716159233 | 0.818441149 |
| FLNA | P21333 | -0.014906885 | 22.91262799 | 7 m$H_2O$ vs 0 mm $H_2O$ | 0.718514777 | 0.818441149 |
| RPL7 | P18124 | 0.013843347 | 21.07465763 | 7 m$H_2O$ vs 0 mm $H_2O$ | 0.719495183 | 0.818441149 |
| RPL3 | P39023 | 0.014978241 | 21.79984935 | 7 m$H_2O$ vs 0 mm $H_2O$ | 0.7201299 | 0.818441149 |
| PSMD2 | Q13200 | 0.018777087 | 17.68793105 | 7 m$H_2O$ vs 0 mm $H_2O$ | 0.739543186 | 0.837645854 |
| NDUFA13 | Q9P0J0 | -0.021623298 | 17.52794809 | 7 m$H_2O$ vs 0 mm $H_2O$ | 0.768141107 | 0.863278006 |
| BGN | P21810 | 0.024899086 | 17.72622742 | 7 m$H_2O$ vs 0 mm $H_2O$ | 0.769766366 | 0.863278006 |
| RPL4 | P36578 | 0.010800978 | 20.17098489 | 7 m$H_2O$ vs 0 mm $H_2O$ | 0.769950654 | 0.863278006 |
| PSMD12 | O00232 | -0.020516921 | 16.03269934 | 7 m$H_2O$ vs 0 mm $H_2O$ | 0.776903981 | 0.868151092 |
| NDUFB10 | O96000 | -0.019969339 | 17.39845643 | 7 m$H_2O$ vs 0 mm $H_2O$ | 0.780685413 | 0.869459005 |
| FGG | P02679 | -0.018192528 | 25.08264222 | 7 m$H_2O$ vs 0 mm $H_2O$ | 0.792314215 | 0.879468779 |
| VARS1 | P26640 | -0.018181367 | 17.26292586 | 7 m$H_2O$ vs 0 mm $H_2O$ | 0.796216486 | 0.880864086 |
| CHCHD3 | Q9NX63 | 0.012156642 | 20.25383758 | 7 m$H_2O$ vs 0 mm $H_2O$ | 0.81349291 | 0.894453529 |
| ANXA1 | P04083 | -0.016810682 | 19.10152145 | 7 m$H_2O$ vs 0 mm $H_2O$ | 0.81387213 | 0.894453529 |
| VDAC2 | P45880 | -0.009762302 | 21.04702323 | 7 m$H_2O$ vs 0 mm $H_2O$ | 0.822247345 | 0.897933689 |
| ATL3 | Q6DD88 | -0.00978926 | 19.24931374 | 7 m$H_2O$ vs 0 mm $H_2O$ | 0.822431757 | 0.897933689 |
| MVP | Q14764 | -0.009653889 | 22.37678295 | 7 m$H_2O$ vs 0 mm $H_2O$ | 0.833446554 | 0.906985955 |
| RPL14 | P50914 | 0.01261894 | 18.62701412 | 7 m$H_2O$ vs 0 mm $H_2O$ | 0.840832826 | 0.910266217 |
| CAPZB | P47756 | -0.008124351 | 18.76183816 | 7 m$H_2O$ vs 0 mm $H_2O$ | 0.841927913 | 0.910266217 |
| RPL21 | P46778 | -0.009649111 | 19.01435715 | 7 m$H_2O$ vs 0 mm $H_2O$ | 0.847690024 | 0.913530026 |
| MFGE8 | Q08431 | 0.011164922 | 17.96391857 | 7 m$H_2O$ vs 0 mm $H_2O$ | 0.851860711 | 0.91506328 |
| GLS | O94925 | -0.012685227 | 15.35759825 | 7 m$H_2O$ vs 0 mm $H_2O$ | 0.869013384 | 0.930487 |
| DECR1 | Q16698 | -0.00840332 | 19.03934709 | 7 m$H_2O$ vs 0 mm $H_2O$ | 0.87460121 | 0.931370561 |
| CCT8 | P50990 | -0.009156478 | 19.07398562 | 7 m$H_2O$ vs 0 mm $H_2O$ | 0.87543239 | 0.931370561 |
| TMED2 | Q15363 | -0.008666824 | 17.21365208 | 7 m$H_2O$ vs 0 mm $H_2O$ | 0.889721297 | 0.943557936 |
| PSMC2 | P35998 | 0.005855821 | 17.63702096 | 7 m$H_2O$ vs 0 mm $H_2O$ | 0.896146043 | 0.946636472 |
| RPS6 | P62753 | 0.011078412 | 21.27019737 | 7 m$H_2O$ vs 0 mm $H_2O$ | 0.898309685 | 0.946636472 |
| NDUFA2 | O43678 | -0.005586587 | 17.93233932 | 7 m$H_2O$ vs 0 mm $H_2O$ | 0.91627267 | 0.960626221 |
| RPS4X | P62701 | 0.003977126 | 20.11765589 | 7 m$H_2O$ vs 0 mm $H_2O$ | 0.919040067 | 0.960626221 |
| HADHA | P40939 | -0.005248082 | 18.98316112 | 7 m$H_2O$ vs 0 mm $H_2O$ | 0.920239533 | 0.960626221 |
| BANF1 | O75531 | 0.003201781 | 19.40338113 | 7 m$H_2O$ vs 0 mm $H_2O$ | 0.944615406 | 0.968884674 |

| Gene | Protein ID | logFC | AveExpr | comparison | pvalue | fdr |
|---|---|---|---|---|---|---|
| STOML2 | Q9UJZ1 | -0.003202801 | 19.8083535 | 7 m$H_2O$ vs 0 mm $H_2O$ | 0.945203321 | 0.968884674 |
| PSMD13 | Q9UNM6 | -0.003001731 | 18.87776101 | 7 m$H_2O$ vs 0 mm $H_2O$ | 0.947348124 | 0.968884674 |
| CCT5 | P48643 | -0.003435356 | 18.65229805 | 7 m$H_2O$ vs 0 mm $H_2O$ | 0.948063316 | 0.968884674 |
| MMRN1 | Q13201 | 0.004504648 | 21.09043115 | 7 m$H_2O$ vs 0 mm $H_2O$ | 0.94920687 | 0.968884674 |
| EEF1D | P29692 | 0.002369798 | 17.18896539 | 7 m$H_2O$ vs 0 mm $H_2O$ | 0.949296362 | 0.968884674 |
| EIF4G1 | Q04637 | -0.003228963 | 17.58873533 | 7 m$H_2O$ vs 0 mm $H_2O$ | 0.949392059 | 0.968884674 |
| TAGLN | Q01995 | 0.00470134 | 18.81489542 | 7 m$H_2O$ vs 0 mm $H_2O$ | 0.951427293 | 0.968884674 |
| COL4A2 | P08572 | -0.00520382 | 21.58486103 | 7 m$H_2O$ vs 0 mm $H_2O$ | 0.960606905 | 0.975250303 |
| PLOD2 | O00469 | -0.001389644 | 18.32332173 | 7 m$H_2O$ vs 0 mm $H_2O$ | 0.974862514 | 0.984944164 |
| RPS20 | P60866 | 0.002529551 | 19.95181265 | 7 m$H_2O$ vs 0 mm $H_2O$ | 0.976070793 | 0.984944164 |
| RPS9 | P46781 | 0.000665791 | 21.30621062 | 7 m$H_2O$ vs 0 mm $H_2O$ | 0.987364711 | 0.992766153 |
| MTCH2 | Q9Y6C9 | 0.000855092 | 19.00153265 | 7 m$H_2O$ vs 0 mm $H_2O$ | 0.989784873 | 0.992766153 |
| SSBP1 | Q04837 | 0.000135109 | 18.41703518 | 7 m$H_2O$ vs 0 mm $H_2O$ | 0.997909299 | 0.997909299 |
| TMED2 | Q15363 | -0.238724209 | 17.21365208 | 7 m$H_2O$ vs 3 mm $H_2O$ | 0.001929567 | 0.472425384 |
| F2 | P00734 | 0.255529375 | 16.10657822 | 7 m$H_2O$ vs 3 mm $H_2O$ | 0.00283739 | 0.472425384 |
| NSF | P46459 | 0.218544289 | 14.86161002 | 7 m$H_2O$ vs 3 mm $H_2O$ | 0.004730728 | 0.493548204 |
| TCP1 | P17987 | 0.338336571 | 19.08413477 | 7 m$H_2O$ vs 3 mm $H_2O$ | 0.008191362 | 0.493548204 |
| CAPZA1 | P52907 | 0.260390042 | 16.07243134 | 7 m$H_2O$ vs 3 mm $H_2O$ | 0.01042397 | 0.493548204 |
| THBS1 | P07996 | -0.138329185 | 22.95722555 | 7 m$H_2O$ vs 3 mm $H_2O$ | 0.011027735 | 0.493548204 |
| FLOT2 | Q14254 | 0.178929887 | 17.96423756 | 7 m$H_2O$ vs 3 mm $H_2O$ | 0.014004026 | 0.493548204 |
| LAMC1 | P11047 | 0.181885716 | 21.18750318 | 7 m$H_2O$ vs 3 mm $H_2O$ | 0.014129636 | 0.493548204 |
| KIF5B | P33176 | -0.21561145 | 14.94681864 | 7 m$H_2O$ vs 3 mm $H_2O$ | 0.015201305 | 0.493548204 |
| SLC25A5 | P05141 | -0.140337211 | 21.66997078 | 7 m$H_2O$ vs 3 mm $H_2O$ | 0.015685758 | 0.493548204 |
| SRPX2 | O60687 | -0.245936769 | 17.80297146 | 7 m$H_2O$ vs 3 mm $H_2O$ | 0.020191761 | 0.493548204 |
| PRSS23 | O95084 | -0.157392883 | 19.72187531 | 7 m$H_2O$ vs 3 mm $H_2O$ | 0.021516237 | 0.493548204 |
| RPS7 | P62081 | -0.205516726 | 18.6908975 | 7 m$H_2O$ vs 3 mm $H_2O$ | 0.024004774 | 0.493548204 |
| BCAP31 | P51572 | -0.258599885 | 19.52713253 | 7 m$H_2O$ vs 3 mm $H_2O$ | 0.024661469 | 0.493548204 |
| UGP2 | Q16851 | 0.123383146 | 16.28042693 | 7 m$H_2O$ vs 3 mm $H_2O$ | 0.026237353 | 0.493548204 |
| DHX9 | Q08211 | 0.189039744 | 16.0006753 | 7 m$H_2O$ vs 3 mm $H_2O$ | 0.026259201 | 0.493548204 |
| SNRPD3 | P62318 | 0.11609614 | 19.04018698 | 7 m$H_2O$ vs 3 mm $H_2O$ | 0.028145235 | 0.493548204 |
| A2M | P01023 | 0.125332504 | 19.99210831 | 7 m$H_2O$ vs 3 mm $H_2O$ | 0.029078238 | 0.493548204 |
| HSP90B1 | P14625 | 0.187781578 | 20.36585658 | 7 m$H_2O$ vs 3 mm $H_2O$ | 0.029550328 | 0.493548204 |
| SND1 | Q7KZF4 | 0.136472064 | 17.04539574 | 7 m$H_2O$ vs 3 mm $H_2O$ | 0.029642535 | 0.493548204 |
| RPL23 | P62829 | 0.088381501 | 19.69082598 | 7 m$H_2O$ vs 3 mm $H_2O$ | 0.031159014 | 0.494092937 |
| HSPE1 | P61604 | 0.153085733 | 17.78317038 | 7 m$H_2O$ vs 3 mm $H_2O$ | 0.033156615 | 0.501870578 |
| PAPSS2 | O95340 | -0.16212675 | 19.62764135 | 7 m$H_2O$ vs 3 mm $H_2O$ | 0.034937992 | 0.503855169 |
| SEC11A | P67812 | -0.110443056 | 18.655312 | 7 m$H_2O$ vs 3 mm $H_2O$ | 0.036313886 | 0.503855169 |
| CTSB | P07858 | -0.150925615 | 18.18104717 | 7 m$H_2O$ vs 3 mm $H_2O$ | 0.040726981 | 0.512429322 |
| HSD17B4 | P51659 | 0.226037084 | 15.898217 | 7 m$H_2O$ vs 3 mm $H_2O$ | 0.041209891 | 0.512429322 |
| HSPA9 | P38646 | -0.245364577 | 20.25499001 | 7 m$H_2O$ vs 3 mm $H_2O$ | 0.041638159 | 0.512429322 |
| FSCN1 | Q16658 | -0.137282775 | 17.22349762 | 7 m$H_2O$ vs 3 mm $H_2O$ | 0.043521077 | 0.512429322 |

| Gene | Protein ID | logFC | AveExpr | comparison | pvalue | fdr |
|---|---|---|---|---|---|---|
| PXDN | Q92626 | -0.128201239 | 20.11908894 | 7 $mH_2O$ vs 3 mm $H_2O$ | 0.044625977 | 0.512429322 |
| DDB1 | Q16531 | 0.114105531 | 16.99934053 | 7 $mH_2O$ vs 3 mm $H_2O$ | 0.050034451 | 0.55538241 |
| BANF1 | O75531 | -0.094823738 | 19.40338113 | 7 $mH_2O$ vs 3 mm $H_2O$ | 0.056528361 | 0.567351605 |
| ADAMTS4 | O75173 | 0.177723137 | 16.22315078 | 7 $mH_2O$ vs 3 mm $H_2O$ | 0.056660718 | 0.567351605 |
| CAD | P27708 | 0.161259073 | 15.11351192 | 7 $mH_2O$ vs 3 mm $H_2O$ | 0.057518582 | 0.567351605 |
| CYB5R3 | P00387 | -0.077674201 | 19.72778346 | 7 $mH_2O$ vs 3 mm $H_2O$ | 0.057927792 | 0.567351605 |
| SHMT2 | P34897 | 0.125153085 | 17.5884267 | 7 $mH_2O$ vs 3 mm $H_2O$ | 0.061567232 | 0.585768233 |
| GANAB | Q14697 | -0.110718447 | 19.37791789 | 7 $mH_2O$ vs 3 mm $H_2O$ | 0.067194736 | 0.62155131 |
| VDAC3 | Q9Y277 | 0.109507956 | 18.97904053 | 7 $mH_2O$ vs 3 mm $H_2O$ | 0.07274671 | 0.652144611 |
| MMRN2 | Q9H8L6 | 0.165713189 | 19.6353503 | 7 $mH_2O$ vs 3 mm $H_2O$ | 0.075819205 | 0.652144611 |
| HSD17B12 | Q53GQ0 | 0.143446514 | 17.61672755 | 7 $mH_2O$ vs 3 mm $H_2O$ | 0.077764186 | 0.652144611 |
| LTBP1 | Q14766 | -0.131151966 | 19.9748236 | 7 $mH_2O$ vs 3 mm $H_2O$ | 0.080533057 | 0.652144611 |
| RPL8 | P62917 | -0.239596495 | 20.07922488 | 7 $mH_2O$ vs 3 mm $H_2O$ | 0.084719998 | 0.652144611 |
| PRDX4 | Q13162 | -0.100357862 | 19.17129669 | 7 $mH_2O$ vs 3 mm $H_2O$ | 0.087230917 | 0.652144611 |
| STOM | P27105 | -0.238563289 | 18.22513008 | 7 $mH_2O$ vs 3 mm $H_2O$ | 0.087251107 | 0.652144611 |
| IMPDH2 | P12268 | 0.094706904 | 17.60641556 | 7 $mH_2O$ vs 3 mm $H_2O$ | 0.088834208 | 0.652144611 |
| GAPDH | P04406 | -0.114481788 | 20.98767695 | 7 $mH_2O$ vs 3 mm $H_2O$ | 0.090000696 | 0.652144611 |
| COL4A2 | P08572 | -0.189567914 | 21.58486103 | 7 $mH_2O$ vs 3 mm $H_2O$ | 0.090086042 | 0.652144611 |
| RPL17 | P18621 | 0.373381346 | 15.43673871 | 7 $mH_2O$ vs 3 mm $H_2O$ | 0.095281122 | 0.675076887 |
| MFAP2 | P55001 | -0.195281567 | 18.75574408 | 7 $mH_2O$ vs 3 mm $H_2O$ | 0.097748538 | 0.678130482 |
| KRT9 | P35527 | -0.230993532 | 19.0598067 | 7 $mH_2O$ vs 3 mm $H_2O$ | 0.100978106 | 0.686238963 |
| SEC61B | P60468 | 0.110895022 | 17.67294288 | 7 $mH_2O$ vs 3 mm $H_2O$ | 0.104755189 | 0.697669559 |
| PSMD13 | Q9UNM6 | 0.076961766 | 18.87776101 | 7 $mH_2O$ vs 3 mm $H_2O$ | 0.108480488 | 0.70563824 |
| DECR1 | Q16698 | 0.089477026 | 19.03934709 | 7 $mH_2O$ vs 3 mm $H_2O$ | 0.110764652 | 0.70563824 |
| CALD1 | Q05682 | -0.125644313 | 19.9361574 | 7 $mH_2O$ vs 3 mm $H_2O$ | 0.113567565 | 0.70563824 |
| GREM1 | O60565 | -0.374066503 | 13.94746047 | 7 $mH_2O$ vs 3 mm $H_2O$ | 0.116261551 | 0.70563824 |
| ACSL3 | O95573 | -0.075981284 | 18.8979977 | 7 $mH_2O$ vs 3 mm $H_2O$ | 0.116890965 | 0.70563824 |
| IDH2 | P48735 | -0.287963875 | 15.22680513 | 7 $mH_2O$ vs 3 mm $H_2O$ | 0.119755072 | 0.70563824 |
| S100A11 | P31949 | -0.080663533 | 19.68174633 | 7 $mH_2O$ vs 3 mm $H_2O$ | 0.120910981 | 0.70563824 |
| CSRP1 | P21291 | -0.070121462 | 18.18444219 | 7 $mH_2O$ vs 3 mm $H_2O$ | 0.128029705 | 0.70563824 |
| HADHA | P40939 | 0.083448602 | 18.98316112 | 7 $mH_2O$ vs 3 mm $H_2O$ | 0.128967729 | 0.70563824 |
| HTRA3 | P83110 | 0.14861605 | 17.52469239 | 7 $mH_2O$ vs 3 mm $H_2O$ | 0.131640488 | 0.70563824 |
| PRDX1 | Q06830 | -0.140987736 | 20.74985169 | 7 $mH_2O$ vs 3 mm $H_2O$ | 0.135232444 | 0.70563824 |
| ARPC4 | P59998 | -0.151358095 | 18.84161516 | 7 $mH_2O$ vs 3 mm $H_2O$ | 0.135684732 | 0.70563824 |
| TUFM | P49411 | -0.06926989 | 19.92690738 | 7 $mH_2O$ vs 3 mm $H_2O$ | 0.136354003 | 0.70563824 |
| ALDOA | P04075 | -0.08088677 | 20.97535175 | 7 $mH_2O$ vs 3 mm $H_2O$ | 0.136393865 | 0.70563824 |
| LMAN1 | P49257 | -0.112167283 | 19.07369945 | 7 $mH_2O$ vs 3 mm $H_2O$ | 0.137737194 | 0.70563824 |
| LMNA | P02545 | 0.088290517 | 22.81706654 | 7 $mH_2O$ vs 3 mm $H_2O$ | 0.142687615 | 0.719923877 |
| COX5B | P10606 | -0.086990539 | 18.89368926 | 7 $mH_2O$ vs 3 mm $H_2O$ | 0.148047506 | 0.727780413 |
| PDHB | P11177 | 0.106865005 | 16.73766943 | 7 $mH_2O$ vs 3 mm $H_2O$ | 0.14861582 | 0.727780413 |
| STOML2 | Q9UJZ1 | -0.069183732 | 19.8083535 | 7 $mH_2O$ vs 3 mm $H_2O$ | 0.154503593 | 0.730165139 |

| Gene | Protein ID | logFC | AveExpr | comparison | pvalue | fdr |
|---|---|---|---|---|---|---|
| SPTBN1 | Q01082 | 0.092837975 | 18.2872298 | 7 m$H_2O$ vs 3 mm $H_2O$ | 0.154738148 | 0.730165139 |
| MAP1B | P46821 | 0.167010917 | 15.38233639 | 7 m$H_2O$ vs 3 mm $H_2O$ | 0.157607853 | 0.730165139 |
| KRT7 | P08729 | -0.082605636 | 19.43973796 | 7 m$H_2O$ vs 3 mm $H_2O$ | 0.159587941 | 0.730165139 |
| HNRNPU | Q00839 | -0.10745056 | 19.03295246 | 7 m$H_2O$ vs 3 mm $H_2O$ | 0.162805634 | 0.730165139 |
| RPS14 | P62263 | 0.06617251 | 18.62611539 | 7 m$H_2O$ vs 3 mm $H_2O$ | 0.162897494 | 0.730165139 |
| RPL21 | P46778 | -0.072638233 | 19.01435715 | 7 m$H_2O$ vs 3 mm $H_2O$ | 0.164451608 | 0.730165139 |
| IMMT | Q16891 | 0.085862032 | 20.00559592 | 7 m$H_2O$ vs 3 mm $H_2O$ | 0.171909156 | 0.730685553 |
| AHCY | P23526 | 0.083260655 | 18.04335454 | 7 m$H_2O$ vs 3 mm $H_2O$ | 0.172940034 | 0.730685553 |
| FLNB | O75369 | -0.06911396 | 20.90348132 | 7 m$H_2O$ vs 3 mm $H_2O$ | 0.176140373 | 0.730685553 |
| RPS20 | P60866 | -0.117745857 | 19.95181265 | 7 m$H_2O$ vs 3 mm $H_2O$ | 0.178621971 | 0.730685553 |
| RPL14 | P50914 | -0.087569669 | 18.62701412 | 7 m$H_2O$ vs 3 mm $H_2O$ | 0.179040502 | 0.730685553 |
| PHGDH | O43175 | 0.066940992 | 16.66674826 | 7 m$H_2O$ vs 3 mm $H_2O$ | 0.179782445 | 0.730685553 |
| TUBB6 | Q9BUF5 | 0.049814467 | 21.29011268 | 7 m$H_2O$ vs 3 mm $H_2O$ | 0.179928575 | 0.730685553 |
| CCT2 | P78371 | -0.051075365 | 19.69240291 | 7 m$H_2O$ vs 3 mm $H_2O$ | 0.188034468 | 0.742084853 |
| NDUFA10 | O95299 | 0.097793795 | 16.62857077 | 7 m$H_2O$ vs 3 mm $H_2O$ | 0.18938764 | 0.742084853 |
| VASP | P50552 | -0.108018616 | 16.99263325 | 7 m$H_2O$ vs 3 mm $H_2O$ | 0.191600144 | 0.742084853 |
| GCN1 | Q92616 | -0.102979024 | 16.06870247 | 7 m$H_2O$ vs 3 mm $H_2O$ | 0.194550975 | 0.742084853 |
| RPL15 | P61313 | -0.08543177 | 19.97908186 | 7 m$H_2O$ vs 3 mm $H_2O$ | 0.197402735 | 0.742084853 |
| CYB5R1 | Q9UHQ9 | -0.070541962 | 17.76261391 | 7 m$H_2O$ vs 3 mm $H_2O$ | 0.198164339 | 0.742084853 |
| STT3A | P46977 | 0.131466008 | 19.16816849 | 7 m$H_2O$ vs 3 mm $H_2O$ | 0.199893313 | 0.742084853 |
| RPL18A | Q02543 | -0.052517425 | 20.57824937 | 7 m$H_2O$ vs 3 mm $H_2O$ | 0.200563474 | 0.742084853 |
| TOMM40 | O96008 | 0.151902137 | 16.40672496 | 7 m$H_2O$ vs 3 mm $H_2O$ | 0.203345291 | 0.744109691 |
| BGN | P21810 | -0.109414166 | 17.72622742 | 7 m$H_2O$ vs 3 mm $H_2O$ | 0.212157087 | 0.755535458 |
| RPL32 | P62910 | -0.053075986 | 20.21761555 | 7 m$H_2O$ vs 3 mm $H_2O$ | 0.214413254 | 0.755535458 |
| RPL4 | P36578 | -0.047046989 | 20.17098489 | 7 m$H_2O$ vs 3 mm $H_2O$ | 0.21630159 | 0.755535458 |
| ACLY | P53396 | -0.060644243 | 20.50835424 | 7 m$H_2O$ vs 3 mm $H_2O$ | 0.217166766 | 0.755535458 |
| FGB | P02675 | 0.077053222 | 26.64713393 | 7 m$H_2O$ vs 3 mm $H_2O$ | 0.217812024 | 0.755535458 |
| RPL27A | P46776 | -0.32847819 | 18.54518335 | 7 m$H_2O$ vs 3 mm $H_2O$ | 0.22774543 | 0.781847713 |
| CCT5 | P48643 | 0.064851925 | 18.65229805 | 7 m$H_2O$ vs 3 mm $H_2O$ | 0.232415864 | 0.786605841 |
| PKM | P14618 | -0.064937108 | 21.11943098 | 7 m$H_2O$ vs 3 mm $H_2O$ | 0.233855791 | 0.786605841 |
| MYO1C | O00159 | 0.065312551 | 19.20310494 | 7 m$H_2O$ vs 3 mm $H_2O$ | 0.238015951 | 0.792593116 |
| HTRA1 | Q92743 | -0.113888954 | 19.88510114 | 7 m$H_2O$ vs 3 mm $H_2O$ | 0.242712949 | 0.800231802 |
| RPL11 | P62913 | -0.070405561 | 20.19794449 | 7 m$H_2O$ vs 3 mm $H_2O$ | 0.259903205 | 0.804945319 |
| DLAT | P10515 | 0.106164014 | 17.13024169 | 7 m$H_2O$ vs 3 mm $H_2O$ | 0.269590056 | 0.804945319 |
| COL6A2 | P12110 | -0.060528228 | 23.03754647 | 7 m$H_2O$ vs 3 mm $H_2O$ | 0.270951338 | 0.804945319 |
| RPL31 | P62899 | -0.05867508 | 20.04161035 | 7 m$H_2O$ vs 3 mm $H_2O$ | 0.273781601 | 0.804945319 |
| FBN1 | P35555 | -0.071076398 | 21.77901541 | 7 m$H_2O$ vs 3 mm $H_2O$ | 0.275010295 | 0.804945319 |
| COL3A1 | P02461 | -0.081673893 | 18.87600336 | 7 m$H_2O$ vs 3 mm $H_2O$ | 0.275103903 | 0.804945319 |
| SRPX | P78539 | -0.065110636 | 19.32751409 | 7 m$H_2O$ vs 3 mm $H_2O$ | 0.277642346 | 0.804945319 |
| GLS | O94925 | 0.085449274 | 15.35759825 | 7 m$H_2O$ vs 3 mm $H_2O$ | 0.277974022 | 0.804945319 |
| EIF4G1 | Q04637 | 0.056450341 | 17.58873533 | 7 m$H_2O$ vs 3 mm $H_2O$ | 0.278706149 | 0.804945319 |

| Gene | Protein ID | logFC | AveExpr | comparison | pvalue | fdr |
|---|---|---|---|---|---|---|
| PSMD12 | O00232 | 0.080180678 | 16.03269934 | 7 m$H_2O$ vs 3 mm $H_2O$ | 0.278924618 | 0.804945319 |
| CLU | P10909 | -0.07243156 | 20.64891502 | 7 m$H_2O$ vs 3 mm $H_2O$ | 0.280368103 | 0.804945319 |
| MYO1B | O43795 | -0.192193149 | 15.60251924 | 7 m$H_2O$ vs 3 mm $H_2O$ | 0.283667698 | 0.804945319 |
| FN1 | P02751 | -0.073732206 | 25.93912475 | 7 m$H_2O$ vs 3 mm $H_2O$ | 0.283715094 | 0.804945319 |
| FGG | P02679 | 0.075630713 | 25.08264222 | 7 m$H_2O$ vs 3 mm $H_2O$ | 0.284337199 | 0.804945319 |
| RPS19 | P39019 | 0.085861616 | 16.54266348 | 7 m$H_2O$ vs 3 mm $H_2O$ | 0.289285916 | 0.804945319 |
| RPS15A | P62244 | -0.054992271 | 16.81721761 | 7 m$H_2O$ vs 3 mm $H_2O$ | 0.290027431 | 0.804945319 |
| CFL1 | P23528 | 0.115006937 | 18.71089364 | 7 m$H_2O$ vs 3 mm $H_2O$ | 0.291810436 | 0.804945319 |
| HNRNPA2B1 | P22626 | 0.087750091 | 18.53624742 | 7 m$H_2O$ vs 3 mm $H_2O$ | 0.299248726 | 0.804945319 |
| P4HB | P07237 | 0.062501487 | 19.25928557 | 7 m$H_2O$ vs 3 mm $H_2O$ | 0.300679916 | 0.804945319 |
| FBN2 | P35556 | -0.111219278 | 19.4779791 | 7 m$H_2O$ vs 3 mm $H_2O$ | 0.303296251 | 0.804945319 |
| ACTN4 | O43707 | -0.052296734 | 20.91981811 | 7 m$H_2O$ vs 3 mm $H_2O$ | 0.312185301 | 0.804945319 |
| RPL23A | P62750 | -0.104821425 | 20.24011738 | 7 m$H_2O$ vs 3 mm $H_2O$ | 0.312797806 | 0.804945319 |
| TMED10 | P49755 | 0.108326644 | 16.93071406 | 7 m$H_2O$ vs 3 mm $H_2O$ | 0.313345197 | 0.804945319 |
| PSMD2 | Q13200 | 0.05798114 | 17.68793105 | 7 m$H_2O$ vs 3 mm $H_2O$ | 0.3136219 | 0.804945319 |
| RAB10 | P61026 | 0.073169408 | 15.89189572 | 7 m$H_2O$ vs 3 mm $H_2O$ | 0.31874791 | 0.804945319 |
| PTGES3 | Q15185 | -0.074426423 | 20.03155888 | 7 m$H_2O$ vs 3 mm $H_2O$ | 0.321604787 | 0.804945319 |
| CAVIN1 | Q6NZI2 | 0.058599947 | 19.48843453 | 7 m$H_2O$ vs 3 mm $H_2O$ | 0.326133352 | 0.804945319 |
| RAB7A | P51149 | 0.057157916 | 17.51936984 | 7 m$H_2O$ vs 3 mm $H_2O$ | 0.328411692 | 0.804945319 |
| NDUFA2 | O43678 | 0.052818109 | 17.93233932 | 7 m$H_2O$ vs 3 mm $H_2O$ | 0.329590912 | 0.804945319 |
| EEF2 | P13639 | -0.040997183 | 21.48552521 | 7 m$H_2O$ vs 3 mm $H_2O$ | 0.330477669 | 0.804945319 |
| RPL6 | Q02878 | 0.042348433 | 21.67881898 | 7 m$H_2O$ vs 3 mm $H_2O$ | 0.331653744 | 0.804945319 |
| C5 | P01031 | 0.067944101 | 18.82768108 | 7 m$H_2O$ vs 3 mm $H_2O$ | 0.333012116 | 0.804945319 |
| RAP1A | P62834 | -0.046502054 | 20.39007357 | 7 m$H_2O$ vs 3 mm $H_2O$ | 0.333772688 | 0.804945319 |
| ANXA1 | P04083 | 0.070039774 | 19.10152145 | 7 m$H_2O$ vs 3 mm $H_2O$ | 0.335314434 | 0.804945319 |
| RPS5 | P46782 | -0.079386197 | 21.16789776 | 7 m$H_2O$ vs 3 mm $H_2O$ | 0.336014111 | 0.804945319 |
| HM13 | Q8TCT9 | 0.082570191 | 17.14646182 | 7 m$H_2O$ vs 3 mm $H_2O$ | 0.342108771 | 0.804945319 |
| KRT18 | P05783 | -0.098852752 | 21.504805 | 7 m$H_2O$ vs 3 mm $H_2O$ | 0.342877953 | 0.804945319 |
| FLNA | P21333 | -0.039727558 | 22.91262799 | 7 m$H_2O$ vs 3 mm $H_2O$ | 0.344340585 | 0.804945319 |
| CALR | P27797 | 0.097179852 | 18.69992242 | 7 m$H_2O$ vs 3 mm $H_2O$ | 0.345672753 | 0.804945319 |
| RPLP2 | P05387 | 0.067171571 | 18.74895696 | 7 m$H_2O$ vs 3 mm $H_2O$ | 0.347187159 | 0.804945319 |
| NDUFB10 | O96000 | -0.068242082 | 17.39845643 | 7 m$H_2O$ vs 3 mm $H_2O$ | 0.349279067 | 0.804945319 |
| PFKL | P17858 | -0.079547924 | 19.53869474 | 7 m$H_2O$ vs 3 mm $H_2O$ | 0.351842419 | 0.804945319 |
| ARF4 | P18085 | 0.065012822 | 18.52619066 | 7 m$H_2O$ vs 3 mm $H_2O$ | 0.353898165 | 0.804945319 |
| MTCH2 | Q9Y6C9 | -0.063000503 | 19.00153265 | 7 m$H_2O$ vs 3 mm $H_2O$ | 0.354030203 | 0.804945319 |
| RPS18 | P62269 | -0.044035644 | 21.59125597 | 7 m$H_2O$ vs 3 mm $H_2O$ | 0.356025237 | 0.804945319 |
| ARPC1B | O15143 | -0.056379199 | 19.59422932 | 7 m$H_2O$ vs 3 mm $H_2O$ | 0.356055949 | 0.804945319 |
| C3 | P01024 | 0.073951153 | 18.48488775 | 7 m$H_2O$ vs 3 mm $H_2O$ | 0.357753475 | 0.804945319 |
| AARS1 | P49588 | 0.066959871 | 16.12354083 | 7 m$H_2O$ vs 3 mm $H_2O$ | 0.364182088 | 0.806071793 |
| NID2 | Q14112 | -0.059790565 | 20.71553975 | 7 m$H_2O$ vs 3 mm $H_2O$ | 0.364421434 | 0.806071793 |
| LETM1 | O95202 | 0.070511578 | 17.41326173 | 7 m$H_2O$ vs 3 mm $H_2O$ | 0.365516038 | 0.806071793 |

| Gene | Protein ID | logFC | AveExpr | comparison | pvalue | fdr |
|---|---|---|---|---|---|---|
| HNRNPK | P61978 | -0.069561532 | 19.95715687 | 7 m$H_2O$ vs 3 mm $H_2O$ | 0.373806329 | 0.80805965 |
| SAMM50 | Q9Y512 | 0.068018273 | 14.15564128 | 7 m$H_2O$ vs 3 mm $H_2O$ | 0.375169412 | 0.80805965 |
| ETF1 | P62495 | 0.075378384 | 16.60031114 | 7 m$H_2O$ vs 3 mm $H_2O$ | 0.375259063 | 0.80805965 |
| COL6A1 | P12109 | -0.05804019 | 22.09386598 | 7 m$H_2O$ vs 3 mm $H_2O$ | 0.376123861 | 0.80805965 |
| LAMA4 | Q16363 | 0.038322917 | 21.38083312 | 7 m$H_2O$ vs 3 mm $H_2O$ | 0.398389052 | 0.850407399 |
| RPL24 | P83731 | 0.095307734 | 15.41834801 | 7 m$H_2O$ vs 3 mm $H_2O$ | 0.412573536 | 0.872424361 |
| ACTN1 | P12814 | -0.034705284 | 21.41145854 | 7 m$H_2O$ vs 3 mm $H_2O$ | 0.414996203 | 0.872424361 |
| CAPZB | P47756 | 0.033312947 | 18.76183816 | 7 m$H_2O$ vs 3 mm $H_2O$ | 0.419365528 | 0.872424361 |
| LRRC59 | Q96AG4 | -0.052442533 | 17.79277083 | 7 m$H_2O$ vs 3 mm $H_2O$ | 0.420907647 | 0.872424361 |
| MYH10 | P35580 | 0.056294567 | 23.55126614 | 7 m$H_2O$ vs 3 mm $H_2O$ | 0.422789569 | 0.872424361 |
| EHD2 | Q9NZN4 | 0.043451278 | 17.32935895 | 7 m$H_2O$ vs 3 mm $H_2O$ | 0.426075668 | 0.872424361 |
| LGALS1 | P09382 | -0.048067614 | 19.93191209 | 7 m$H_2O$ vs 3 mm $H_2O$ | 0.427042555 | 0.872424361 |
| COL6A3 | P12111 | -0.042479054 | 25.25110137 | 7 m$H_2O$ vs 3 mm $H_2O$ | 0.437774522 | 0.888895829 |
| HSPG2 | P98160 | -0.039018482 | 24.74423208 | 7 m$H_2O$ vs 3 mm $H_2O$ | 0.447993913 | 0.890528863 |
| RPS4X | P62701 | -0.030016813 | 20.11765589 | 7 m$H_2O$ vs 3 mm $H_2O$ | 0.448267894 | 0.890528863 |
| RPS16 | P62249 | 0.045651041 | 21.02085815 | 7 m$H_2O$ vs 3 mm $H_2O$ | 0.450412758 | 0.890528863 |
| TAGLN | Q01995 | -0.058624219 | 18.81489542 | 7 m$H_2O$ vs 3 mm $H_2O$ | 0.452705625 | 0.890528863 |
| PRKDC | P78527 | 0.031033025 | 19.38685821 | 7 m$H_2O$ vs 3 mm $H_2O$ | 0.454841168 | 0.890528863 |
| VCL | P18206 | 0.08556574 | 16.93515267 | 7 m$H_2O$ vs 3 mm $H_2O$ | 0.458580368 | 0.890528863 |
| TPM3 | P06753 | 0.081721106 | 19.40677325 | 7 m$H_2O$ vs 3 mm $H_2O$ | 0.459553111 | 0.890528863 |
| COL12A1 | Q99715 | 0.045509372 | 19.23200628 | 7 m$H_2O$ vs 3 mm $H_2O$ | 0.459972866 | 0.890528863 |
| COL5A1 | P20908 | -0.045582927 | 19.65823476 | 7 m$H_2O$ vs 3 mm $H_2O$ | 0.463060487 | 0.891319669 |
| MFGE8 | Q08431 | -0.044057653 | 17.96391857 | 7 m$H_2O$ vs 3 mm $H_2O$ | 0.465734602 | 0.891319669 |
| ATL3 | Q6DD88 | 0.031925243 | 19.24931374 | 7 m$H_2O$ vs 3 mm $H_2O$ | 0.468579066 | 0.891639023 |
| SSBP1 | Q04837 | 0.036946617 | 18.41703518 | 7 m$H_2O$ vs 3 mm $H_2O$ | 0.478202371 | 0.898590587 |
| RPL19 | P84098 | -0.046680205 | 19.74579175 | 7 m$H_2O$ vs 3 mm $H_2O$ | 0.479320099 | 0.898590587 |
| MMRN1 | Q13201 | -0.049714584 | 21.09043115 | 7 m$H_2O$ vs 3 mm $H_2O$ | 0.486332559 | 0.898590587 |
| FLOT1 | O75955 | 0.044624954 | 18.47467429 | 7 m$H_2O$ vs 3 mm $H_2O$ | 0.487055666 | 0.898590587 |
| WDR1 | O75083 | -0.048458238 | 18.94923163 | 7 m$H_2O$ vs 3 mm $H_2O$ | 0.489724386 | 0.898590587 |
| RPL28 | P46779 | -0.038680034 | 21.24799089 | 7 m$H_2O$ vs 3 mm $H_2O$ | 0.489909524 | 0.898590587 |
| CCT8 | P50990 | 0.040521066 | 19.07398562 | 7 m$H_2O$ vs 3 mm $H_2O$ | 0.49183309 | 0.898590587 |
| RPL10A | P62906 | -0.03164489 | 19.96141059 | 7 m$H_2O$ vs 3 mm $H_2O$ | 0.499606756 | 0.898590587 |
| CS | O75390 | 0.034123514 | 18.86813435 | 7 m$H_2O$ vs 3 mm $H_2O$ | 0.500314481 | 0.898590587 |
| SERPINE1 | P05121 | 0.042335946 | 20.70463064 | 7 m$H_2O$ vs 3 mm $H_2O$ | 0.501163787 | 0.898590587 |
| CCT3 | P49368 | -0.035339994 | 20.32312181 | 7 m$H_2O$ vs 3 mm $H_2O$ | 0.501915463 | 0.898590587 |
| RPL18 | Q07020 | -0.034561328 | 21.23440702 | 7 m$H_2O$ vs 3 mm $H_2O$ | 0.509170332 | 0.904078261 |
| VIM | P08670 | -0.039440034 | 23.56319756 | 7 m$H_2O$ vs 3 mm $H_2O$ | 0.51041055 | 0.904078261 |
| VDAC2 | P45880 | -0.028386172 | 21.04702323 | 7 m$H_2O$ vs 3 mm $H_2O$ | 0.516786187 | 0.909570646 |
| RPS3A | P61247 | -0.025919942 | 21.14483502 | 7 m$H_2O$ vs 3 mm $H_2O$ | 0.518974242 | 0.909570646 |
| HSP90AB1 | P08238 | -0.03754769 | 20.68956491 | 7 m$H_2O$ vs 3 mm $H_2O$ | 0.527759062 | 0.910419105 |
| IKBIP | Q70UQ0 | -0.035790533 | 16.34942842 | 7 m$H_2O$ vs 3 mm $H_2O$ | 0.531289483 | 0.910419105 |

| Gene | Protein ID | logFC | AveExpr | comparison | pvalue | fdr |
|---|---|---|---|---|---|---|
| EIF3E | P60228 | 0.031001531 | 17.00277122 | 7 m$H_2O$ vs 3 mm $H_2O$ | 0.532907759 | 0.910419105 |
| MACROH2A1 | O75367 | 0.055760211 | 18.22980609 | 7 m$H_2O$ vs 3 mm $H_2O$ | 0.536450178 | 0.910419105 |
| MVP | Q14764 | -0.028496225 | 22.37678295 | 7 m$H_2O$ vs 3 mm $H_2O$ | 0.537592723 | 0.910419105 |
| DNAJB4 | Q9UDY4 | 0.033471591 | 17.32499825 | 7 m$H_2O$ vs 3 mm $H_2O$ | 0.541711881 | 0.910419105 |
| LARS1 | Q9P2J5 | 0.045608778 | 16.82750163 | 7 m$H_2O$ vs 3 mm $H_2O$ | 0.547560536 | 0.910419105 |
| RPL13 | P26373 | -0.035792545 | 19.8746121 | 7 m$H_2O$ vs 3 mm $H_2O$ | 0.548355849 | 0.910419105 |
| NDUFA13 | Q9P0J0 | 0.044148008 | 17.52794809 | 7 m$H_2O$ vs 3 mm $H_2O$ | 0.549461212 | 0.910419105 |
| VWF | P04275 | -0.054332596 | 22.88791985 | 7 m$H_2O$ vs 3 mm $H_2O$ | 0.55258734 | 0.910419105 |
| ATP6V0D1 | P61421 | 0.043690328 | 17.34164973 | 7 m$H_2O$ vs 3 mm $H_2O$ | 0.55561842 | 0.910419105 |
| RPS9 | P46781 | 0.024883891 | 21.30621062 | 7 m$H_2O$ vs 3 mm $H_2O$ | 0.55671632 | 0.910419105 |
| VDAC1 | P21796 | 0.045829154 | 20.43149002 | 7 m$H_2O$ vs 3 mm $H_2O$ | 0.563919327 | 0.910419105 |
| RPS8 | P62241 | 0.035198323 | 20.45506954 | 7 m$H_2O$ vs 3 mm $H_2O$ | 0.57052259 | 0.910419105 |
| PLOD2 | O00469 | -0.025102299 | 18.32332173 | 7 m$H_2O$ vs 3 mm $H_2O$ | 0.571736086 | 0.910419105 |
| EEF1D | P29692 | -0.021165886 | 17.18896539 | 7 m$H_2O$ vs 3 mm $H_2O$ | 0.572539283 | 0.910419105 |
| DDOST | P39656 | 0.040090854 | 18.12022848 | 7 m$H_2O$ vs 3 mm $H_2O$ | 0.575113496 | 0.910419105 |
| LAMB1 | P07942 | 0.030688909 | 21.8352099 | 7 m$H_2O$ vs 3 mm $H_2O$ | 0.583502227 | 0.910419105 |
| CAV1 | Q03135 | -0.031543815 | 21.4061969 | 7 m$H_2O$ vs 3 mm $H_2O$ | 0.584938475 | 0.910419105 |
| CCT4 | P50991 | 0.021961456 | 18.88952591 | 7 m$H_2O$ vs 3 mm $H_2O$ | 0.585662938 | 0.910419105 |
| DHCR7 | Q9UBM7 | 0.033705952 | 16.32955993 | 7 m$H_2O$ vs 3 mm $H_2O$ | 0.58898571 | 0.910419105 |
| RPL7 | P18124 | -0.020861694 | 21.07465763 | 7 m$H_2O$ vs 3 mm $H_2O$ | 0.58960002 | 0.910419105 |
| G6PD | P11413 | 0.043211609 | 19.4860753 | 7 m$H_2O$ vs 3 mm $H_2O$ | 0.590735428 | 0.910419105 |
| NDUFS2 | O75306 | 0.034998653 | 15.97505329 | 7 m$H_2O$ vs 3 mm $H_2O$ | 0.593272323 | 0.910419105 |
| ITIH2 | P19823 | -0.085174157 | 21.56628132 | 7 m$H_2O$ vs 3 mm $H_2O$ | 0.593352708 | 0.910419105 |
| COL18A1 | P39060 | 0.030918355 | 21.76400023 | 7 m$H_2O$ vs 3 mm $H_2O$ | 0.598667998 | 0.910419105 |
| TINAGL1 | Q9GZM7 | -0.037151788 | 19.31518041 | 7 m$H_2O$ vs 3 mm $H_2O$ | 0.599895883 | 0.910419105 |
| ARPC2 | O15144 | -0.030474701 | 18.22115014 | 7 m$H_2O$ vs 3 mm $H_2O$ | 0.600479591 | 0.910419105 |
| MACF1 | Q9UPN3 | -0.029060792 | 19.0594487 | 7 m$H_2O$ vs 3 mm $H_2O$ | 0.60484067 | 0.910419105 |
| CANX | P27824 | -0.046809711 | 20.42117326 | 7 m$H_2O$ vs 3 mm $H_2O$ | 0.60594025 | 0.910419105 |
| TMEM33 | P57088 | 0.073113469 | 15.21113383 | 7 m$H_2O$ vs 3 mm $H_2O$ | 0.606111786 | 0.910419105 |
| DARS1 | P14868 | 0.023382548 | 18.31722998 | 7 m$H_2O$ vs 3 mm $H_2O$ | 0.606971679 | 0.910419105 |
| BAG2 | O95816 | -0.029748889 | 16.49986499 | 7 m$H_2O$ vs 3 mm $H_2O$ | 0.609680061 | 0.910419105 |
| TUBB | P07437 | 0.026755463 | 22.27281811 | 7 m$H_2O$ vs 3 mm $H_2O$ | 0.614544393 | 0.913586084 |
| DYNC1H1 | Q14204 | 0.019598478 | 21.33031749 | 7 m$H_2O$ vs 3 mm $H_2O$ | 0.622940511 | 0.916375877 |
| EMILIN1 | Q9Y6C2 | -0.030369385 | 18.74132285 | 7 m$H_2O$ vs 3 mm $H_2O$ | 0.626592646 | 0.916375877 |
| MYADM | Q96S97 | -0.030936964 | 17.86105498 | 7 m$H_2O$ vs 3 mm $H_2O$ | 0.629012872 | 0.916375877 |
| H1-0 | P07305 | 0.034364065 | 18.2800493 | 7 m$H_2O$ vs 3 mm $H_2O$ | 0.629691725 | 0.916375877 |
| PRDX6 | P30041 | -0.037224096 | 16.40196639 | 7 m$H_2O$ vs 3 mm $H_2O$ | 0.63842436 | 0.916375877 |
| ANXA2 | P07355 | -0.042460479 | 21.4349227 | 7 m$H_2O$ vs 3 mm $H_2O$ | 0.639026481 | 0.916375877 |
| DSTN | P60981 | -0.025706327 | 18.25003562 | 7 m$H_2O$ vs 3 mm $H_2O$ | 0.644420391 | 0.916375877 |
| COL1A2 | P08123 | 0.057209266 | 16.18730792 | 7 m$H_2O$ vs 3 mm $H_2O$ | 0.645682638 | 0.916375877 |
| FLNC | Q14315 | -0.021620417 | 20.94363974 | 7 m$H_2O$ vs 3 mm $H_2O$ | 0.650231046 | 0.916375877 |

| Gene | Protein ID | logFC | AveExpr | comparison | pvalue | fdr |
|---|---|---|---|---|---|---|
| COX4I1 | P13073 | -0.03926026 | 18.4693388 | 7 $mH_2O$ vs 3 mm $H_2O$ | 0.652397211 | 0.916375877 |
| RPL35A | P18077 | 0.020704818 | 19.59741322 | 7 $mH_2O$ vs 3 mm $H_2O$ | 0.654707004 | 0.916375877 |
| CCT6A | P40227 | -0.027463889 | 18.08843594 | 7 $mH_2O$ vs 3 mm $H_2O$ | 0.656613642 | 0.916375877 |
| RPS3 | P23396 | -0.017489715 | 21.84558542 | 7 $mH_2O$ vs 3 mm $H_2O$ | 0.658091156 | 0.916375877 |
| ECI2 | O75521 | 0.025526462 | 16.270599 | 7 $mH_2O$ vs 3 mm $H_2O$ | 0.663801592 | 0.916375877 |
| RPL22 | P35268 | -0.031323862 | 19.68733867 | 7 $mH_2O$ vs 3 mm $H_2O$ | 0.672522222 | 0.916375877 |
| ACTR3 | P61158 | -0.030761577 | 18.92023602 | 7 $mH_2O$ vs 3 mm $H_2O$ | 0.673511533 | 0.916375877 |
| RPS2 | P15880 | -0.021911391 | 20.34911658 | 7 $mH_2O$ vs 3 mm $H_2O$ | 0.67768217 | 0.916375877 |
| PLEC | Q15149 | 0.017911357 | 23.08702195 | 7 $mH_2O$ vs 3 mm $H_2O$ | 0.678676777 | 0.916375877 |
| ATP5PD | O75947 | -0.025626637 | 17.79180696 | 7 $mH_2O$ vs 3 mm $H_2O$ | 0.680309725 | 0.916375877 |
| CKAP4 | Q07065 | 0.021268257 | 20.5524488 | 7 $mH_2O$ vs 3 mm $H_2O$ | 0.681160485 | 0.916375877 |
| ALDH18A1 | P54886 | -0.032561504 | 15.2913844 | 7 $mH_2O$ vs 3 mm $H_2O$ | 0.685014645 | 0.916375877 |
| COL1A1 | P02452 | 0.01821884 | 19.50614173 | 7 $mH_2O$ vs 3 mm $H_2O$ | 0.685031348 | 0.916375877 |
| NT5E | P21589 | 0.023836129 | 18.06619473 | 7 $mH_2O$ vs 3 mm $H_2O$ | 0.688403825 | 0.916375877 |
| RPL7A | P62424 | -0.030872278 | 20.68475958 | 7 $mH_2O$ vs 3 mm $H_2O$ | 0.695907754 | 0.916375877 |
| CHCHD3 | Q9NX63 | 0.020115533 | 20.25383758 | 7 $mH_2O$ vs 3 mm $H_2O$ | 0.696822936 | 0.916375877 |
| RPS25 | P62851 | 0.027429261 | 18.6137467 | 7 $mH_2O$ vs 3 mm $H_2O$ | 0.699673527 | 0.916375877 |
| SPTAN1 | Q13813 | 0.039996114 | 18.24571041 | 7 $mH_2O$ vs 3 mm $H_2O$ | 0.699914645 | 0.916375877 |
| PLOD1 | Q02809 | -0.0263696 | 18.19084371 | 7 $mH_2O$ vs 3 mm $H_2O$ | 0.702497346 | 0.916375877 |
| TLN1 | Q9Y490 | 0.017705694 | 19.12623259 | 7 $mH_2O$ vs 3 mm $H_2O$ | 0.708412405 | 0.916375877 |
| ACOT7 | O00154 | 0.017232945 | 15.53290167 | 7 $mH_2O$ vs 3 mm $H_2O$ | 0.709647017 | 0.916375877 |
| ATP5F1D | P30049 | -0.027565271 | 19.44075806 | 7 $mH_2O$ vs 3 mm $H_2O$ | 0.710451405 | 0.916375877 |
| PRXL2A | Q9BRX8 | 0.041724302 | 18.71078175 | 7 $mH_2O$ vs 3 mm $H_2O$ | 0.711813682 | 0.916375877 |
| HSPD1 | P10809 | 0.018760127 | 21.18631581 | 7 $mH_2O$ vs 3 mm $H_2O$ | 0.714333057 | 0.916375877 |
| EIF4A1 | P60842 | -0.016902041 | 21.20872477 | 7 $mH_2O$ vs 3 mm $H_2O$ | 0.715222349 | 0.916375877 |
| LRPPRC | P42704 | -0.022673626 | 19.08842444 | 7 $mH_2O$ vs 3 mm $H_2O$ | 0.721902061 | 0.916375877 |
| VARS1 | P26640 | -0.024960586 | 17.26292586 | 7 $mH_2O$ vs 3 mm $H_2O$ | 0.723254279 | 0.916375877 |
| KRT8 | P05787 | -0.079383091 | 22.31188608 | 7 $mH_2O$ vs 3 mm $H_2O$ | 0.728058852 | 0.916375877 |
| XRCC6 | P12956 | -0.026546917 | 17.85962569 | 7 $mH_2O$ vs 3 mm $H_2O$ | 0.728157613 | 0.916375877 |
| GSN | P06396 | -0.061138783 | 18.09615138 | 7 $mH_2O$ vs 3 mm $H_2O$ | 0.728483118 | 0.916375877 |
| EIF2AK2 | P19525 | 0.018353926 | 15.71603506 | 7 $mH_2O$ vs 3 mm $H_2O$ | 0.729019896 | 0.916375877 |
| RPS11 | P62280 | -0.021077552 | 20.10277157 | 7 $mH_2O$ vs 3 mm $H_2O$ | 0.73132517 | 0.916375877 |
| MARCKS | P29966 | -0.033056396 | 16.38719431 | 7 $mH_2O$ vs 3 mm $H_2O$ | 0.73199995 | 0.916375877 |
| CLTC | Q00610 | -0.029924809 | 20.30142585 | 7 $mH_2O$ vs 3 mm $H_2O$ | 0.741499258 | 0.920186362 |
| PFKP | Q01813 | -0.015416034 | 20.72340758 | 7 $mH_2O$ vs 3 mm $H_2O$ | 0.742705613 | 0.920186362 |
| RPS12 | P25398 | -0.028889744 | 16.01206618 | 7 $mH_2O$ vs 3 mm $H_2O$ | 0.743333728 | 0.920186362 |
| RPS13 | P62277 | -0.015694424 | 18.74926133 | 7 $mH_2O$ vs 3 mm $H_2O$ | 0.75055694 | 0.92234656 |
| RPL34 | P49207 | 0.01821332 | 19.23488626 | 7 $mH_2O$ vs 3 mm $H_2O$ | 0.750618372 | 0.92234656 |
| PECAM1 | P16284 | -0.058902312 | 15.97467544 | 7 $mH_2O$ vs 3 mm $H_2O$ | 0.75819322 | 0.923590453 |
| TMEM43 | Q9BTV4 | 0.026323032 | 18.65996781 | 7 $mH_2O$ vs 3 mm $H_2O$ | 0.76074934 | 0.923590453 |
| IQGAP1 | P46940 | -0.019542688 | 18.57483499 | 7 $mH_2O$ vs 3 mm $H_2O$ | 0.7629661 | 0.923590453 |

| Gene | Protein ID | logFC | AveExpr | comparison | pvalue | fdr |
|---|---|---|---|---|---|---|
| EIF4G2 | P78344 | 0.015781444 | 18.67469545 | 7 m$H_2O$ vs 3 mm $H_2O$ | 0.764275494 | 0.923590453 |
| VCAN | P13611 | -0.013966745 | 19.33335978 | 7 m$H_2O$ vs 3 mm $H_2O$ | 0.765498393 | 0.923590453 |
| PHB2 | Q99623 | -0.026766257 | 20.07531942 | 7 m$H_2O$ vs 3 mm $H_2O$ | 0.770913842 | 0.926168458 |
| ATP5F1A | P25705 | -0.0157546 | 20.00502696 | 7 m$H_2O$ vs 3 mm $H_2O$ | 0.775109772 | 0.926168458 |
| HSD17B10 | Q99714 | 0.01408564 | 18.62299534 | 7 m$H_2O$ vs 3 mm $H_2O$ | 0.775978978 | 0.926168458 |
| RPL3 | P39023 | -0.010988877 | 21.79984935 | 7 m$H_2O$ vs 3 mm $H_2O$ | 0.792414926 | 0.942407751 |
| RUVBL2 | Q9Y230 | -0.01373392 | 18.15268106 | 7 m$H_2O$ vs 3 mm $H_2O$ | 0.800920972 | 0.947057741 |
| GARS1 | P41250 | 0.012220871 | 17.42784741 | 7 m$H_2O$ vs 3 mm $H_2O$ | 0.802012861 | 0.947057741 |
| MYL6 | P60660 | -0.022074632 | 21.82604227 | 7 m$H_2O$ vs 3 mm $H_2O$ | 0.805058471 | 0.947081226 |
| RPL27 | P61353 | -0.015958561 | 19.40289322 | 7 m$H_2O$ vs 3 mm $H_2O$ | 0.811310322 | 0.947081226 |
| POSTN | Q15063 | -0.034577547 | 16.30898796 | 7 m$H_2O$ vs 3 mm $H_2O$ | 0.815658936 | 0.947081226 |
| RAN | P62826 | 0.01302491 | 20.33795383 | 7 m$H_2O$ vs 3 mm $H_2O$ | 0.815852918 | 0.947081226 |
| FASN | P49327 | -0.010786669 | 19.88841937 | 7 m$H_2O$ vs 3 mm $H_2O$ | 0.816253189 | 0.947081226 |
| PSMC2 | P35998 | 0.010166139 | 17.63702096 | 7 m$H_2O$ vs 3 mm $H_2O$ | 0.820848646 | 0.949106247 |
| RPL30 | P62888 | -0.016915664 | 16.60151038 | 7 m$H_2O$ vs 3 mm $H_2O$ | 0.82982558 | 0.949996708 |
| MSN | P26038 | -0.012003576 | 18.77231066 | 7 m$H_2O$ vs 3 mm $H_2O$ | 0.829840968 | 0.949996708 |
| EEF1G | P26641 | -0.008314968 | 19.6645041 | 7 m$H_2O$ vs 3 mm $H_2O$ | 0.830630523 | 0.949996708 |
| RPS17 | P08708 | 0.013198748 | 19.20866766 | 7 m$H_2O$ vs 3 mm $H_2O$ | 0.833413627 | 0.949996708 |
| MYH9 | P35579 | -0.01466176 | 25.82383901 | 7 m$H_2O$ vs 3 mm $H_2O$ | 0.840450899 | 0.949996708 |
| MDH2 | P40926 | -0.012849753 | 18.21421021 | 7 m$H_2O$ vs 3 mm $H_2O$ | 0.841392588 | 0.949996708 |
| EIF4A3 | P38919 | 0.022692021 | 17.58417509 | 7 m$H_2O$ vs 3 mm $H_2O$ | 0.844882185 | 0.949996708 |
| COPA | P53621 | -0.010524652 | 18.80077329 | 7 m$H_2O$ vs 3 mm $H_2O$ | 0.845014948 | 0.949996708 |
| RPS23 | P62266 | -0.020756622 | 19.05334112 | 7 m$H_2O$ vs 3 mm $H_2O$ | 0.847725825 | 0.949996708 |
| RRBP1 | Q9P2E9 | 0.019144982 | 16.81832112 | 7 m$H_2O$ vs 3 mm $H_2O$ | 0.850526446 | 0.949996708 |
| SERPINH1 | P50454 | 0.015693307 | 20.57627706 | 7 m$H_2O$ vs 3 mm $H_2O$ | 0.853000047 | 0.949996708 |
| CCN1 | O00622 | -0.011129617 | 20.29624494 | 7 m$H_2O$ vs 3 mm $H_2O$ | 0.862400109 | 0.956355707 |
| TGM2 | P21980 | -0.00838671 | 22.41037841 | 7 m$H_2O$ vs 3 mm $H_2O$ | 0.867403633 | 0.956355707 |
| PSMC5 | P62195 | -0.00844414 | 16.40322065 | 7 m$H_2O$ vs 3 mm $H_2O$ | 0.869527214 | 0.956355707 |
| RPN1 | P04843 | -0.007343555 | 20.58979474 | 7 m$H_2O$ vs 3 mm $H_2O$ | 0.870197535 | 0.956355707 |
| PCBP1 | Q15365 | 0.006256483 | 18.89923787 | 7 m$H_2O$ vs 3 mm $H_2O$ | 0.878410068 | 0.959209459 |
| HSPA8 | P11142 | -0.00774493 | 22.30402347 | 7 m$H_2O$ vs 3 mm $H_2O$ | 0.880675148 | 0.959209459 |
| RUVBL1 | Q9Y265 | -0.022388373 | 14.1186164 | 7 m$H_2O$ vs 3 mm $H_2O$ | 0.88143572 | 0.959209459 |
| RACK1 | P63244 | -0.006160038 | 20.05788219 | 7 m$H_2O$ vs 3 mm $H_2O$ | 0.888555374 | 0.963807621 |
| RPS6 | P62753 | 0.011394227 | 21.27019737 | 7 m$H_2O$ vs 3 mm $H_2O$ | 0.8954292 | 0.968110142 |
| PHB1 | P35232 | -0.01157469 | 19.87316936 | 7 m$H_2O$ vs 3 mm $H_2O$ | 0.902062088 | 0.970484926 |
| AHNAK | Q09666 | -0.010369186 | 20.16252586 | 7 m$H_2O$ vs 3 mm $H_2O$ | 0.903454436 | 0.970484926 |
| NONO | Q15233 | -0.008761192 | 19.26168669 | 7 m$H_2O$ vs 3 mm $H_2O$ | 0.908988271 | 0.973289692 |
| HSPA5 | P11021 | -0.006433007 | 21.89509302 | 7 m$H_2O$ vs 3 mm $H_2O$ | 0.921834013 | 0.981239451 |
| PTX3 | P26022 | 0.012024985 | 20.76887638 | 7 m$H_2O$ vs 3 mm $H_2O$ | 0.923664838 | 0.981239451 |
| PRDX5 | P30044 | 0.006377706 | 16.63267706 | 7 m$H_2O$ vs 3 mm $H_2O$ | 0.925252816 | 0.981239451 |
| HSPB1 | P04792 | -0.004190541 | 20.58486216 | 7 m$H_2O$ vs 3 mm $H_2O$ | 0.932666898 | 0.982886955 |

| Gene | Protein ID | logFC | AveExpr | comparison | pvalue | fdr |
|---|---|---|---|---|---|---|
| ATP5F1B | P06576 | -0.003502911 | 21.08547363 | 7 m$H_2O$ vs 3 mm $H_2O$ | 0.935054579 | 0.982886955 |
| MOGS | Q13724 | 0.00521333 | 16.31339399 | 7 m$H_2O$ vs 3 mm $H_2O$ | 0.935661155 | 0.982886955 |
| ATP2A2 | P16615 | -0.003279664 | 19.13691058 | 7 m$H_2O$ vs 3 mm $H_2O$ | 0.943704701 | 0.988219074 |
| TGFBI | Q15582 | 0.002804034 | 22.50578879 | 7 m$H_2O$ vs 3 mm $H_2O$ | 0.95684905 | 0.995314595 |
| NAMPT | P43490 | 0.005784073 | 15.42557579 | 7 m$H_2O$ vs 3 mm $H_2O$ | 0.959067337 | 0.995314595 |
| H1-5 | P16401 | -0.011309981 | 22.04840624 | 7 m$H_2O$ vs 3 mm $H_2O$ | 0.959739727 | 0.995314595 |
| LMNB2 | Q03252 | -0.002878105 | 20.43563345 | 7 m$H_2O$ vs 3 mm $H_2O$ | 0.965890083 | 0.995314595 |
| KRT10 | P13645 | 0.009234885 | 21.96807875 | 7 m$H_2O$ vs 3 mm $H_2O$ | 0.967055281 | 0.995314595 |
| GNAI2 | P04899 | -0.002107021 | 19.49499447 | 7 m$H_2O$ vs 3 mm $H_2O$ | 0.968414201 | 0.995314595 |
| NID1 | P14543 | 0.001634621 | 20.1895549 | 7 m$H_2O$ vs 3 mm $H_2O$ | 0.97568355 | 0.995889356 |
| RSL1D1 | O76021 | 0.002406067 | 14.99516928 | 7 m$H_2O$ vs 3 mm $H_2O$ | 0.97912034 | 0.995889356 |
| H4C16 | P62805 | 0.00543323 | 23.19715251 | 7 m$H_2O$ vs 3 mm $H_2O$ | 0.980514462 | 0.995889356 |
| HADHB | P55084 | 0.000653127 | 19.70348331 | 7 m$H_2O$ vs 3 mm $H_2O$ | 0.989501387 | 0.995889356 |
| PFN1 | P07737 | 0.000802 | 17.42183319 | 7 m$H_2O$ vs 3 mm $H_2O$ | 0.991760316 | 0.995889356 |
| LMNB1 | P20700 | -0.000658144 | 20.31075579 | 7 m$H_2O$ vs 3 mm $H_2O$ | 0.992789145 | 0.995889356 |
| DDX5 | P17844 | 0.000433114 | 18.93492511 | 7 m$H_2O$ vs 3 mm $H_2O$ | 0.992806767 | 0.995889356 |
| VAPA | Q9P0L0 | 0.001478908 | 16.66742819 | 7 m$H_2O$ vs 3 mm $H_2O$ | 0.992898698 | 0.995889356 |
| PDLIM5 | Q96HC4 | -6.50E-05 | 17.8332416 | 7 m$H_2O$ vs 3 mm $H_2O$ | 0.999570647 | 0.999570647 |
